# Rapid simulation of organ-scale oxygen transport for tissue ischemia prediction

**DOI:** 10.64898/2026.10.08.757708

**Authors:** Carl Anthony Kohnke, Yee Lin Tan, Ian A. Coates, Zachary A. Sexton, Alison L. Marsden, Joseph M. DeSimone, Eric S. G. Shaqfeh

## Abstract

A densely vascularized tissue can still be starved of oxygen. Tissue oxygenation depends heavily on vessel arrangement, blood flow distributions, and cellular metabolism. Resolving these effects from individual constituent vessels to whole organs has historically been computationally prohibitive. We introduce the Cardiovascular Analysis and Simulation of Circulatory Advection, Diffusion, and Exchange (CASCADE) toolkit, which makes vessel-resolved oxygen transport tractable at the organ scale through a novel screened Green’s-function formulation. On laptop-class hardware, CASCADE simulates networks containing thousands of vessels in under one second and a full biventricular human heart model in about two minutes. We validate spatial oxygen and cell-survival predictions against experiments in perfused engineered tissues and matched finite-element simulations. In engineered tissues, sweeps across thousands of network designs and physiological conditions reveal when vessel density, oxygen supply, or network architecture limits viability, establishing design rules for sustaining thick constructs. In a human heart model, CASCADE translates coronary occlusion, anemia, polycythemia, and hypoxemia into distinct myocardial oxygenation patterns, revealing where oxygen delivery fails and how much tissue is at risk.

## 1 Introduction

The capacity of a vascular network to sustain living tissue emerges from the collective behavior of its vessels, where even small differences in vessel organization and blood flow can reshape perfusion across large regions of tissue. Oxygen delivery is perhaps the most unforgiving aspect of this process, as oxygen has a low solubility in tissue, is poorly stored by cells, and is rapidly consumed, restricting the analyte to a diffusive reach on the order of only ∼100 *µ*m from the nearest vessel ^1,2^.

The challenge of providing adequate oxygenation spans both engineered and native tissues. In engineered constructs, insufficient oxygen delivery limits tissue thickness and achievable cellular density, imposing vascular architecture and perfusion as central design variables ^3–5^. With generative vascular design and biofabrication techniques reaching organ scales ^3,5,6^, the ability to inscribe vascular networks within tissues is beginning to outpace our ability to predict their performance. In native organs, stenosis, occlusion, hypoxemia, and changes in hematocrit disrupt oxygen delivery within a mostly fixed anatomy, producing mercurial and spatially heterogeneous ischemia ^7^. Understanding ischemia is especially consequential in cardiovascular health, as ischemic heart disease remains the leading cause of death worldwide ^8,9^. Despite its importance, oxygen transport is comparatively underrepresented in emerging cardiovascular digital twins, which more commonly resolve anatomy, electrophysiology, hemodynamics, and solid mechanics ^10–12^.

The central challenge is computational. Predicting oxygenation requires coupling network hemodynamics, convective oxygen transport, oxyhemoglobin binding, individual-vessel exchange, diffusion through tissue, and cellular metabolism across scales from micrometer-wide vessels to centimeter-scale tissues. Since Krogh introduced his idealized capillary cylinder model ^13^ over a century ago, researchers have approached this multiscale problem with various mesh- and voxel-based, mixed-dimensional, and Green’s-function approaches ^14–20^.

Yet a fundamental tradeoff persists: lumped-parameter or homogenized models can describe large domains but lose important microscale information, whereas microvessel-resolved approaches retain such physics at a computational cost that rises rapidly with tissue volume and resolution. A model scalable enough to simulate large tissues must therefore retain individual-vessel transport without forcing the surrounding tissue to be resolved at the same spatial resolution. Classical Green’s-function methods (GFMs) attempt this, avoiding body-fitted meshes by representing vessels as oxygen sources and tissue as a dense grid of unknown-strength sinks and superimposing analytical expressions to find the oxygen field^18–20^. However, the resulting source-sink system remains globally coupled, confining its use still to small microvascular territories.

Even for these small territories, most existing solvers require minutes to hours per simulation. This computational barrier largely confines vessel-resolved perfusion investigations to isolated case studies, preventing methodical interrogation of vascular design and physiology, let alone tracing consequences across entire organs. To make such analyses tractable, we introduce the Cardiovascular Analysis and Simulation of Circulatory Advection, Diffusion, and Exchange (CAS-CADE) toolkit, a vessel-resolved framework built around a new Yukawa-Green’s-function method (YGFM) for oxygen transport. By embedding cellular oxygen consumption in exponentially screened Green’s-functions, YGFM eliminates the cost of explicit tissue discretization and localizes the influence of each vessel. Coupled to network hemodynamics, phase separation modeling, and an intravascular transport solver, CASCADE resolves networks containing thousands of discrete vessels in fractions of a second and systems with tens of millions of vessels in minutes on laptop-class hardware. We validate CASCADE against engineered-tissue oxygen measurements, cell-survival patterns, and matched finite-element method (FEM) simulations.

We use CASCADE’s novel throughput to expose the organizing principles of vascular oxygenation across scales. In centimeter-scale tissues, hundreds of thousands of variations in vascular architecture, blood and tissue properties, and metabolic demand collapse into distinct transport regimes that define the perfusion conditions and vessel organization required to sustain living tissue. Extending the same framework to a human biventricular heart model, we trace how focal coronary lesions and systemic physiological changes propagate through the vascular hierarchy, reshaping myocardial oxygenation across the organ. By extending the tractable frontier of vessel-resolved perfusion calculations, CASCADE brings previously inaccessible questions of multiscale vascular function within reach of direct mechanistic exploration.

## 2 Results

### 2.1 A scalable framework for vessel-resolved oxygen transport

CASCADE targets the computational cost of computing three-dimensional oxygen fields surrounding discrete vascular networks. Hemodynamics and intravascular transport can be solved on a vascular graph, but classical vessel-resolved methods also finely discretize the surrounding tissue to capture diffusion and consumption. As networks grow, these tissue unknowns become a major computational burden. Our approach removes this necessity.

We incorporate cellular oxygen consumption directly into the tissue transport operator through local linearization of cellular uptake about an effective oxygen concentration *C*^eff^, giving

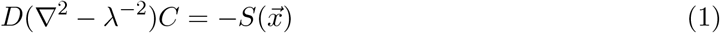

where *C* is oxygen concentration, *D* is tissue diffusivity, 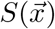 is the vascular point-source distribution and 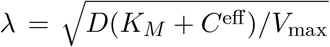 is the local reaction-diffusion length scale, with *V*_max_ and *K*_*M*_ being Michaelis-Menten reaction constants. The corresponding Green’s-function solution is the Yukawa kernel,

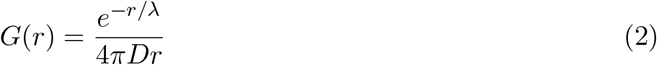

which simultaneously describes oxygen diffusion away from a vessel and its approximate consumption by the surrounding tissue. This embedding is consequential for two reasons. First, this change eliminates the field of tissue sinks necessary for classical GFMs (Fig. 1f), as the Yukawa kernel already encodes their approximate effect. Second, the known exponential decay of the Yukawa response allows sufficiently distant (∼6*λ*) source contributions to be rejected prior to evaluation. These two factors are central to the efficiency of the Yukawa-Green’s-function method (YGFM) which forms the computational core of CASCADE (Fig. 1g).

**Figure 1.**
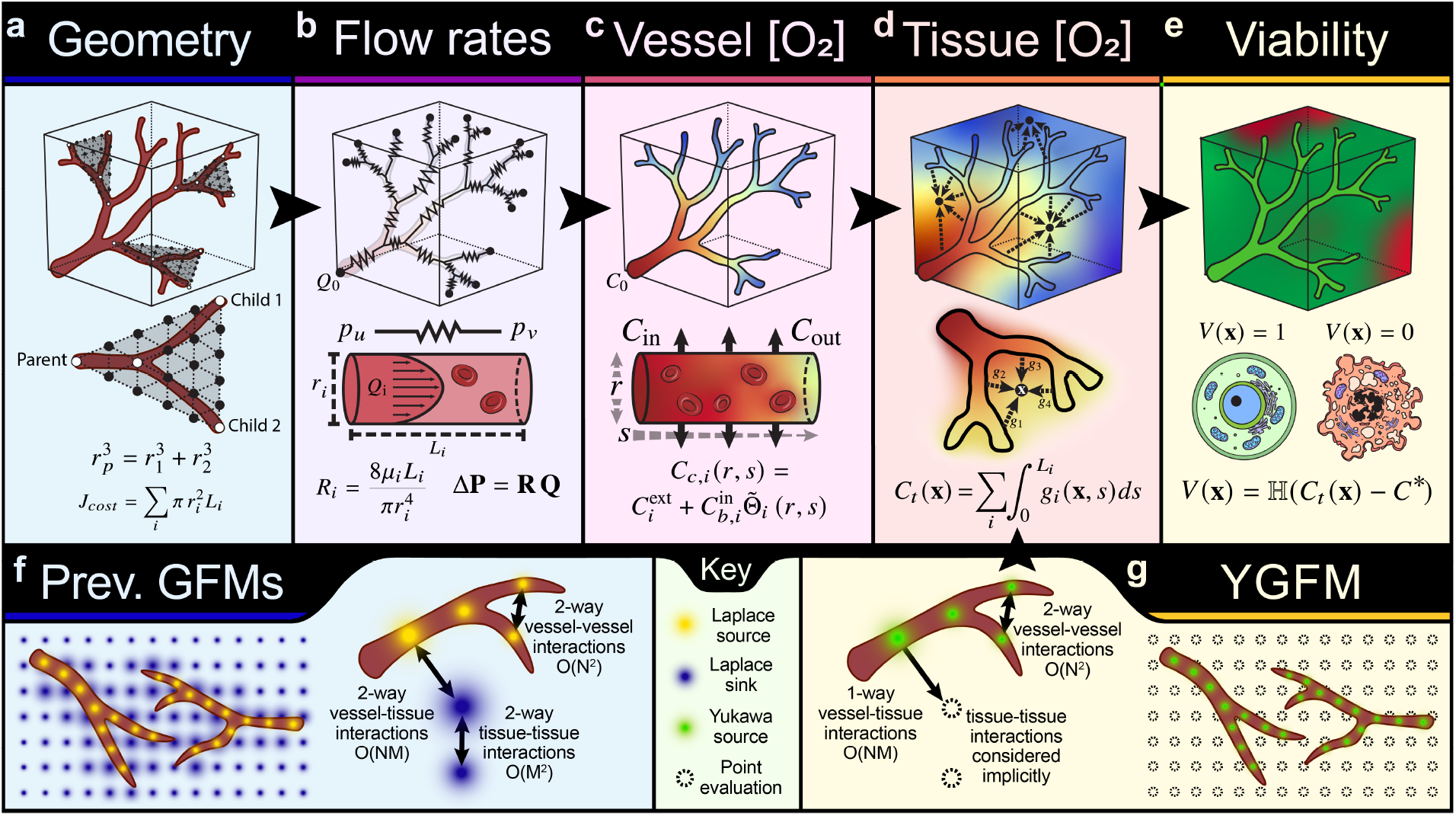
Overview of the CASCADE computational framework. a–e, A schematic of the CASCADE workflow. From left to right: a, Generate or upload a segmented vessel architecture. Shown is a schematic of Constrained Constructive Optimization (CCO), a generative design method which minimizes a cost function while optimizing for low flow resistance and agreement with Murray’s law of vessel bifurcations; b, Compute flow rates and pressures within the network using a reduced-order network flow solver; c, Determine vessel oxygen propagation profiles down the network, considering oxygen depletion, re-uptake from other vessels, and various transport resistances; d, Estimate the oxygenation of cellular tissues surrounding the vascular architecture using the Yukawa-Green’s-function method (YGFM); e, Predict tissue viability distributions by comparing tissue oxygenation to an oxygen-concentration threshold for viability or functionality. f, Visualization of previous Green’s-function methods ^18,20^ showing that two-way pairwise interactions must be computed for vessel-vessel, vessel-tissue, and tissue-tissue interactions. g, Visualization of the Yukawa-Green’s-function method (YGFM) developed in this work, where only vessel-vessel interactions require two-way coupling, vessel-tissue interactions are one-way, and tissue-tissue computations are eliminated.

Vessels remain coupled to the tissue they supply through a linear driving force relationship (Eqn 15). Once source strengths and wall concentrations are known, the 3D tissue oxygen field is obtained by superimposing the screened contribution from each vessel (Fig. 1c,d). Integrating moments along vessel lengths and summing all vessel contributions yields

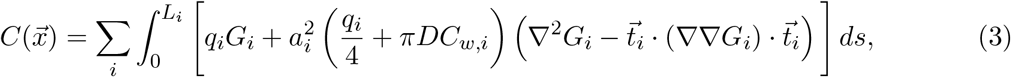

where *L*_*i*_, *a*_*i*_, and 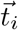 are the length, radius, and local orientation of vessel *i*, and 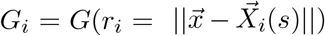 where 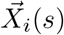 is the position of the contributing vessel’s centerline, along the vessel’s axial coordinate *s*. The leading term treats each vessel as a flux-emitting line source while the second captures finite-radius and orientation-dependent near-field effects. Evaluating a corresponding expression along vessel centerlines gives the external field *C*_ext_ felt by each vessel, closing the interactions between neighboring vessels.

The remaining components of CASCADE provide the vessel-wall concentrations required by Eqn. 3. Hemodynamics and intravessel oxygen transport are solved on a reduced-order graph (Fig. 1a–c), with further hydraulic, phase-separation, intraluminal oxygen gradient, vessel-vessel exchange, and vessel-tissue exchange solvers as described in the Methods and Supplementary Information.

Furthermore, if a critical oxygen concentration (*C*^*\**^) for survival or functionality can be estimated, the resulting oxygen field can be mapped to a tissue state prediction such as Eqn. 4^5,21,22^ (Fig. 1e).

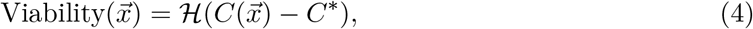

where *ℋ* is the Heaviside step function. Together, these formulations allow CASCADE to retain individual-vessel oxygen transport and three-dimensional concentration gradients without explicit discretization of tissue metabolism during the solve.

### 2.2 Validation of oxygenation and tissue viability predictions

We first assessed whether this reduction preserves the spatial oxygen gradients and biological outcomes it was designed to predict. Straight and serpentine channels were formed in HEK-293T cell-laden gelatin methacrylate (GelMA) constructs using photopatterned-Channel Architectures with Sacrificial Templates (pCAST) ^5^ (Fig. 2a). Oxygen-sensitive foils provided spatial dissolved-oxygen measurements (Fig. 2b), while matched 3D FEM simulations supplied an independent numerical solution under functionally identical conditions.

**Figure 2.**
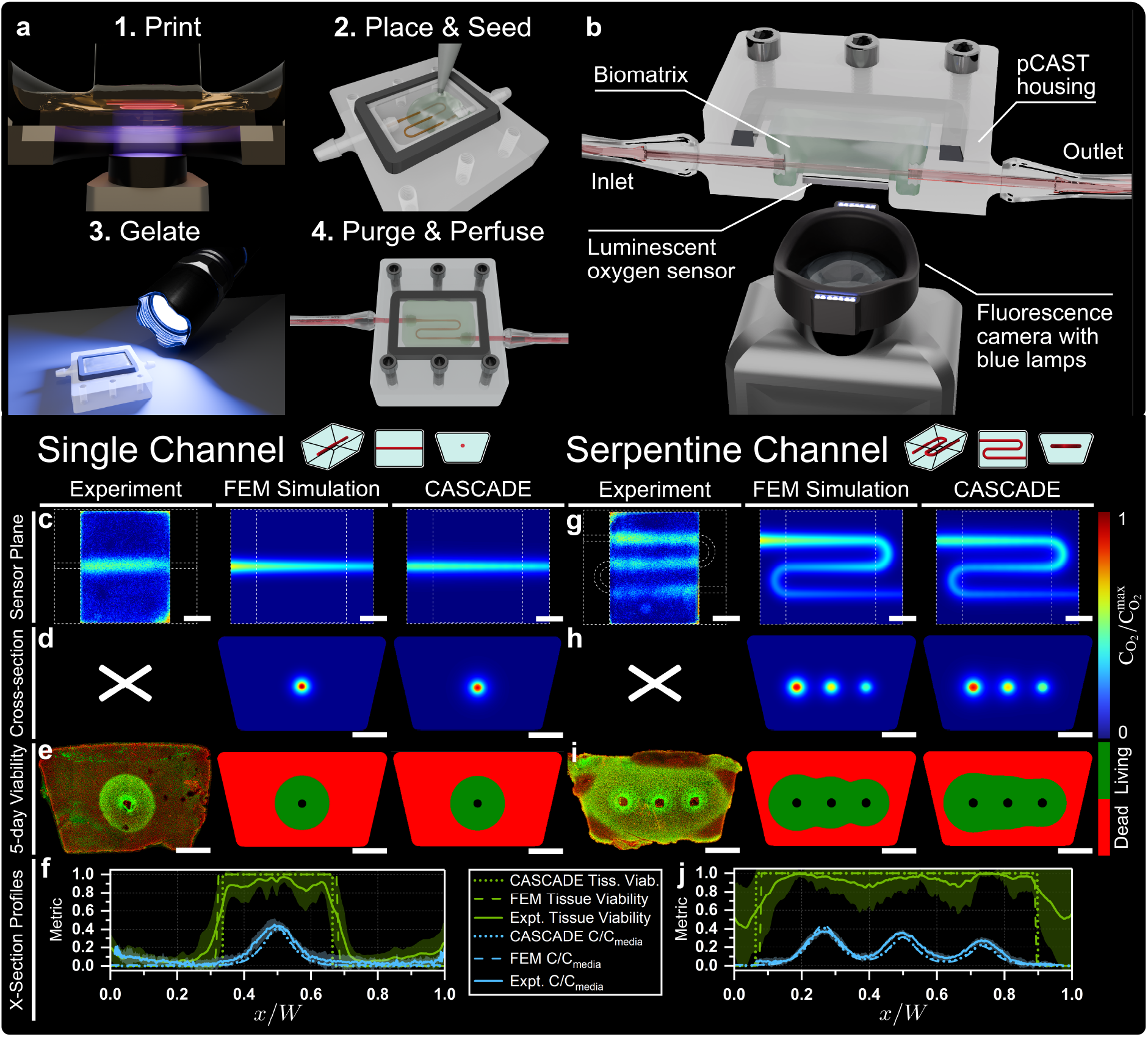
Fabrication and experimental setup diagrams; oxygen and tissue viability comparisons to experiments and numerical simulations. a, A schematic of the pCAST method for sacrificial templating of channel architectures ^5^, showing (1) CLIP 3D printing of sacrificial structures, (2) placing printed structures within a casting chamber and seeding with cellularized GelMA, (3) photocuring of the GelMA matrix, and (4) purging of the sacrificial material and perfusion with cell culture medium. b, Diagram showing a perfused pCAST housing with an embedded oxygen sensor and fluorescence camera for imaging. c, Experimental oxygen-sensor measurements (N=4) and corresponding FEM and CASCADE predictions at the matched sensor plane. d, FEM and CASCADE oxygen fields in a central cross-section through the channel. e, A representative experimental live/dead confocal image of a single-channel construct (N=6) and corresponding FEM and CASCADE oxygen-thresholded viability predictions in a cross-section. f, Oxygen concentration (blue) and tissue viability (green) across the construct width, comparing experimental measurements, FEM simulations, and CASCADE predictions. Oxygen profiles were obtained by averaging measurements across a narrow strip through the center of the sensor face, perpendicular to channel flow. Viability profiles were collected from the confocal cross-sectional images along a line spanning the construct approximately 100 µm below the bottom of the channel. Position and concentration are normalized by construct width and inlet concentration, respectively. Viability is a normalized fluorescence-based survival index. Experimental curves show µ *±* σ with error bands excluding sensor noise. g–j, Corresponding measurements and predictions for the serpentine geometry (N=3 oxygen measurements and N=5 live/dead images). All scale bars are 2 mm.

The two geometries probed complementary transport behavior. The straight channel isolated radial diffusion-reaction competition in the metabolically active tissue. CASCADE reproduced the measured oxygen profile and closely matched the simulated field produced by FEM (Fig. 2c,d,f). The serpentine geometry tested interactions among neighboring oxygen fields while exhibiting axial depletion along the extended flow path. CASCADE captured the resulting asymmetry and progressive weakening of oxygen peaks (Fig. 2g,h,j).

We next compared oxygen-limited survival with live/dead confocal imaging after five days of perfusion. To do this, we applied a prespecified critical concentration of 1.4 mmol/m^3^ (*≈*1 mmHg), below which oxidative metabolism becomes severely impaired and sustained exposure can compromise cell viability^23–25^. CASCADE replicated the transition from viable cells near perfused channels to nonviable tissue farther away, with agreement being strongest for the straight channels and remaining qualitatively consistent in the more variable serpentine constructs (Fig. 2e,f,i,j).

Because these experiments necessarily sampled simple geometries, we extended validation to 25 higher-complexity computational studies, in which we compared CASCADE to FEM simulations spanning domain sizes, biomimetic architectures, perfusion rates and metabolic conditions. Across 233,140 point-wise oxygen-measurement comparisons, CASCADE achieved a concordance correlation coefficient of 0.955 relative to FEM solutions. Applying matching viability thresholds yielded 96.8% classification agreement, with false-viable and false-nonviable rates of 2.0% and 4.5%, respectively, and no substantial case-level error outliers (Supplementary Fig. S10 and Tab. S2). CASCADE also outperformed an optimized Krogh-cylinder benchmark model in concentration concordance and viability-classification agreement (Supplementary Figs. S18–S19, Tabs. S4–S5).

### 2.3 Characterizing computational efficiency

In changing the representation of the tissue metabolism, Yukawa screening also changes the computational structure of the problem. Conventional GFMs require global coupling of oxygen-source (vessels) and oxygen-sink (tissue) quadrature nodes, causing the solution complexity to scale as *O*((*N* +*M*)^2^) for *N* source quadrature nodes and *M* distributed tissue sinks. Elimination of tissue sinks and restriction of pairwise interactions to local neighborhoods greatly reduce the dominant workload, causing the problem to instead scale as *O*(*K*_*v*_*N* + *K*_*t*_*M*) with *K*_*v*_ and *K*_*t*_ representing an average number of neighborhood-retained sources for vessel and tissue evaluations, respectively. Note that in the screened formulation, *M* now only refers to the number of tissue measurements, as there are no tissue quadrature nodes. CASCADE further accelerates these operations using GPU parallelization, k-d tree spatial indexing, precomputed intraluminal gradients, and a Fourier-space implementation for densely vascularized systems.

We performed simulations on 36 computationally generated networks spanning 1 to 10 million vessel segments in 1 cm^3^ of virtual myocardium, showing that the complete workflow requires 0.08–1 seconds for networks up to 400,000 segments and about 23 seconds for 10 million segments (Fig. 3a,b).

**Figure 3.**
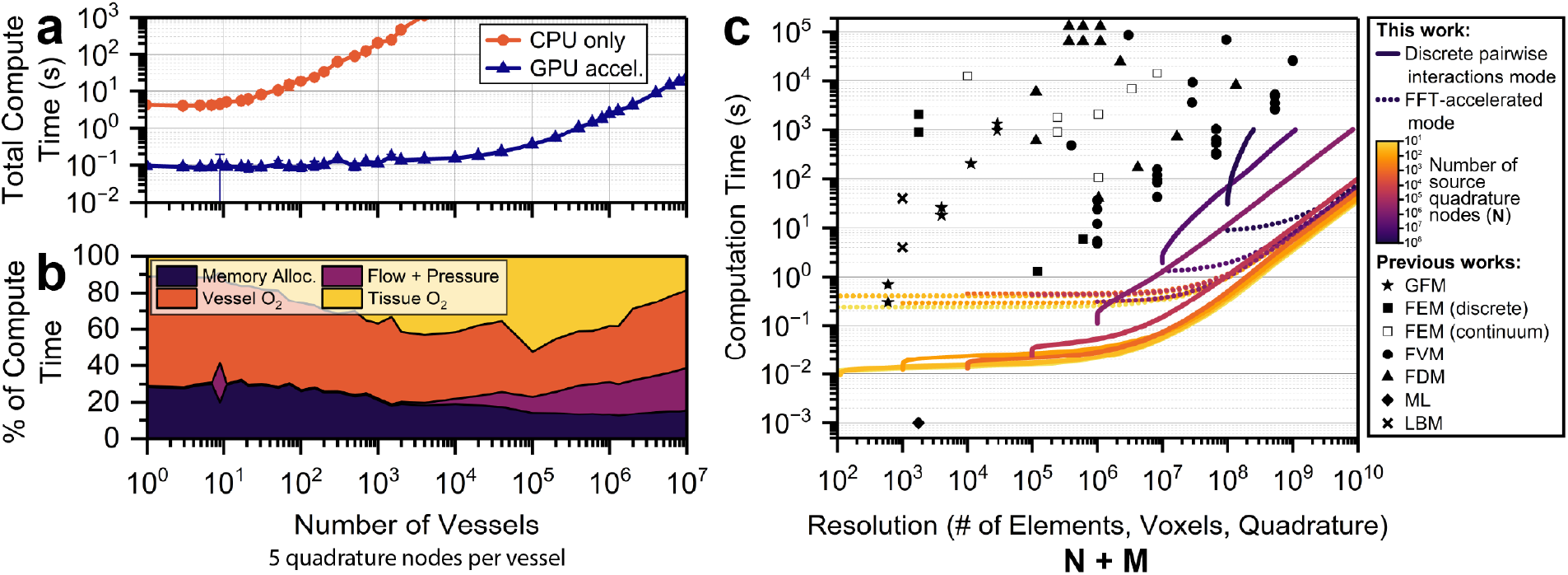
Computational advantage characterizations. a, Total compute time versus vascular complexity for the entire CASCADE simulation workflow (excluding vessel architecture generation) with and without GPU acceleration. Five Gauss-Legendre quadrature nodes were used per vessel segment. b, Percent breakdown of computational load as complexity increases with GPU acceleration active. c, Computation time versus simulation resolution (N +M). For this work, solid lines represent computations performed with all pairwise interactions within a distance *≤*6λ, dotted lines indicate simulations performed in Fourier space on a Cartesian grid, the line color indicates the complexity of the vascular network according to the number of source quadrature nodes N, and M refers to the number of tissue evaluation locations. Scatter-plotted shapes represent literature-reported computation times for Green’s-function methods ^20,26^, Finite-Element Methods (FEM) ^27–30^, Finite-Volume Methods (FVM) ^15,31–33^, Finite-Difference Methods (FDM) ^16,34,35^, machine learning (ML) methods ^30^, and Lattice Boltzmann methods (LBM) ^36^. For the literature-reported methods, resolution refers to each method’s reported discretization.

We then conducted more than 7,800 additional timing runs that varied the number of source nodes *N* and number of tissue oxygen field point evaluations *M*, denoting our method’s resolution as *N* + *M* and comparing to the discretization-speed tradeoff of existing methods. Fig. 3c shows that across a range of spatial resolutions, both our discrete pairwise-interaction and fast Fourier transform (FFT) implementations were up to 2–3 orders of magnitude faster than published runtimes of conventional GFMs^20,26^, and 1–5 orders of magnitude faster than existing finite-volume approaches ^15,31–33^. The cross-study comparisons are approximate because hardware, domain size, and implementation differ. We propose that with CASCADE’s efficiency, thousands of vascular conditions can be explored as parameter sweeps rather than isolated case studies.

### 2.4 Defining design regimes in thick, engineered tissues

We next applied CASCADE to vascular tissues at scales where methodical experimental exploration becomes prohibitive. Large vascular constructs can require hours to days to fabricate and days to weeks for culturing, perfusion, and analysis. Meanwhile, increased 3D complexity introduces additional constraints on achievable feature size, materials, fabrication methods, and handling ^37^. Consequently, even basic design questions (how much vasculature is required, how complex must it be, or which vascular geometries best meet particular transport demands) remain difficult to resolve.

We began by varying the vascular density of branched networks supplying 1 cm^3^ of myocardium perfused with blood or cell-culture medium at 900 *µ*L/min (Fig. 4a–d).Sparse networks sustained only narrow perivascular regions, while increasing vascular density caused merging of neighboring oxygen fields into interconnected viable tissue. Networks containing ∼10^6^ segments supported more than 95% of the volume. O_2_-limited viability approached unity once the average cell’s distance to the nearest channel wall (DNCW) reached roughly 20–40 *µ*m, comparable to the tens-of-micrometers spacing of capillaries in native human myocardium^38,39^. Increasing network density also reduced hydraulic resistance per unit vessel length, simultaneously improving tissue access and flow distribution efficiency.

**Figure 4.**
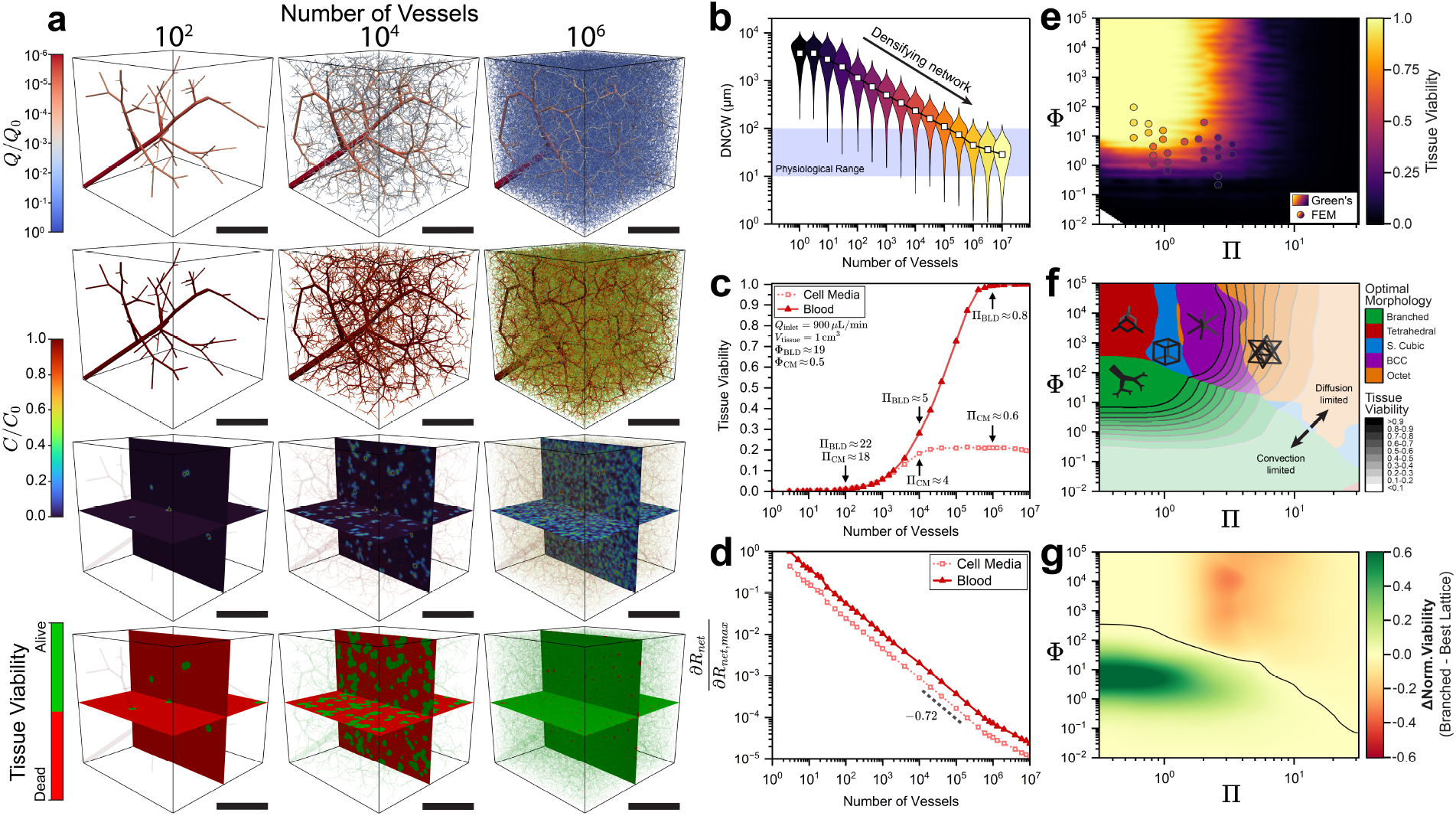
Oxygen dynamics of thick vascular tissues and architectural design rules. a, 3D depiction of simulation results for a 1 cm^3^ cube of myocardial tissue with an increasing number of vessels from left to right. Shown are volumetric flow rate (top row), bulk vessel O_2_ concentration (upper middle row), tissue O_2_ concentration (lower middle row), and predicted tissue viability distributions (bottom row). Cube side lengths are all 1 cm. Scale bars are all 5 mm. b, Violin plots of distance to nearest channel wall (DNCW) distributions from within the tissue as a function of vessel count. At each vessel count, violin width indicates the relative frequency of tissue measurements taken at a given distance. c, Mean tissue viability as a function of vessel count for blood and cell culture medium as perfusates. d, Normalized net resistance per unit length as a function of vessel count for blood and cell culture medium. e, Heatmap of mean tissue viability in relation to an oxygen supply-vs-demand parameter Φ and a relative channel spacing parameter Π. f, Best-performing vessel morphology across the oxygen-supply and vessel-spacing regimes. At each (Π, Φ) pairing, the color identifies the architecture achieving the largest viable tissue fraction after accounting for the volume occupied by the channels. Contour lines and color opacity indicate the achieved fraction. g, Difference in this volume-adjusted viable fraction between branched networks and the best-performing lattice at each (Π, Φ) coordinate. Positive values favor branched networks while negative values favor lattices. The black line marks the equal performance boundary.

Simulations of blood versus culture medium perfusion revealed substantial differences in oxygenation under otherwise identical conditions. Blood could ultimately support the entire tissue volume, but culture medium remained strongly supply-limited despite equivalent vascular density. Without hemoglobin buffering the reduction of oxygen content along channel lengths, oxygen was consumed before reaching distal regions of the tissue. Yet, supply alone was also insufficient; even if abundant oxygen entered a network, substantial tissue volume could remain beyond its diffusive reach. To distinguish these physical failure modes, we defined two primary governing parameters.

The first was a supply-to-demand parameter

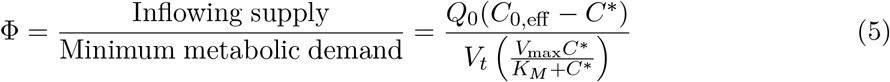

which compares inflowing oxygen above the viability threshold with average tissue demand evaluated at that threshold. *Q*_0_ is the inlet blood flow rate, *V*_*t*_ is the volume of the tissue being perfused, *C*_0,eff_ = *C*_0_ + *C*_0,*Hb*_ accounts for both dissolved and hemoglobin-bound inlet oxygen, and *V*_max_ and *K*_*M*_ are the Michaelis-Menten parameters.

The second was a ratio of length scales

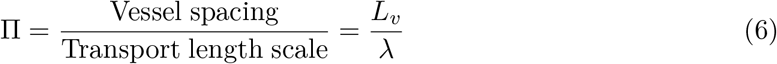

which compares the mean vessel-to-vessel spacing with the characteristic reaction-diffusion distance *λ* (using *C*^eff^ = *C*_0_).

We mapped 8,560 simulations across this Π *™* Φ space and independently overlaid the 25 FEM validation cases. Despite broad variation in geometry, vascular density, metabolism, diffusivity, perfusion, oxygen-carrying capacity, and viability threshold, the simulations collapsed onto a common viability landscape (Fig. 4e). Near-complete viability generally required both adequate incoming supply (Φ ≳ 10) and vessel spacing within about 1.5 reaction-diffusion lengths (Π ≲ 1.5). Neither condition alone was sufficient. Increasing supply could not compensate for poor spatial access, while tighter spacing produced diminishing benefit if incoming oxygen was insufficient to meet demand. The earlier myocardial tissue simulations followed the map, as blood remained supply-sufficient while increasing vascular density drove viability above 99%, yet culture medium remained near Φ *≈* 0.5 and only reached ∼20%.

These regimes raised a further design question: is a hierarchical, branched network the optimal morphology under either transport constraint? Regular lattice channel networks are readily manufactured by established patterning and sacrificial-templating approaches, whereas reproducing the multiscale hierarchy and topology of native vasculature remains substantially more involved ^3^. To assess these design types, we compared biomimetic branched trees with tetrahedral, simple-cubic, body-centered cubic, and octet lattice networks across matched physical conditions, comparing them by viable tissue fraction after accounting for vascular volume. In total, 115,248 simulations were performed (Fig. 4f,g).

Architecture was most consequential near apparent physical transport limits. At low Φ and low Π, where oxygen supply was limiting, branched networks outperformed lattices by up to 60% in volume-normalized viability. Their larger proximal conduits preserved efficient long-range delivery before progressively smaller vessels released oxygen distally. The opposite behavior emerged at high Φ and high Π, where oxygen was abundant but widely spaced vessels limited diffusive coverage. Under high-supply conditions, the regular, redundant lattice pathways reduced the occurrence of interspersed under-supplied regions. In this regime, the best lattices exceeded branched network performance by up to ∼30%, with higher node-edge connectivity becoming more favorable with increasing spacing.

Topology was less consequential elsewhere. When oxygen was plentiful and readily accessible, any architecture was sufficient. When neither requirement was satisfied, geometry alone could not rescue the tissue. The consequence of network architecture was therefore most significant near diffusion-limited and convection-limited transport boundaries, where hydraulic efficiency and homogeneous delivery were the determining factors.

### 2.5 Simulating myocardial responses to vascular and systemic perturbations

The preceding analyses treated vascular architecture as a design variable. Native organs instead impose a largely fixed anatomy within which disease and physiology perturb transport. We therefore tested CASCADE in a 141 cm^3^ biventricular myocardial model containing segmented proximal coronary arteries and computationally generated distal vasculature (Fig. 5). Each whole-heart simulation required about 2 minutes on a personal laptop.

**Figure 5.**
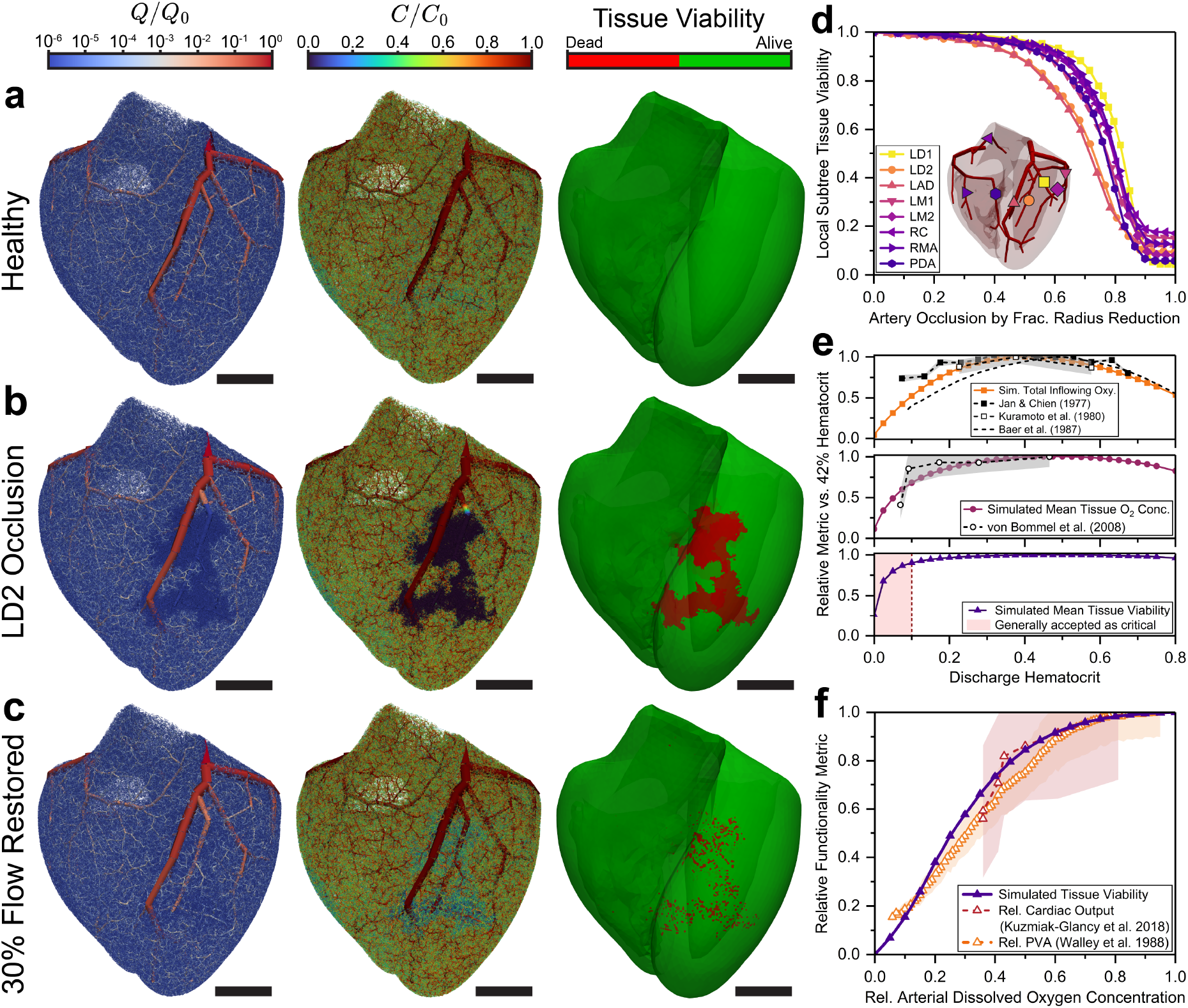
Biventricular heart myocardial oxygenation under coronary occlusion, altered hematocrit, and hypoxemia. a, Biventricular heart CASCADE simulation of healthy ventricles showing, from left to right, vessel flow rate normalized by total inlet flow (Q/Q_0_), intravascular dissolved oxygen concentration normalized by inlet concentration (C/C_0_), and predicted myocardial tissue viability. Green and red indicate tissue above and below the prescribed oxygen threshold, respectively. The largest 25 million vessels are visualized. b, Corresponding distributions after complete occlusion of the second left diagonal (LD2) coronary branch. c, Distributions after restoring LD2 flow to 30% of its baseline value. All scale bars in a–c are 2 cm. d, Predicted viable fraction of the myocardial territory downstream of each coronary branch as its radius is progressively reduced. A fractional radius reduction of 0 indicates no narrowing and 1 indicates complete occlusion. The inset identifies the independently perturbed branches: the first and second left diagonal branches (LD1, LD2), left anterior descending (LAD), first and second left marginal branches (LM1, LM2), right conus (RC), right marginal artery (RMA), and posterior descending artery (PDA). e, Effects of arterial discharge hematocrit at a fixed coronary pressure drop. Top, total oxygen inflow compared with coronary oxygen-transport measurements in dogs ^40–42^. Middle, mean myocardial oxygen concentration compared with cardiac microvascular oxygenation measurements in rats ^43^. Bottom, predicted viable myocardial fraction. Oxygen inflow and oxygenation are shown relative to their values at 42% hematocrit. The shaded low-hematocrit region marks severe anemia associated with loss of cardiovascular compensation ^44^. f, Predicted viable myocardial tissue fraction as arterial oxygenation decreases (progressive hypoxemia), shown alongside normalized cardiac output from isolated working rabbit hearts ^45^ and left-ventricular pressure-volume area from dogs ^46^. These experimental measures provide physiological comparisons with the simulated viability responses. Experimental measures of cardiac function are normalized to their respective maxima; shaded bands in e and f show reported experimental variability.

Under baseline conditions, intravascular oxygen partial pressure declined to 20–30 mmHg near terminal paths, consistent with reported coronary venous oxygenation and the high resting extraction of myocardium^39^, while delivery remained sufficient to maintain the entire myocardial domain above the non-ischemic viability threshold.

Complete occlusion of the second left diagonal (LD2) branch extinguished downstream flow and formed a contiguous region of severe hypoxia extending transmurally from epicardium to endocardium (Fig. 5b), although adjacent coronary territories preserved tissue near the margins. Restoring 30% of baseline LD2 flow rescued most of the affected regions but left scattered ischemic foci as distal paths became depleted before reaching their terminal ends (Fig. 5c).

Progressive stenosis produced similarly nonlinear, anatomy-dependent effects (Fig. 5d). LD2 and left anterior descending (LAD) territories were most sensitive, with ∼50% of tissue remaining above the oxygen threshold at 72% radius reduction, while the first left diagonal (LD1) territory reached the same point near 82%. The predicted transition occurred within the same range reported in classical canine experiments, which identified a diameter reduction of 74 *±* 2% as critical in the left circumflex artery^47^. Branch-to-branch variation further illustrates that stenosis severity alone does not uniquely define consequences in the tissue.

We next examined the effects of anemia and polycythemia by varying hematocrit from 0 to 80% at a fixed coronary pressure drop (Fig. 5e). Myocardial oxygen inflow peaked broadly near the physiological range, falling at both extremes as oxygen-carrying capacity competed with blood viscosity. Oxygenation and viability remained comparatively stable through moderate hemodilution, mostly declining below about 20% hematocrit and falling steeply near 10%, consistent with independent measurements of coronary oxygen transport and microvascular oxygenation ^40–44^. By contrast, polycythemia reduced transport reserve without materially compromising basal tissue viability.

Finally, reducing arterial dissolved oxygen content produced little change through moderate hypoxemia, followed by a nonlinear decline at lower oxygen content (Fig. 5f). Mean viability crossed 50% at ∼26% of normoxic arterial oxygenation, and the simulated viability curve closely followed previously reported deterioration of cardiac function during progressive hypoxemia in animal hearts ^45,46^.

## 3 Discussion

Resolving vascular oxygen transport has conventionally required a compromise between microvascular detail and tractable scale. CASCADE relaxes this compromise by retaining individual vessels while avoiding explicit volumetric resolution of the surrounding tissue. The significance lies not only in the reduced computational cost but also in the analyses that such speed makes practical. Dense vascular networks, centimeter-scale tissues, whole organs, and high-dimensional parameter spaces can be examined within a common mechanistic framework, as simulations requiring only seconds expose relationships that would remain hidden in isolated calculations.

The parameters Π and Φ separate the limits of adequate oxygenation, showing that vessel density cannot overcome inadequate supply, while increased inflowing oxygen content provides diminishing benefits when vessels are too widely spaced. The Π–Φ landscape may therefore guide which transport constraint should be addressed, potentially mitigating unnecessary experimental iteration during future engineered tissue design.

Vascular architecture further partitions these regimes. Hierarchical trees favor long-range convective delivery, but lattices distribute exchange surfaces more uniformly. Neither is universally optimal, as their relative advantages depend on the active transport bottleneck. Native hierarchy therefore need not be an objective in itself when simpler architectures satisfy the relevant constraints (although lattices may impose significantly higher hydraulic costs^6^). The division between hierarchical delivery and mesh-like local exchange reflects native angiosome organization ^3,48^, though natural morphologies also consider other factors such as redundancy against occlusions^49,50^ and caliber regulation^51^.

At the organ scale, patient-specific computational fluid dynamics models have used CT angiography to estimate coronary fractional flow reserve (FFR)^52,53^, while multiscale models have coupled three-dimensional coronary flow to zero-dimensional synthetic distal trees and myocardial perfusion^54^. Our whole-heart calculations demonstrate a new capability to directly examine the downstream oxygen-delivery consequences of vascular perturbations, showing how regional coronary lesions and systemic changes can produce different myocardial oxygen deficits and spatial patterns of tissue at risk. Vessel-scale oxygen transport may therefore add a complementary functional connection between vascular state and organ-scale ischemia in future digital twins.

Several limitations define the interpretation and next steps of our framework. CASCADE presently describes steady or quasisteady oxygen transport for prescribed vascular, hemodynamic, and metabolic states. Predicted nonviability therefore represents tissue exposed to sustained oxygen limitation rather than a direct prediction of irreversible infarction or death. A mechanistic injury model may require additional chemical dynamics, autoregulation, collateral recruitment, changing myocardial work, thrombosis, reperfusion injury, and other time-dependent processes. The heart demonstration uses computationally generated distal vasculature for one myocardial anatomy, so branch-specific responses should not be interpreted as clinical thresholds. The locally linearized metabolic formulation also becomes less accurate in hyperoxic conditions as Michaelis-Menten consumption approaches zeroth-order behavior, and the infinite-domain Green’s-function assumption weakens when the characteristic reaction-diffusion length approaches tissue dimensions.

More broadly, CASCADE opens a new computational regime for studying vascular oxygen transport in which discrete microvascular structure can be retained while simulations extend to large tissues, whole organs, and high-dimensional parameter spaces. By making these scales computationally accessible while retaining structure and relevant physics, the framework provides a practical foundation for high-throughput interrogation of questions in vascular biology and pathology, tissue engineering, and organ-scale transport that have previously been difficult to address without sacrificing either microscale fidelity or computational tractability.

## 4 Methods

Further details for all Methods sections, including derivations, implementation details, and rationales, are provided in the Supplementary Information. Parameter definitions and values are listed in Supplementary Tab. S3.

### 4.1 Sacrificial templates and perfusion chambers

Sacrificial channel architectures were created using the photopatterned-Channel Architectures with Sacrificial Templates (pCAST) approach following a protocol adapted from Coates *et al*. ^5^. Templates were 3D printed from 4-acryloylmorpholine (ACMO; Sigma-Aldrich) containing 5% (w/v) phenylbis(2,4,6-trimethylbenzoyl)phosphine oxide (BAPO; Sigma-Aldrich) using a prototype high-resolution S2 Continuous Liquid Interface Production (CLIP) printer (Carbon). Channels were manufactured to have a diameter of 250 *µ*m and were measured to be within roughly *±*5 *µ*m using an Olympus DSX Digital Microscope (DSX10-UZH).

Straight channels had a path length of 1.16 cm. Serpentine channels comprised three straight sections spaced 2 mm apart and connected by two curved sections, with a total path length of 3.09 cm. In both channel designs, the inlet and outlet ends were flared to 450 *µ*m for improved handling.

Perfusion chambers were 3D printed from KeySplint Hard Clear resin (Keystone Industries), sealed using custom gaskets made from EPU 40 (Carbon), and connected to the perfusion circuit using silicone tubing (Golander, 1 mm inner diameter). For oxygen measurements, the chamber base was replaced by a glass microscope slide carrying an oxygen-sensitive foil (SF-RPSu4-L4/W4-OIW; PreSens). Chamber components were sterilized by autoclaving; sensor-containing chambers were instead immersed in 70% ethanol and allowed to dry under sterile conditions.

### 4.2 Gelatin methacrylate preparation

Gelatin methacrylate (GelMA) was synthesized from cold-water fish-skin gelatin using a modified published protocol ^55^, yielding a degree of methacrylation of 72.3% as determined by proton nuclear magnetic resonance (NMR) spectroscopy (Supplementary Fig. S1). Mechanical characterization is reported in Supplementary Fig. S2 and Tab. S1. Complete synthesis, NMR, and mechanical characterization procedures are provided in the Supplementary Information.

### 4.3 Cell culture and cell-laden construct fabrication

Immortalized Human Embryonic Kidney 293T (HEK-293T) cells (gifted by Prof. Shaomin Tian, University of North Carolina at Chapel Hill, and Prof. Mark Skylar-Scott, Stanford University) were cultured in Dulbecco’s Modified Eagle’s Medium (DMEM; 10-013-CV, Corning) supplemented with 10% fetal bovine serum (FBS; F2442, Sigma-Aldrich) and 1% antibiotic-antimycotic solution (A5955, Sigma-Aldrich) at 37^*°*^C and 5% CO_2_ in a Forma Stericycle incubator (Thermo Scientific). Routine expansion and passaging procedures are described in the Supplementary Information.

For the engineered tissue constructs, HEK-293T cells were suspended at 60 million cells per mL in a precursor containing 10% (w/v) GelMA and 2 mM sterile-filtered LAP in 1*×* phosphate-buffered saline (PBS). The precursor was cast around a positioned pCAST template and photocrosslinked for 5 min using 395–405 nm irradiation (UltraFire). The sacrificial template was then dissolved and removed by gentle DPBS perfusion at approximately 10 *µ*L/min for ∼5 min using a BT100S-1 peristaltic pump (Golander). Sealed gasketed chambers were connected to the peristaltic pump by gas-permeable silicone tubing (1 × 0.92-S; Golander) to a medium reservoir fitted with a sterile gas-exchange filter.

Cell-laden constructs were perfused with cell culture medium at 25 *µ*L/min for 5 days in a humidified incubator at 37^*°*^C and 5% CO_2_, with medium replaced every 2 days (see Supplementary Fig. S3). At day 5, constructs were sectioned orthogonal to the channel direction into slices approximately 2 mm thick.

Static controls of the same cellular and hydrogel composition were cultured with channels in pCAST chambers wrapped in Parafilm.

HEK-293T cells were selected as a robust and tractable model for controlled validation of oxygen transport predictions, owing to their reproducible culture characteristics and established use in high-density perfused hydrogel constructs^5,6,56^. These experiments were intended to test spatial oxygenation and oxygen-limited survival in a controlled cellular system rather than to reproduce any tissue-specific physiology. GelMA was selected as the matrix material due to its relative ease of handling and established use in perfused engineered tissues^5,56,57^.

### 4.4 Live/dead staining and viability image analysis

Cell survival was evaluated using a live/dead viability assay (Thermo Fisher Scientific, L3224). Before staining, sliced cellular constructs were rinsed briefly with 1*×* DPBS. Samples were then incubated for 20 min at 37 °C in 1*×* DPBS containing calcein AM at 0.5 *µ*L/mL and ethidium homodimer-1 (EthD-1) at 2 *µ*L/mL. Calcein AM was used to label viable cells, while EthD-1 was used to identify cells with compromised membranes.

Live/dead-stained cross-sections were imaged using a Zeiss LSM980 laser scanning confocal microscope. Z-stacks were acquired with 488 nm excitation and a pinhole setting of 1 Airy unit. For each region of interest, a volume 300 *µ*m thick was collected using a 58.7 *µ*m spacing between optical sections, yielding six slices per stack. Detector settings were held constant across samples.

Confocal Z-stacks were converted into orthogonal maximum-intensity projections for quantification. Because red-stain retention was inconsistent among individual sections, cell survival was quantified from normalized green fluorescence rather than from the green-to-total fluorescence ratio. The resulting survival index was

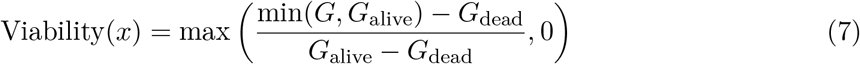

This quantity should be interpreted as an index of cell survival rather than a direct percentage of surviving cells. “Alive” reference intensities were measured just outside the dense green region immediately surrounding each perfused channel. “Dead” reference intensities were measured outside visibly viable regions, primarily at the furthest construct edges. Distance-resolved profiles were measured along lines spanning each construct approximately 100 *µ*m below the channel base. Presentation images were adjusted by red/green exposure changes and removal of background speckles outside the region of interest; quantitative analyses used unmodified image stacks.

### 4.5 Spatial oxygen measurements

Oxygen-measurement constructs were prepared as above, with the exception that an oxygen-sensitive foil was positioned beneath the hydrogel and constructs were perfused for 24 hours. After 24 hours at 25 *µ*L/min, chambers were switched from peristaltic to multichannel syringe pump perfusion at the same flow rate for more controlled, non-pulsatile flow. Sensor images were acquired from within the incubator using a VisiSens TD optical oxygen camera and analyzed using VisiSens ScientifiCal Software.

Each sensor was calibrated at 37^*°*^C using saturated aqueous sodium sulfite as the nominal zero-oxygen reference and air-equilibrated water as the high-oxygen reference, with the latter concentration determined from Henry’s law parameters ^58^. One-dimensional oxygen profiles were extracted along an x-averaged thin rectangular section perpendicular to flow through the center of the sensor field.

### 4.6 Simulations of engineered constructs

Finite-element method (FEM) simulations were performed as described below. CASCADE simulations used direct pairwise coupling with a source-contribution cutoff distance of 6*λ*, three external concentration field iterations, and a tissue-evaluation resolution of about 20 *µ*m. Prior work in a similar experimental system observed 20–30% loss of cells in well-perfused tissue after several hours of perfusion^5^. The nominal volumetric consumption rate was therefore prespecified at 0.7*V*_max_ to account for the corresponding reduction in metabolically active cell density. Poly-ACMO-cast channels within 10% (w/v) GelMA are known to swell by a factor of approximately 1.8–2 in diameter ^5^; this was also accounted for in the simulations. All simulation parameters and thresholds were specified before the experiments in this study were conducted. Sensitivity to the *V*_max_ multiplier and *C*^*\**^ is reported in Supplementary Fig. S20.

### 4.7 Synthetic vascular network generation

CASCADE accepts any network represented as circular, straight vessel segments with specified endpoints, radii, and connectivity. Biomimetic branching networks were generated using a modified version of svVascularize, an implementation of Constrained Constructive Optimization (CCO) ^6^. CCO sequentially adds vessels such that (a) all terminal segments have the same outgoing flow, (b) vascular volume is minimized, and (c) vessel radii obey Murray’s Law (shown in Eqn. 8 and derived in the Supplementary Info), a law which governs how the radii of daughter vessels relate to their parent vessel’s radius at a bifurcation. Supplementary Fig. S5 shows a representative diagram of a single optimization step.

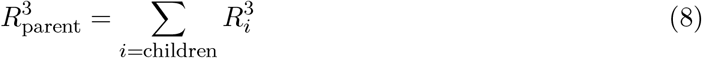

For large trees (≳300,000 vessels for unit-cube simulations, ≳700,000 vessels for heart simulations), full CCO insertion was followed by a faster terminal-only equal-bifurcation procedure. Selected terminal vessels were divided into two equal-radius daughter branches, after which the complete tree was rescaled to satisfy the prescribed root flow and/or pressure drop.

Tetrahedral, simple-cubic, body-centered-cubic, and octet lattices were generated using a simple graph connection algorithm on a Cartesian grid inscribed within a unit cube.

### 7.8 Predicting network hemodynamics

Each vessel segment was treated as a hydraulic edge *e* connecting two pressure nodes. Under steady, incompressible, fully developed, low-Reynolds-number flow, edge resistance was calculated according to Hagen-Poiseuille flow. The network was then solved similarly to how an electrical circuit is solved using Kirchhoff’s laws (see Supplementary Fig. S6). With *B* defined as the edge-node incidence matrix and 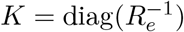 as the diagonal matrix of vessel conductances, nodal pressures and edge flows satisfy:

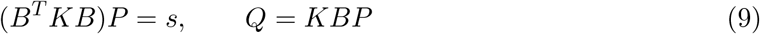

where *P* is the vector of nodal pressures and *s* contains prescribed nodal inflows (sources) and outflows (sinks). Pressure boundary conditions were imposed by eliminating prescribed nodes from the resulting sparse conductance-weighted graph Laplacian, which was solved using a sparse direct solver (SciPy spsolve). The solver supports two boundary condition modes: (a) prescribed inlet flow and terminal pressure, as was done in Fig. 4 (in Figs. 4a–d: 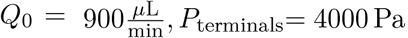) and Figs. 5a–d,f; and (b) prescribed inlet and terminal pressures, as was held constant in Fig. 5e (*P*_0_= 12665.6 Pa, *P*_terminals_= 4000 Pa). Cell-culture medium was treated as a Newtonian fluid with dynamic viscosity of ∼0.7 cP at 37^*°*^C. For blood-perfused networks, diameter- and hematocrit-dependent apparent viscosities and red-blood-cell partitioning were calculated by Pries-Secomb Fåhræus–Lindqvist and phase-separation relations implemented through NetFlowV2^20,59–66^. Flow and hematocrit were recomputed iteratively until convergence (0.1% relative change). Baseline inlet discharge hematocrit was 0.42.

### 4.9 Intraluminal oxygen transport

After the hydraulic solve, edges were oriented according to the computed flow direction. Oxygen transport was represented on the same directed graph, with oxygen-conserving nodal balances at junctions. Over each vessel edge, intravascular transport was reduced to an affine update,

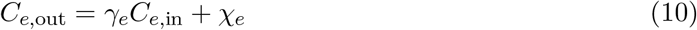

where *γ*_*e*_ describes attenuation of the incoming concentration and *χ*_*e*_ accounts for exchange with surrounding tissue. At each internal node *i*, conservation of advected oxygen requires

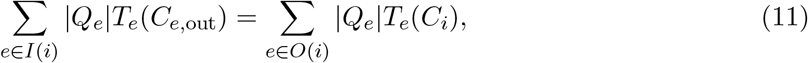

with total oxygen content 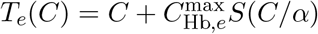 and hemoglobin oxygen capacity 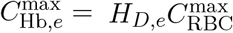 where *I*(*i*) and *O*(*i*) denote incoming and outgoing edges, *C*_*i*_ is the mixed dissolved-oxygen concentration, *H*_*D,e*_ is discharge hematocrit, 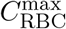 is the fully saturated oxygen capacity per unit red-cell volume, *α* is the Henry’s law coefficient, and *S*(*P*) is the Severinghaus oxyhemoglobin-dissociation curve^67^. Without hemoglobin present, *T*_*e*_(*C*) = *C*, equivalent to flow-weighted mixing.

Vessel edges were advanced through multiple (typically 5) substeps, over which vessel radius, flow rate, external concentration, and local transport coefficients were held constant. Rather than assuming that oxygen was uniformly mixed across the lumen, finite radial diffusion under Poiseuille flow was resolved using a high-Péclet-number Graetz eigenfunction expansion, subject to radial symmetry at the centerline and continuity of oxygen flux at the vessel wall. The resulting wall concentration for a subsegment *i* was

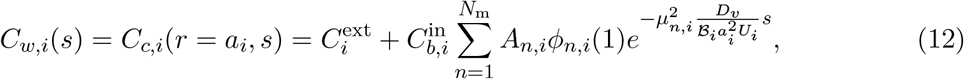

where 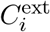 is the local external concentration, 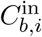 is the incoming bulk dissolved-oxygen concentration, *A*_*n,i*_, *ϕ*_*n,i*_, and *µ*_*n,i*_ are the modal coefficients for dimensionless excess concentration, radial eigenfunctions, and eigenvalues, respectively. *D*_*v*_ is the intravascular oxygen diffusivity, and *a*_*i*_ and *U*_*i*_ are the vessel radius and mean velocity. The corresponding flow-weighted bulk concentration is recovered as

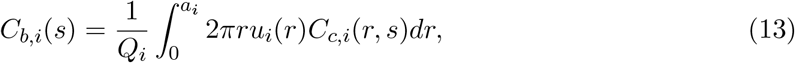

where *C*_*c,i*_(*r, s*) is the intraluminal concentration field. *C*_*b*_ is the flow-weighted bulk dissolved-oxygen concentration, while *C*_*w*_ determines the outward tissue flux according to Eqn. 15. For blood, dissolved and hemoglobin-bound oxygen were assumed to remain in instantaneous equilibrium, with total oxygen content calculated as 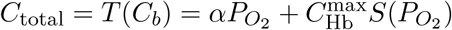. The corresponding frozen hemoglobin-buffer factor was 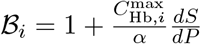, with *ℬ*_*i*_ = 1 for cell-culture medium. See Supplementary Fig. S7 for two examples of this intraluminal solve.

For Hagen-Poiseuille flow, the eigenvalues and velocity-weighted modal coefficients of the Graetz eigenproblem were precomputed over a lumen-tissue Biot number Bi_*i*_ = *κ*_*i*_*/*(2*πD*_*v*_) and interpolated at runtime. External field concentrations were determined by evaluating a modified form of Eqn. 3 along the target vessel centerline. The intravascular graph and external-field solutions were coupled through iterations. Intraluminal axial diffusion was neglected under the high-Péclet-number approximation. Tissue-side axial diffusion was neglected in the local radial exchange closure; its effect on radial exchange was assessed in Supplementary Appendix F and Fig. S16.

### 4.10 Yukawa-Green’s-function tissue oxygen theory

Tissue oxygen was computed using Eqn. 3. The equation was evaluated using a locally linearized reaction rate according to the tissue reaction-diffusion length *λ*_*t,i*_ and using the segment flux per unit length *q*_*i*_(*s*), radius *a*_*i*_, wall concentration *C*_*w,i*_(*s*), Green’s-function *G*_*i*_, and orientation vector 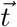.

For *λ*_*t,i*_, the effective concentration was estimated via evaluation of the single-vessel modified cylindrical Bessel function solution

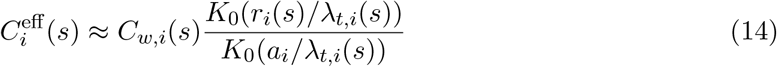

where *K*_0_ is the zeroth-order modified Bessel function of the second kind.

The outward flux per unit length from each vessel was defined through a linear driving force flux law,

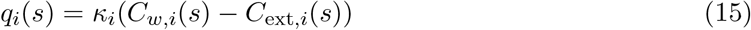

where for a vessel *i, q*_*i*_ is the outward flux per unit length, *κ*_*i*_ is the wall transfer coefficient, *C*_*w,i*_ is the internal vessel-wall concentration, and *C*_ext,*i*_ is the external tissue concentration imposed by surrounding vessels. In the definition of *κ*, the effective concentration is the vessel wall concentration 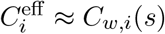.

Using the locally frozen first-order reduction of Eqn. 3, tissue concentration was evaluated as

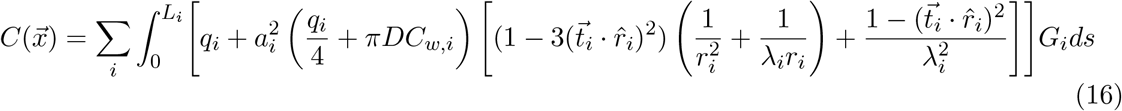

### 4.11 Simulated viability thresholding

Tissue was classified as viable or nonviable by comparing the predicted dissolved oxygen concentration with a prespecified critical concentration *C*^*\**^. All analyses in Fig. 4 estimated tissue viability from 1 million point samples (100*×*100*×*100 Cartesian grid). For Fig. 4a, slices were evaluated at a resolution of 500*×*500 (20*µ*m spacing) and interpolated by a single-closest-neighbor Gaussian interpolation kernel. Biventricular heart outputs were analyzed using 8 million equally spaced points and displayed viability maps in Figs. 5a–c were subject to the same interpolation kernel.

### 4.12 Tissue-scale design space analysis definitions

Distance to nearest channel wall (DNCW), as shown in Fig. 4b, was defined as the minimum distance to a segment centerline minus that segment’s local radius. This is distinct from the mean vessel-to-vessel spacing *L*_*v*_ used in the definition of Π. In the definition of Π, we set the effective concentration as the inlet dissolved oxygen concentration *C*^eff^ = *C*_0_.

For the general design maps shown in Figs. 4e–g, we varied inlet dissolved concentration (*C*_0_ = 0.07–0.16 mol/m^3^), inlet flow (*Q*_0_ = 0.005–20, 000 *µ*L/min, logarithmically distributed), discharge hematocrit (*H*_*D*_ = 0–0.42), maximum reaction velocity (*V*_max_ = 0.5 *™* 40 mmol/(m^3^s)), Michaelis-Menten constant (*K*_*M*_ = 0.69–3.6 mmol/m^3^), tissue diffusivity (*D* = 10^*™*12^–10^*™*9^ m^2^/s), and viability thresholds (*C*^*\**^ = 1.4–7 mmol/m^3^). For Fig. 4f, branched networks were compared with tetrahedral (diamond cubic), simple cubic, body-centered cubic, and octet lattices under matched physical conditions. Viable tissue fraction was adjusted to account for the volume occupied by the vessels 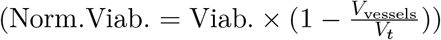. Lattice radii ranged from 10 to 500 *µ*m, with geometrically infeasible radii excluded. For each lattice and physical condition, the radius with the maximum normalized viability was retained.

### 4.13 Creation of the biventricular heart model

The biventricular myocardial geometry (3DPX-002636) was obtained from the NIH 3D repository and represents a healthy 17-year-old female ^68^. Coronary arteries were segmented from Heart0069 of the Atlas of Human Cardiac Anatomy from the University of Minnesota^69^. The resulting left and right coronary artery graphs were extended through the myocardium using the aforementioned generative design techniques. The final geometry was represented by 125 million quadrature nodes, distributed between the right and left coronary trees in proportion to their prescribed inlet flows. Further details, including simulation parameters, are available in the Supplementary Information.

### 4.14 Coronary and systemic perturbations

For the occlusion study, eight proximal coronary sites were perturbed independently, as shown in Fig. 5d, corresponding to the first and second left diagonal branches (LD1, LD2), left anterior descending (LAD), first and second left marginal (LM1, LM2), right conus (RC), right marginal (RMA), and posterior descending artery (PDA). The LAD occlusion site was placed immediately after the bifurcation corresponding to the LD2, as opposed to above the LD1 or LD2. Myocardial watershed territories were created during generative growth by assigning each new-growth point to its closest vascular segment or downstream arterial subtree. The radius of each target coronary segment was progressively reduced from the healthy baseline to complete occlusion, with local resistance updated according to

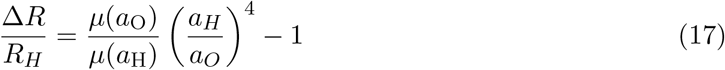

where *H* denotes the healthy case and *O* an occluded case.

For the hematocrit sweep, as seen in Fig. 5e, inlet discharge hematocrit was varied from 0% to 80% Hct while coronary pressure drop, geometry, and all other conditions were held fixed. Diameter-dependent viscosity, phase separation, tube hematocrit, and hemoglobin binding capacity were iteratively recomputed as with all other studies in this work. Clotting, erythrocyte aggregation, and autoregulatory responses were not included.

For systemic hypoxia, inlet arterial dissolved oxygen concentration was varied from 0 to 0.14 mol/m^3^ (0–100 mmHg) while all other parameters were held at baseline. At each inlet value, incoming hemoglobin saturation was recomputed via the Severinghaus relation. Digitized experimental cardiac output and pressure-volume-area (PVA) data were used only for external physiological comparison. These values, as shown in Fig. 5f, were normalized by their respective maxima.

### 4.15 Numerical implementation details

The solver generally follows the diagram shown in Supplementary Fig. S4. Pairwise Green’s-function calculations used k-d trees to restrict exact source-to-target evaluations. Unless otherwise stated, a target considered sources within six local reaction-diffusion lengths (6*λ*).

For the FFT implementation, quadrature-weighted vessel sources were deposited by triangular-shaped-cloud (TSC) assignment on a fixed and padded Cartesian grid. Unless otherwise stated, the resolution of this grid was 288*×*288*×*288 (per cm^3^), corresponding to an average relative spacing of Δ*x/λ ≈* 0.44. Sources were divided into five dynamic quantile bins in *λ* (see Supplementary Fig. S14). For each bin, the screened Poisson equation was solved in Fourier space with cuFFT, after which fields were summed and interpolated to vessel or tissue targets. Smoothed same-quadrature self-contributions were subtracted. Whole-heart simulations did not employ the FFT solver.

Nonlinear coupling between vessel concentration and the external field used adaptive Aitken relaxation with safeguarded Anderson acceleration.

Further solver details and analyses are provided in the Supplementary Information. See Supplementary Figs. S11–S13, S15 for analyses on the sensitivity of *Nλ* cutoff distances, *C*_ext_ iteration counts, FFT grid resolution, and Graetz intraluminal eigenfunction resolution.

The solver was implemented in Python 3.9.20 and GPU calculations used CUDA 13.0, cuFFT 12.2.0.46, and 20 custom CUDA C kernels invoked through CuPy RawKernel. *K*_1_*/K*_0_ Bessel solution ratios and Graetz solution bases were interpolated from precomputed lookup tables over *a/λ* and Biot number (Bi) values, respectively. Blood transport used five axial substeps per segment due to the nonlinearity in the Severinghaus relationship.

### 4.16 Numerical validation via FEM

Finite-element method (FEM) simulations were performed in COMSOL Multiphysics 6.3 using explicit vascular lumens coupled to a three-dimensional tissue domain. Prescribed conditions included inlet flow and oxygen concentration, open external boundaries (to agree with the Green’s infinite tissue assumption), no-slip walls, and continuity of oxygen concentration and flux at the vessel-tissue interface. Tissue oxygen consumption was modeled using Michaelis-Menten kinetics. All simulations were performed at steady state using P2+P1 elements (quadratic velocity, linear pressure) for flow and quadratic elements for oxygen concentration.

Twenty-five validation cases were constructed with tissue volumes of 0.037, 0.125, and 1 cm^3^ and vascular networks containing tens to hundreds of segments. Cases spanned Π=0.5–3.4 and Φ=0.2–93.5 through variation of geometry, inlet conditions, and transport properties. See an example visual comparison in Supplementary Fig. S8. Vessel surface meshes were generated via an implicit-web meshing algorithm detailed in the Supplementary Information (Section 8). See three examples in Supplementary Fig. S9. Tetrahedral lumen and tissue meshes were generated in COMSOL Multiphysics and were refined with COMSOL adaptive mesh refinement (using the “L2 norm of error squared” criterion and COMSOL’s “Automatic” tolerance setting). Boundary-layer elements were added near the lumen wall and further local refinement was performed at bends and junctions.

For each case, 10,000 locations were placed on a uniform Cartesian grid. Sampling locations returning NaN were excluded, leaving 233,140 pooled comparison points. Agreement was quantified using the concordance correlation coefficient (CCC), root mean squared error (RMSE), mean absolute error (MAE), coefficient of determination (R^2^), and mean signed error. A low-concentration subset was prespecified as all points with FEM reference concentrations below 0.005 mol/m^3^. Classification accuracy, false-alive rate, and false-dead rate were calculated using a critical concentration of *C*^*\**^ = 0.0014 mol/m^3^, treating the FEM result as the reference. All points were weighted equally. These analyses were performed for both the Green’s method developed in this work (Supplementary Fig. S10, Tab. S2) and an “optimized” Krogh model (Fig. S19, Tabs. S4–S5). Fig. S18 compares the Green’s and Krogh’s models on two simple geometries. For the Krogh model (described in Supplementary Appendix J and illustrated in Supplementary Fig. S17), we evaluated eight methods of inscribing Krogh radii (where we selected the best-performing formulation), while independently fitting the zeroth-order oxygen consumption rate for each validation case against the FEM reference, thereby providing a deliberately favorable benchmark for comparison with the Green’s model.

### 4.17 Statistical analyses

Experimental data are shown as mean *±* standard deviation. Error bands in Figs. 2f,j do not include sensor noise. Exact N values are reported in each figure legend and correspond to the number of samples, not the number of measurements. No null-hypothesis significance testing was performed.

Wall-clock measurements in Fig. 3a,b are reported as *µ ± σ* on five independent runs. Phase timings were measured using time.perf_counter, with the default CUDA stream synchronized before ending each GPU timing interval. Whole-run elapsed times included model loading, flow and oxygen solves, and tissue evaluation. Line plots in Fig. 3c show sweeps across both tissue and vessel resolution (*M, N*) with a single evaluation for each point recorded, using each of the two modes of operation.

### 4.18 Hardware and operating system

All computations were performed on a Dell Precision 7560 laptop with an 11th-generation Intel Core i9-11950H CPU at 2.60 GHz, 64 GB system memory and an NVIDIA GeForce RTX 3080 Laptop GPU with 16 GB dedicated memory. The production environment was Ubuntu 24.04.3 LTS under Windows Subsystem for Linux 2.

## Data availability

Raw or processed datasets are available from the corresponding author upon reasonable request.

## Code availability

The CASCADE code is available on GitHub at https://github.com/carlkohnke/CASCADE under a license permitting academic, noncommercial use. Redistribution of the software or derivatives requires prior written permission from Stanford University.

## Acknowledgments

We thank Saksham Malik, Manasvi Vora, Vedika Shenoy, Elias Matthews, John MacArthur, and Mark Skylar-Scott for fruitful discussions and advice. We are thankful for equipment access through the Soft & Hybrid Materials Facility (SMF), part of the Stanford Nano Shared Facilities (SNSF).

## Author contributions

Conceptualization: CAK, IAC, ZAS, ALM, JMD, ESGS; Experimental methodology: CAK, YLT, IAC, JMD; Theory: CAK, ESGS; Computational methodology: CAK, ESGS; Software: CAK, ZAS; Cell culturing: CAK, YLT, IAC; Validation (experimental): CAK, YLT, IAC; Validation (computational): CAK, ESGS; Analysis and investigation: CAK, YLT, IAC, ZAS, ALM, JMD, ESGS; Visualization: CAK; Funding acquisition: JMD, ESGS; Supervision: ALM, JMD, ESGS; Writing (draft): CAK; Writing (review & editing): CAK, YLT, IAC, ZAS, ALM, JMD, ESGS.

## Artificial intelligence usage statement

Large language models were used to assist with coding and manuscript revisions prior to peer review. The authors developed the ideas, performed the scientific work, and reviewed all manuscript content.

