## Supplementary Information for "Rapid simulation of organ-scale oxygen transport for tissue ischemia prediction"

**C Reducing the pairwise problem to accelerate tissue and external concentration field calculation** 42

**D Rapid evaluation of the external field $C^{\text{ext}}$ using FFT on a regular grid** 43

**E Graetz eigenfunction resolution assessment for intraluminal diffusion** 47

**F Impact of neglecting axial diffusion $D\frac{\partial^2 c}{\partial s^2}$ in the diffusion operator $D\nabla^2 c$.** 47

**H Murray's Law derivation** 51

**I Net resistance scaling** 51

**J Krogh cylinder model derivation and comparison to FEM/Green's** 52

**K Sensitivity of $V_{\text{max}}$ multiplier and $C^*$ choice in simulations of GelMA constructs** 57

**L Additional methodological details** 58

### 1 Tissue engineering experiments

#### 1.1 Materials and templating technique

Note that the majority of this section matches, or is very similar to, the protocol described in the Supplemental Information of *Coates, I.A. et al. (2026)* [4].

##### 1.1.1 Sacrificial template synthesis

**pCAST resin preparation:** The pCAST printing resin was prepared by combining 4-acryloylmorpholine (ACMO; Sigma-Aldrich) with phenylbis(2,4,6-trimethylbenzoyl)phosphine oxide (BAPO; Sigma-Aldrich) at 5% (w/v). The resin mixture was sonicated for approximately 10 min to fully dissolve the photoinitiator. Prepared resin was protected from light and used for printing on the day it was made.

**CLIP printing of pCAST templates:** Template designs were generated in Autodesk Fusion, and slicing and support structures were prepared using Carbon software. All pCAST template structures were fabricated using a prototype S2 high-resolution Continuous Liquid Interface Production (CLIP) printer from Carbon.

Following printing, residual resin trapped within template features was carefully cleared using compressed air. The printed templates were then rinsed briefly in isopropyl alcohol (IPA; Sigma-Aldrich), dried again with compressed air, and post-cured for 6 min in a UV curing oven (APM Technica). Completed templates were kept protected from light until use.

##### 1.1.2 Fabrication of perfusion chambers

Perfusion chamber bodies and covers for cell culture studies were designed in Autodesk Fusion and sliced using Carbon software. Chamber components were printed on a Carbon M1 printer using KeySplint Hard Clear dental resin (Keystone Industries). Custom gaskets were printed on a Carbon M2 printer using Carbon EPU 40 resin to create an airtight interface between the chamber body and cover. The gasket was compressed using screws to produce a leak-resistant seal, and 1 mm silicone tubing was connected to the printed parts for perfusion via a peristaltic pump. The chamber's internal basin for perfusion was an outward-tapered rectangular prism centrally measuring 1.06×1.00×0.55 cm at its widest.

Perfusion chambers used for oxygen measurements were produced using the same general workflow, with one difference. The chamber bottom was left open during fabrication, after which a glass microscope slide was cured in place to serve as the base. An oxygen-sensitive foil sensor (PreSens SF-RPSu4-L4/W4-OIW) was attached to the glass surface using transparent double-sided 3M tape.

Before cell-based experiments, all chamber components were sterilized by autoclaving. The only exceptions were the printed gaskets and oxygen-sensing chambers containing the sensor foil, which were sterilized by soaking in 70% ethanol (LabChem) and allowed to dry prior to use.

##### 1.1.3 Preparation and characterization of gelatin methacrylate (GelMA)

**GelMA synthesis:** Gelatin methacrylate (GelMA) was produced from cold-water fish skin gelatin using a modified methacrylation reaction based on a previously established protocol [45]. To begin the synthesis, 60 g of cold-water fish skin gelatin (CWF gelatin; Sigma-Aldrich, G7041) was added to 300 mL of 0.1 M carbonate–bicarbonate buffer. The mixture was held at 50 °C for approximately 1 h with mixing until the gelatin was fully dissolved.

After dissolution, the solution was adjusted to pH 10 using 10 M sodium hydroxide (Sigma-Aldrich, 415413). The temperature was then increased to 70 °C before initiating the methacrylation reaction. Methacrylic anhydride (5 mL; Sigma-Aldrich, 276685) was added to the gelatin solution while stirring at 100–200 rpm. The reaction was allowed to continue for 2 h at 70 °C. During this period, the reaction vessel was covered with aluminum foil to limit evaporative loss.

Once the reaction was complete, the mixture was allowed to cool to room temperature. GelMA was then isolated by pouring the reaction solution into an excess volume of absolute ethanol, corresponding to approximately three times the reaction volume (≥99.5%; Sigma-Aldrich, 459844). The precipitated polymer was collected and dried for more than 24 h. After drying, the material was rehydrated in 100 mL of deionized water and heated to approximately 80 °C for 1–2 h with stirring to assist in removing residual ethanol. The resulting GelMA stock solution was cooled and stored at 4 °C. For experiments involving cells, the GelMA stock was sterilized by autoclaving before use.

The concentration of the GelMA stock was measured by drying a known volume of solution and determining the remaining polymer mass. Specifically, an aliquot of GelMA solution was lyophilized overnight, and the dry weight was used to calculate the stock concentration. The stock solution was then diluted as needed to prepare GelMA formulations for subsequent experiments.

**Determining methacrylation extent via NMR:** The extent of gelatin methacrylation was quantified using proton nuclear magnetic resonance spectroscopy ($^1$H-NMR). A small portion of the GelMA stock was first lyophilized to obtain dry polymer. Dried GelMA and unmodified CWF gelatin were then prepared as separate samples by dissolving 5 mg of each material in 1 mL of $D_2O$ (Sigma-Aldrich, 151882). Spectra were collected on a 600 MHz NMR spectrometer (Varian). Following acquisition, spectra were phase corrected, baseline corrected, and integrated using MesReNova.

For analysis, the aromatic amino acid region at 7.05–7.25 ppm in $D_2O$ was used as the internal normalization signal. The lysine-associated signal at 2.80–2.90 ppm in $D_2O$ was then compared between native gelatin and GelMA. Because methacrylic anhydride consumes primary amines on lysine residues during GelMA formation, successful modification is reflected by a reduction in the normalized lysine peak area. The degree of methacrylation was calculated from the relative decrease in this lysine signal according to:

$$\text{Degree of methacrylation} = \left[1 - \frac{\left(I_{\text{lysine}}/I_{\text{aromatic}}\right)_{\text{GelMA}}}{\left(I_{\text{lysine}}/I_{\text{aromatic}}\right)_{\text{CWF gelatin}}}\right] \times 100\%.$$

Using this approach, the synthesized GelMA had a methacrylation degree of 72.3%. Figure S1 shows NMR spectra of CWF gelatin and GelMA samples with labeled peaks of interest.

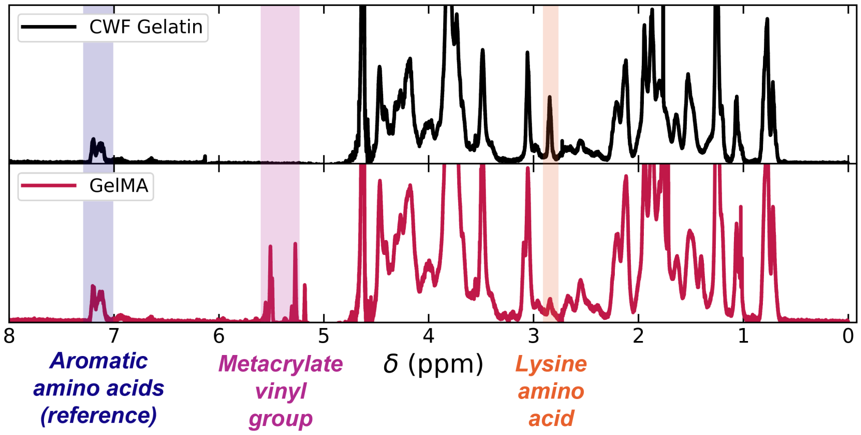

Figure S1: Representative $^1$H-NMR spectra of CWF gelatin (top) and GelMA (bottom) samples.

**GelMA compression testing:** To prepare samples for mechanical characterization, GelMA precursor solutions were formulated at 10% (w/v) in 1× Dulbecco's phosphate-buffered saline (DPBS; Corning, 21-031-CV). Lithium phenyl-2,4,6-trimethylbenzoylphosphinate (LAP; Sigma-Aldrich) was included at a final concentration of 2 mM as the photoinitiator. The precursor solutions were transferred into cylindrical molds with dimensions of 10 mm diameter and 3 mm height. The samples were then crosslinked for 5 min using 395–405 nm light from a handheld UV flashlight (UltraFire UV).

Following photocrosslinking, the GelMA discs were removed from the molds and immersed overnight in 1× DPBS at room temperature to equilibrate. Before mechanical testing, excess buffer was gently removed from the surface of each disc using a Kimwipe. Unconfined compression testing was performed with an Instron 5565 mechanical tester equipped with a 100 N load cell. Samples were compressed at a strain rate of 20% min$^{-1}$. Table S1 shows the resulting compressive moduli at 5%, 10%, and 15% strain. Figure S2 shows the compressive stress-strain curves as collected from the load cell.

| Strain | Tangent modulus, avg ± SD | Secant modulus, avg ± SD |
|---|---|---|
| 5% | 3.34 ± 0.74 kPa | 2.97 ± 0.58 kPa |
| 10% | 5.29 ± 0.95 kPa | 3.75 ± 0.74 kPa |
| 15% | 9.58 ± 2.86 kPa | 4.88 ± 0.88 kPa |

Table S1: Compressive tangent and secant moduli calculated from stress–strain measurements at 5%, 10%, and 15% compressive strain. Values are reported as mean ± standard deviation across three samples.

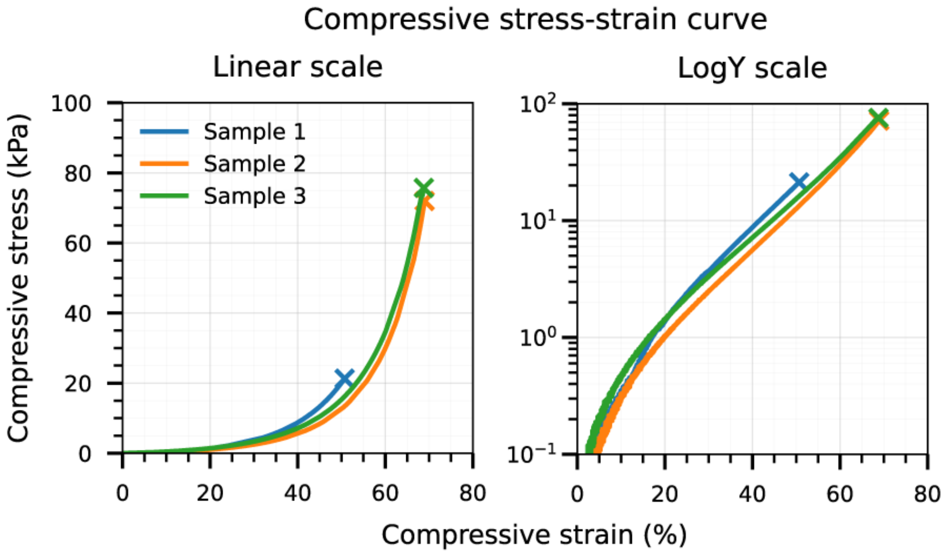

Figure S2: Compressive stress-strain curves for three GelMA samples. Curves are truncated upon material failure. The left plot uses a linear y-axis; the right plot uses a logarithmic y-axis.

#### 1.2 Cellular construct preparation, perfusion, and analysis

##### 1.2.1 Cell culturing

All cell culture work was carried out in a humidified incubator maintained at 37°C with 5% $CO_2$. Immortalized human embryonic kidney 293T cells (HEK-293T) were provided by Prof. Shaomin Tian at the University of North Carolina at Chapel Hill and Prof. Mark Skylar-Scott at Stanford University, and were expanded in Dulbecco's Modified Eagle's Medium (DMEM; Corning, 10-013-CV). HEK-293T cells were chosen due to their low batch-to-batch variability, moderate oxygen consumption kinetics, and common use in 3D tissue engineering literature. The culture medium was supplemented with 10% fetal bovine serum (FBS; Sigma-Aldrich, F2442) and 1% antibiotic-antimycotic solution (Anti-anti; Sigma-Aldrich, A5955).

Cultures were refreshed with new medium every 2 days and subcultured at a 1:4 split ratio once they reached confluence. Before enzymatic detachment, flasks were washed with 1× DPBS. Cells were then incubated with 0.1% Trypsin-EDTA (Sigma-Aldrich, T4049) for 5 min at 37°C. Trypsinization was stopped using medium containing 10% FBS, after which the cells were pelleted by centrifugation at 1000 rpm for 4 min. The resulting cell pellets were resuspended in fresh medium or buffer before downstream seeding.

##### 1.2.2 Perfusion of cell-laden pCAST constructs

Cells were collected and resuspended in 1× DPBS before incorporation into the hydrogel precursor. The cell-laden precursor formulation contained 10% (w/v) GelMA and 2 mM sterile-filtered LAP photoinitiator. This formulation was used for all reported cell-containing constructs.

To fabricate cellularized perfusable constructs, the cell-containing precursor solution was pipetted into the chamber around the positioned pCAST template. The material was then photocrosslinked immediately for 5 min using 395–405 nm irradiation from a handheld UV source (UltraFire). Following the same template-removal procedure used for the aqueous GelMA constructs, the sacrificial pCAST template was dissolved and removed through gentle DPBS perfusion to generate an open channel. To ensure that residual template material was eliminated from the lumen, constructs were additionally flushed with 1× DPBS for more than 5 min at about 10 $\mu$L/min using an external peristaltic pump (Golander BT100S-1). After this flushing step, the constructs were connected to the culture-medium perfusion circuit.

For perfusion culture, each cell-laden construct was sealed inside a chamber using a gasketed cover. This configuration helped maintain sterility, reduce evaporative losses, and restrict oxygen transport through the chamber surfaces so that the internal channels served as the dominant route for mass exchange. The sealed chambers were connected to an external peristaltic pump and medium reservoir using gas-permeable silicone tubing with a 1 mm inner diameter (Golander, 1×0.92-S). The medium reservoir was equipped with a sterile filter port to allow gas exchange.

An image of our perfusion setup immediately prior to being placed inside the incubator and undergoing 5-day perfusion is shown in Figure S3. In this example, we were running six cellular constructs.

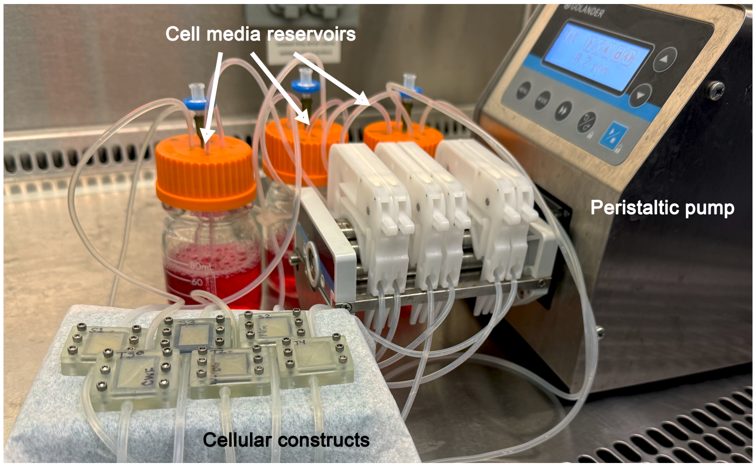

Figure S3: Image of perfusion setup after pCAST channels were flushed but before 5-day perfusion. Shown is our setup for a 6-construct run.

Constructs were perfused for 5 days at 25 $\mu$L/min with DMEM. The entire perfusion setup was maintained inside a standard cell culture incubator at 37 °C and 5% $CO_2$. Each 40 mL DMEM reservoir supplied 1–3 perfusion chambers, and the medium was replaced every 2 days.

At the end of the 5-day culture period, perfused constructs were sectioned into cross-sections approximately 2 mm thick. These sections were then processed for viability staining and imaging.

Negative control constructs were also built and imaged, following the same procedure (including channel embedding) as above, except with no perfusion.

#### 1.3 Spatial oxygen tracking experiments

Oxygen tracking experiments were set up similarly to the constructs described in the above sections, except: the chamber bottom was replaced with a glass slide, to which an oxygen-sensitive photoluminescent foil (SF-RPSu4-L4/W4-OIW; PreSens) was attached with transparent, double-sided tape (3M); the constructs were perfused for 24 hours, as opposed to 5 days; the constructs were not sliced and live/dead stained, but rather were subjected to the measurements described in the following paragraph.

$O_2$-tracking chambers were seeded with cells and perfused with a peristaltic pump at 25 $\mu$L/min for 24 hours. Each chamber was then switched to a multi-channel syringe pump for higher-precision flow rate control at low flows. Inside the incubator at 37°C, images were taken of the chamber sensors using a VisiSens TD optical $O_2$ tracking camera.

Sensor intensity values were compared to images with known oxygen values for calibration. For the low end of oxygen sensor calibration measurements, we prepared saturated solutions of sodium sulfite (a strong oxygen scavenger) in water and measured sensor readout when covered with this solution. Furthermore, we measured oxygen values immediately after photocrosslinking of GelMA placed atop the sensor. Both measurements produced the same intensity values, and were assumed to have minimal oxygen. For the high end of sensor calibration measurements, we measured the intensity of sensors covered with water in equilibrium with air. All measurements were taken at 37°C.

Sensor readouts were analyzed using VisiSens ScientifiCal and converted to heatmaps of oxygen concentration.

### 2 Overall solver structure & introduction

Figure S4 shows the general architecture of our method and how arguments/solutions are passed down the ladder of interconnected solvers. The calculations shown in the following sections are performed by the solver in the order indicated here.

We present the following sections roughly in the same order they appear in the solver. We cover generative design of vascular architectures (Section 3), our 0D blood flow solver (Section 4), network intravessel concentration solver (Section 5), and our tissue concentration solver (Section 6). We also cover mesh generation and comparison to Finite Element Method (FEM) simulations (Section 8), big $O$ time scaling, and numerical/algorithmic tricks for high-speed computation (Section 9).

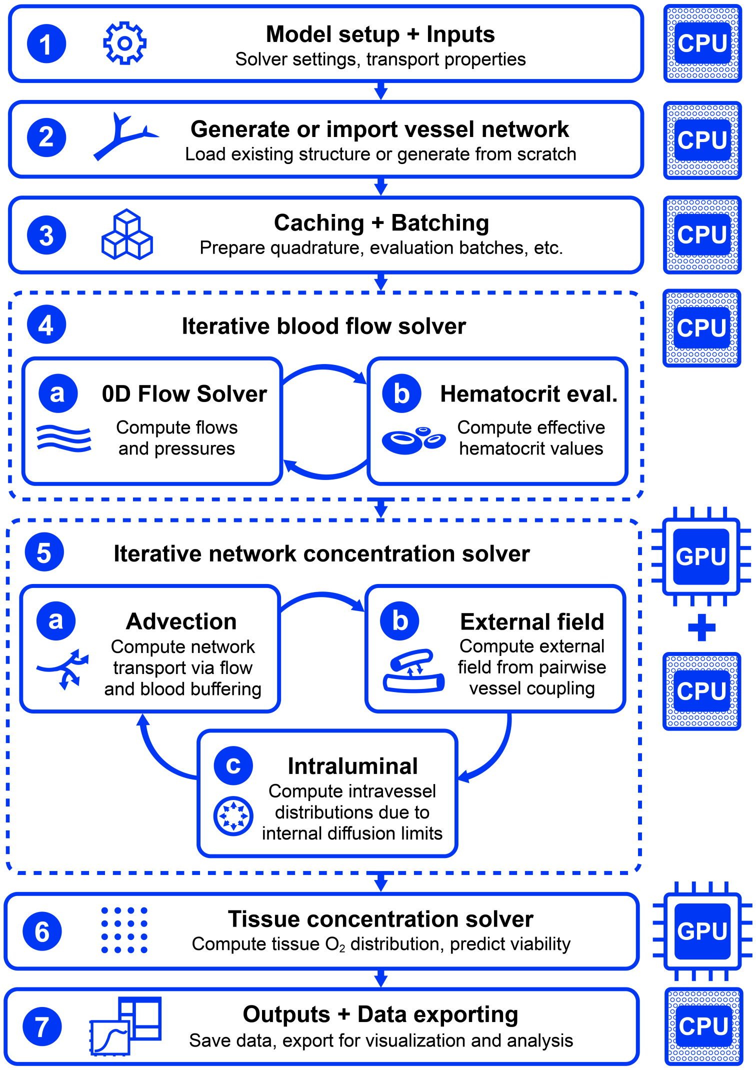

Figure S4: Flowchart showing the major steps in the solver and whether they are processed by the CPU or GPU.

### 3 Generating biomimetic vascular trees

The semianalytical simulation techniques used in this solver are agnostic to the vascular structure being analyzed (as long as the channels are circular in cross-section).

Generative design of vascular structures is a rich field of study, offering several methods of approach. Several algorithms have been developed using fractal rules [11, 19, 22], Growth Based Optimization (GBO) [10, 15], Constrained Constructive Optimization (CCO) [1, 10, 13, 15, 38, 42, 49], stochastic L-systems [7], etc.

In this work, we generate vascular structures using Constrained Constructive Optimization (CCO). Specifically, we use the SimVascular implementation of this method, svVascularize, which uses spatial indexing and partial binding techniques to accelerate the algorithm's most intensive steps. Please see *Sexton, Z.A. et al. (2025)* [42] for more details of the svVascularize implementation.

#### 3.1 Constrained Constructive Optimization (CCO)

CCO constructs a binary vascular tree incrementally. Starting from a root vessel or an imported "seed" vascular structure, the algorithm repeatedly samples a new growth target point inside the tissue domain and determines the lowest-cost way to connect that terminal to the existing tree. For each proposed terminal, nearby existing vessels are considered as possible parent segments, the local bifurcation point is optimized, geometrically infeasible insertions are rejected, and the feasible candidate with the lowest tree cost is accepted.

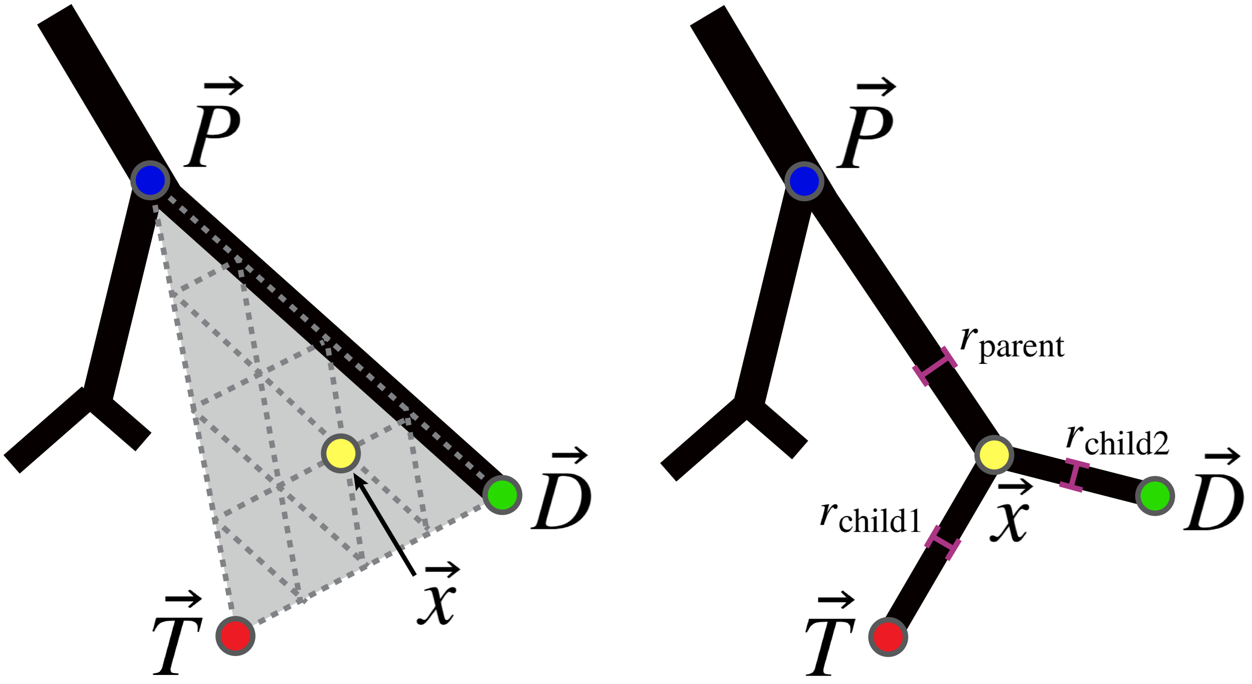

Figure S5: Diagram of the CCO bifurcation optimization problem (left) and a committed terminal vessel growth (right). The placement of point $\vec{x}$ is optimized, affecting the geometry of the edges connecting to $\vec{P}$, $\vec{D}$, and $\vec{T}$, as well as the radii of the rest of the tree.

##### 3.1.1 Cost function

Each candidate vessel insertion is evaluated by the resulting cost of the vascular tree. The tree is represented as a collection of cylindrical vessel segments, and the objective (cost function) is written as

$$J = \pi \sum_{e} r_e^{\alpha} L_e^{\beta}$$

Where $r_e$ and $L_e$ are the radius and length of each segment (edge) $e$. In the usual case, $\alpha = 2$ and $\beta = 1$, so $J$ is proportional to intravascular volume.

##### 3.1.2 Radius determination

The radii of the vessels are not chosen independently for each vessel. Instead, once a candidate bifurcation geometry has been proposed, the algorithm determines the daughter radius ratios from the downstream flow demands and hydraulic resistances. Then, the tree is rescaled so that the prescribed root flow $Q_{\text{in}}$ is achieved under the imposed pressure drop $\Delta P$.

The terminal (outlet) vessels are assigned equal flow, as the tree is intended to have $N_t$ terminals with equal flow rates $q_t$ through them, adding to the inlet flow rate $Q_{\text{in}} = q_t N_t$.

The flow through any edge $e$ is therefore $Q_e = n_e q_t$ where $n_e$ is the number of terminal outlets downstream of that segment. A branch supplying many downstream terminals naturally carries more flow and receives a larger radius than a branch supplying only one or a few terminals.

For a cylindrical vessel segment, Poiseuille resistance scales as

$$R_e = \frac{8\mu L_e}{\pi r_e^4}$$

It is therefore convenient to separate the length-dependent part of the resistance from the radius-dependent $r^{-4}$ scaling. For a candidate bifurcation point $\vec{x}$, the reduced resistance of the new terminal branch is

$$R_t = \frac{8\mu}{\pi} ||\vec{x} - \vec{T}||$$

while the reduced resistance of the existing distal (sister) subtree is updated as

$$R_{sis} = \frac{8\mu}{\pi} (||\vec{x} - \vec{D}|| - ||\vec{D} - \vec{P}||) + R_{\downarrow}(\vec{D})$$

where $\vec{P}$ and $\vec{D}$ are the proximal and distal endpoint locations of the original parent segment, $\vec{T}$ is the sampled terminal target point location, and $R_{\downarrow}(\vec{D})$ is the previously stored downstream reduced resistance. The subtraction appears because the original parent segment is being replaced by a new bifurcation geometry.

The daughter radii are expressed as fractions of the parent radius

$$r_t = f_t r_{parent}, \qquad r_{sis} = f_{sis} r_{parent}$$

The split fractions are chosen to satisfy a Murray-style scaling,

$$f_t^{\gamma} + f_{sis}^{\gamma} = 1$$

with

$$\kappa = \left( \frac{Q_{sis} R_{sis}}{Q_t R_t} \right)^{\frac{\gamma}{4}}$$

$$f_t = (1 + \kappa)^{-\frac{1}{\gamma}}, \qquad f_{sis} = (1 + \kappa^{-1})^{-\frac{1}{\gamma}}$$

where $\gamma$ is the Murray exponent, commonly taken to be $\sim 3$. This comes from Murray's Law, which is described as the physiological principle of minimum work: increasing vessel radius lowers viscous pumping losses, but also increases the metabolic/volumetric cost of maintaining blood and vessel tissue [20]. The optimum balance between these opposing costs gives the branching rule

$$r_{\text{parent}}^{\gamma} = r_{\text{child1}}^{\gamma} + r_{\text{child2}}^{\gamma}$$

at each bifurcation (typically with $\gamma = 3$). The derivation for Murray's Law is shown in Appendix H.

The absolute radius scale is set by the root pressure-flow condition. If $R_0$ is the reduced equivalent of the updated tree,

$$R_0 = R_{\uparrow} + \left( \frac{f_t^4}{R_t} + \frac{f_{sis}^4}{R_{sis}} \right)^{-1}$$

then the root radius is

$$r_{root} = \left( \frac{Q_{\text{in}} R_0}{P_{\text{root}} - P_t} \right)^{\frac{1}{4}}$$

All downstream radii are then obtained by multiplying the root radius by the appropriate sequence of bifurcation ratios. In other words, the local bifurcation optimization changes the geometry and equivalent resistance, while the radius update enforces the prescribed flow delivery and pressure drop.

###### 3.1.3 Bifurcation optimization

For each sampled terminal target location $\vec{T}$, the algorithm searches over nearby potential parent segments. Suppose a candidate parent segment has a proximal endpoint location $\vec{P}$ and distal endpoint location $\vec{D}$. Inserting a new terminal requires replacing the original segment with three segments: $\vec{P} \rightarrow \vec{x}, \vec{x} \rightarrow \vec{D}$, and $\vec{x} \rightarrow \vec{T}$ where $\vec{x}$ is the new bifurcation point.

The bifurcation point is optimized within the triangular plane defined by $\vec{P}$, $\vec{D}$, and $\vec{T}$. We parameterize using barycentric coordinates,

$$x(s,t) = (1 - s - t)\vec{P} + s\vec{D} + t\vec{T}$$

with constraints $s \geq 0, t \geq 0, s+t \leq 1$. This maps the bounded two-variable optimization problem $(s,t)$ onto a physical point inside a 3D triangle spanned by the parent segment and the proposed terminal. Points with $t = 0$ lie directly on the original parent segment, while points with $t > 0$ allow the bifurcation to move off the parent segment within the local branching plane. For each candidate parent, the optimizer varies $(s,t)$, excludes locations which violate geometric constraints (e.g., out of tissue bounds, colliding with existing vessels), recomputes the resulting branch lengths, reduced resistances, Murray radius ratios, and tree cost, ultimately selecting the bifurcation location that minimizes the cost $J$ and committing it to the tree. This process is repeated until the desired number of terminal outlets has been reached.

The optimizer used is a Sequential Least Squares Programming (SLSQP) approach.

##### 3.2 Equal-bifurcation growth

For large trees ($\gtrsim$300,000 vessels for cube domains, $\gtrsim$700,000 for the heart), we switch from the full local CCO insertion step to a faster terminal-only equal-bifurcation mode. In this mode, new growth is not produced by sampling an arbitrary new terminal point $\vec{T}$ and optimizing a bifurcation point $\vec{x}$ on a nearby parent segment. Instead, the algorithm takes existing terminal segments and splits them into two daughter terminal segments with equal length and radius. Thus, if a terminal segment $e$ has proximal endpoint $\vec{P}_e$, distal endpoint $\vec{D}_e$, length $L_e$, and direction

$$\hat{w}_e = \frac{\vec{D}_e - \vec{P}_e}{||\vec{D}_e - \vec{P}_e||},$$

the new bifurcation is placed at the current terminal endpoint $\vec{D}_e$, and two new terminal endpoints are generated as

$$\vec{D}_{e,1} = \vec{D}_e + \ell_{\text{new}} \hat{d}_1, \qquad \vec{D}_{e,2} = \vec{D}_e + \ell_{\text{new}} \hat{d}_2.$$

The directions $\hat{d}_1$ and $\hat{d}_2$ are chosen symmetrically around the parent direction. If $\alpha$ is the daughter angle relative to $\hat{w}_e$ and $\hat{p}$ is a random transverse unit vector, then,

$$\hat{d}_1 = \hat{w}_e \cos\alpha + \hat{p} \sin\alpha, \qquad \hat{d}_2 = \hat{w}_e \cos\alpha - \hat{p} \sin\alpha.$$

The angle $\alpha$ is sampled from a triangular distribution.

The new daughter length $\ell_{\text{new}}$ is prescribed by a configured length law rather than by direct minimization of $J$.

$$\ell_{\text{new}} = c_\ell N_t^p,$$

where $N_t$ is the current terminal count and $p < 0$ causes new vessels to shorten as the tree becomes denser. $p$ was determined to be -0.33 from the scaling that naturally emerged from CCO growth.

The name "equal-bifurcation" refers to the local Murray split. The two new daughters are assigned equal radius ratios,

$$f_1 = f_2 = 2^{-1/\gamma}$$

so that

$$(f_1 r_p)^\gamma + (f_2 r_p)^\gamma = r_p^\gamma$$

or equivalently,

$$r_1 = r_2 = r_p 2^{-1/\gamma}$$

##### 3.3 Biventricular heart model

For the heart simulations, vascular trees were generated from a coronary "seed" structure rather than from an abstract root vessel alone. Initial left and right coronary artery "seed" graphs were created using a simple segmenting procedure on a 3D coronary artery model which extracts only centerlines and radii, estimating the arteries as straight vessel segments.

The "seed" tree's two arterial trees were then compiled into objects containing segment endpoints, connectivity, initial flow assignments, root and terminal pressures, reduced resistances, radii, Murray-scaling parameters, vessel maps, and spatial indices. Final terminal-count targets were allocated between the LCA and RCA trees according to their prescribed root-flow fractions, while individual growth points during the collision-aware phase were sampled in the myocardial tissue domain and assigned to the nearest active coronary tree. The CCO growth phase proceeded using the procedure described above. The full heart model took about 20 hours to generate.

#### 4 Zero-dimensional (0D) flow solver via directed graph

##### 4.1 Network definition and vessel conductances

To compute the flow and pressure distributions throughout large vascular structures, it would be computationally prohibitive to solve full three-dimensional fluid equations. Rather, we model the vasculature as a zero-dimensional (0D) hydraulic network, in which each vessel is represented as a directed edge $e$ connecting two pressure nodes $i$ and $j$.

Each vessel segment is assumed to have a parabolic velocity profile, consistent with Hagen-Poiseuille flow through a radially symmetric tube. This is a common assumption in vascular network modeling [20, 33, 44]. For a vessel segment $e$ of length $L_e$, radius $a_e$, and viscosity $\mu_e$, the hydraulic resistance is

$$R_e = \frac{8\mu_e L_e}{\pi a_e^4}$$

Equivalently, we define the hydraulic conductance $K_e = R_e^{-1}$. If edge $e$ connects node $i$ to node $j$, the flow through that vessel is $Q_e = K_e(P_i - P_j)$, where $P_i$ and $P_j$ are the nodal pressures. A positive value of $Q_e$ indicates flow in the assigned graph direction $i \longrightarrow j$.

The unknown pressures are obtained by enforcing a conservation of flow at each internal node, considering that each edge has a resistance to such flow. This is directly analogous to solving an electrical circuit (made entirely of interconnected resistors) with Kirchhoff's laws. Compactly, we solve

$$(B^T K B)P = s, \qquad Q = KBP$$

where $P \in \mathbb{R}^N$ is the vector of nodal pressures, $Q \in \mathbb{R}^E$ is the vector of edge flow rates, $K$ is the diagonal matrix of vessel conductances, $B \in R^{E \times N}$ is the edge-node incidence matrix, and $s \in R^N$ contains any prescribed source or sink flows. For an edge $e = (i, j)$,

$$B_{e,k} = \begin{cases} +1, & k = i \\ -1, & k = j \\ 0, & \text{otherwise} \end{cases} \qquad \text{for } e = (i, j)$$

This construction makes $BP$ the pressure drop across each vessel such that multiplying by $K$ results in the corresponding vessel flow ($Q = K\Delta P$).

#### 4.2 Boundary conditions

Our solver allows two hydraulic boundary-condition choices...

In the first, the total inlet flow $Q_{\text{in}}$ is prescribed at the root/inlet node, while the terminal pressure $P_{\text{term}}$ is prescribed at all outlet nodes. In this case, the solver determines the inlet pressure required to drive the specified total flow through the vascular network.

In the second, both the inlet pressure $P_{\text{in}}$ and terminal pressure $P_{\text{term}}$ are prescribed and the total inlet flow is determined by the network resistances.

Pressure boundary conditions are imposed by holding the corresponding node pressures fixed. Flow boundary conditions enter the Kirchhoff equations as a source at the root node $s_{\text{root}} = Q_{\text{in}}$. The remaining pressures and flows are solved from the conservation equations.

Figure S6 shows a schematic of this flow circuit to electrical circuit analogy.

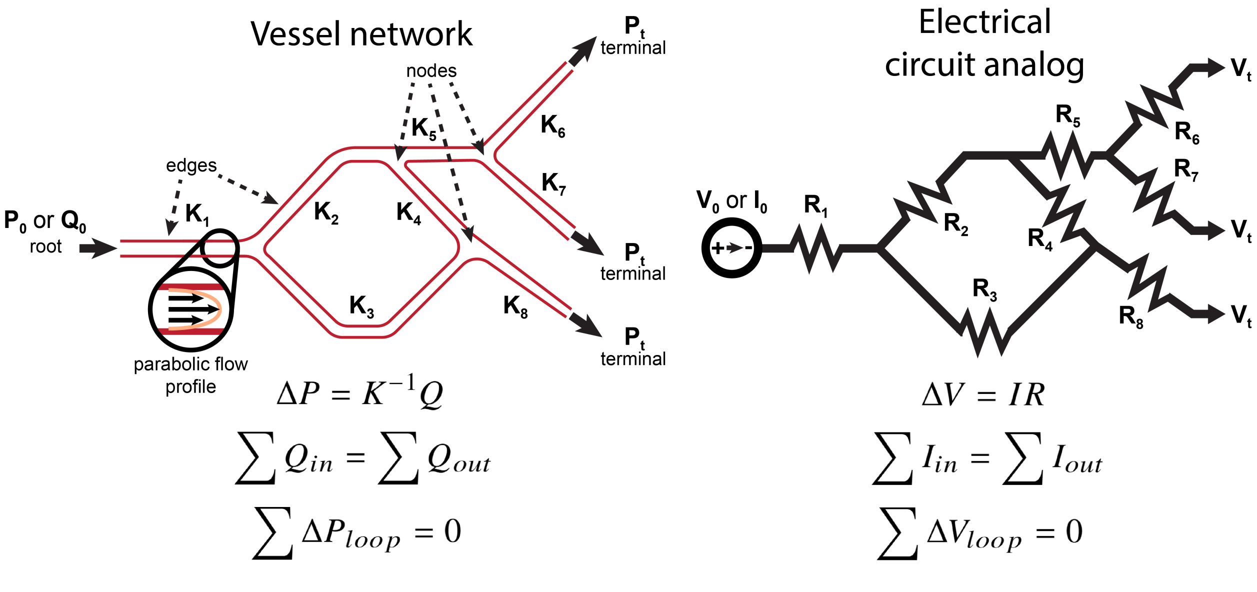

Figure S6: Schematic depicting the flow circuit to electrical circuit analogy for low-Reynolds number networks. Equations representing Ohm's law and Kirchhoff's laws are shown.

#### 4.3 Sparse graph solver for general networks

For a general vascular graph, including networks with loops or merged branches, the flow problem is solved directly as a sparse linear system. The sparse matrix is assembled by adding each vessel's conductance to the two connected node diagonals and subtracting it from the corresponding off-diagonal entries. For a vessel connecting nodes $i$ and $j$,

$$L_{ii} \leftarrow L_{ii} + K_e, \qquad L_{jj} \leftarrow L_{jj} + K_e,$$
$$L_{ij} \leftarrow L_{ij} - K_e, \qquad L_{ji} \leftarrow L_{ji} - K_e.$$

Therefore the conductance-weighted graph Laplacian is $L = B^T K B$. After pressure boundary conditions are applied, the reduced sparse system is solved. We allow several sparse linear solver options, including a direct sparse solve (spsolve), conjugate gradients (CG), and Generalized minimal residual method (GMRES) with incomplete-LU (ILU) preconditioning. The direct solver is robust for moderate network sizes, while the iterative solvers are useful for larger graphs where factorization becomes expensive. Vascular Laplacians, as we define here, can be very poorly conditioned (some very small and some very large eigenvalues); therefore, preconditioning is often necessary for the iterative methods.

#### 4.4 Tree-specific hydraulic solver

When the vascular graph is strictly a tree (no loops or merges, just bifurcations), we may compute the hydraulic solution more efficiently by taking advantage of the hierarchy of the tree. In this case, each downstream subtree can be replaced by a single effective conductance.

Starting from the terminal vessels and moving upward toward the root, the solver computes the equivalent conductance of each branch. A vessel with conductance $K_e$ feeding a downstream subtree with net conductance $K_{\text{down}}$ behaves like two hydraulic resistances in series,

$$K_{\text{branch}} = \left(\frac{1}{K_e} + \frac{1}{K_{\text{down}}}\right)^{-1}$$

At a bifurcation, daughter branches act in parallel, so their effective conductances add,

$$K_{\text{subtree}} = K_1 + K_2 + \ldots$$

This upward pass gives the total conductance $K_{\text{tree}}$ of the tree as seen from the inlet. If the inlet flow and terminal pressure are prescribed, the inlet pressure is

$$P_{\text{in}} = P_{\text{term}} + \frac{Q_{\text{in}}}{K_{\text{tree}}}$$

If the inlet and terminal pressures are prescribed, the inlet flow must be

$$Q_{\text{in}} = K_{\text{tree}}(P_{\text{in}} - P_{\text{term}})$$

Once the total conductance and root flow are known, a downward pass assigns flows and pressures to every vessel. At each bifurcation, the incoming flow is divided among daughter branches in proportion to their downstream conductances,

$$Q_k = Q_{\text{parent}} \frac{K_k}{\sum_m K_m}$$

and the pressure drop across each vessel is simply $\Delta P_e = \frac{Q_e}{K_e}$.

#### 4.5 Blood viscosity and hematocrit

For Newtonian fluids such as water or cell media, the viscosity $\mu_e$ is globally fixed, so each vessel conductance $K_e$ is fixed and the hydraulic problem is only solved once. For blood, however, the effective viscosity varies with vessel diameter and hematocrit due to non-Newtonian rheology and the Fåhræus–Lindqvist effect...

$$\frac{H_{T,i}}{H_D} = H_D + (1 - H_D)(1 + 1.7e^{-0.83a_i} - 0.6e^{-0.022a_i})$$

where $H_T$ and $H_D$ are the "tube" (vessel) and "discharge" (upstream/inlet) hematocrit values, and $a_i$ is the vessel radius in $\mu$m.

In practice, hematocrit distributions also depend on bifurcation angles, plasma skimming effects, and more. To compute hematocrit values and viscosities at each vessel, we employ the NetFlowV2 phase-separation model developed and maintained by Tim Secomb and others at the University of Arizona [27–33, 37, 39]. With this method, along with our flow solver, flow and hematocrit fields are updated iteratively until flows converge. The final flows and hematocrit values are fed as inputs to the following oxygen transport calculations.

#### 4.6 Resistance scaling

In the main text, we mention that the net resistance per unit length of a vascular tree decreases with tree size. Appendix I provides a potential explanation for this.

### 5 One-dimensional (1D) concentration solver via directed graph

#### 5.1 Graph representation of intravessel concentrations

Once the hydraulic solve, as defined by Section 4, is complete, each vessel segment has been assigned a flow rate $Q_e$. We use these flow rates to solve for the intravessel concentration field on the same directed graph used for the flow problem. The goal of this step is not yet to compute the full tissue oxygen field, but rather to determine how the concentration carried inside the vascular network changes as fluid moves from the inlet, through vessel segments, and across junctions.

The vascular network is represented as a graph with $N$ nodes and $E$ vessel edges. Each node $i$ is assigned a nodal concentration $C_i$ such that the full nodal concentration vector is denoted by $C \in \mathbb{R}^N$. For each vessel segment $e$, the upstream and downstream nodes are chosen according to the computed flow direction. If the hydraulic solve gives a negative flow relative to the original graph orientation, the edge is reversed for the concentration calculation. We then work only with the nonnegative flow magnitude $|Q_e|$.

For each edge $e$, we denote the upstream and downstream nodes by $\mathrm{up}(e)$ and $\mathrm{down}(e)$. By definition, the concentration entering each vessel is the concentration at its upstream node $C_{e,\mathrm{in}} = C_{\mathrm{up}(e)}$. Over a vessel segment, concentration changes due to exchange with the surrounding tissue and, for blood, due to buffering by hemoglobin. These details are defined in the oxygen transport sections that follow. At the graph-solver level, however, we only need the resulting one-dimensional edge update. With the local transport coefficients and external concentration field (outside the vessel) held fixed, this update can be written as

$$C_{e,\mathrm{out}} = \gamma_e C_{e,\mathrm{in}} + \chi_e$$

$\gamma_e$ is an edgewise attenuation factor and $\chi_e$ is the contribution from the external concentration field surrounding the vessel. If the external concentration field is approximated as constant over the vessel segment, then the edge update can be written more intuitively as a relaxation toward the local external concentration

$$C_{e,\mathrm{out}} = C_e^{\mathrm{ext}} + \gamma_e(C_{e,\mathrm{in}} - C_e^{\mathrm{ext}}) = \gamma_e C_{e,\mathrm{in}} + (1-\gamma_e)C_e^{\mathrm{ext}} = \gamma_e C_{e,\mathrm{in}} + \chi_e$$

where $\chi_e = (1-\gamma_e)C_e^{\mathrm{ext}}$. Essentially, $\gamma_e$ describes how strongly the vessel retains its upstream concentration, while $\chi_e$ describes relaxation toward the local external concentration. If $\gamma_e \approx 1$, the vessel concentration changes weakly over the segment. Smaller values mean the vessel more strongly equilibrates with the surrounding tissue.

The inlet concentration is prescribed at the root node $C_{\mathrm{root}} = C_{\mathrm{in}}$, but all other nodal concentrations are determined by mass conservation at graph junctions.

#### 5.2 Axial subdivision of vessel segments

In practice, a vessel segment does not necessarily see a constant external field $C_e^{\mathrm{ext}}$ nor have a constant attenuation factor $\gamma_e$ due to nonlinear kinetics. Therefore, it can be evaluated by using multiple shorter axial substeps. This allows the transport coefficients and external concentration to vary along the vessel while retaining the same edge-level update.

Suppose we divide vessel $e$ into substeps $m = 1, 2, ..., M_e - 1, M_e$. Over substep $m$, the local update is

$$C_{m+1} = \gamma_{e,m} C_m + (1-\gamma_{e,m})C_{e,m}^{\mathrm{ext}}$$

where $C_{e,m}^{\mathrm{ext}}$ is the local external concentration sampled along that specific portion of the vessel. Applying these substep updates sequentially gives a single effective edge update where

$$\gamma_e = \prod_{m=1}^{M_e} \gamma_{e,m}$$

The external field contribution is the accumulated contribution from every substep, with each contribution attenuated by the downstream substeps that follow it

$$\chi_e = \sum_{m=1}^{M_e} (1-\gamma_{e,m})C_{e,m}^{\mathrm{ext}} \prod_{\ell=m+1}^{M_e} \gamma_{e,\ell}$$

The empty product for the final substep is taken as one.

#### 5.3 Mass-conserving mixing at junctions

At each internal node, incoming streams mix to a common dissolved oxygen concentration $C_i$. For blood, conservation of total advected oxygen requires

$$\sum_{e\in I(i)} |Q_e| T_e(C_{e,\mathrm{out}}) = \sum_{e\in O(i)} |Q_e| T_e(C_i) \tag{1}$$

with total oxygen content $T_e(C) = C + C_{\mathrm{Hb}^{\max},e} S(C/\alpha)$ and hemoglobin oxygen capacity $C^{\max}_{\mathrm{Hb},e} = H_{D,e} C^{\max}_{\mathrm{RBC}}$ where $I(i)$ and $O(i)$ denote incoming and outgoing edges, $C_i$ is the mixed dissolved-oxygen concentration, $H_{D,e}$ is discharge hematocrit, $C^{\max}_{\mathrm{RBC}}$ is the fully saturated oxygen capacity per unit red-cell volume, $\alpha$ is the Henry's law coefficient, and $S(P)$ is the Severinghaus oxyhemoglobin-dissociation curve[41]. Without hemoglobin present, $T_e(C) = C$, and the balance reduces to

$$C_i \sum_{e\in O(i)} |Q_e| = \sum_{e\in I(i)} |Q_e| C_{e,\mathrm{out}} = \sum_{e\in I(i)} |Q_e| (\gamma_e C_{\mathrm{up}(e)} + \chi_e)$$

For ordinary internal nodes, the total inflow and outflow are equal. For terminal nodes, however, there are no outgoing vessels. To keep terminal concentrations well-defined we use the incoming flow as the terminal normalization.

$$w_i = \begin{cases} \sum_{e\in O(i)} |Q_e|, & \text{internal node} \\ \sum_{e\in I(i)} |Q_e|, & \text{terminal/outflow node} \end{cases}$$

Then the hemoglobin-free nodal balance can be compactly written as

$$w_i C_i - \sum_{e\in I(i)} |Q_e| \gamma_e C_{\mathrm{up}(e)} = \sum_{e\in I(i)} |Q_e| \chi_e$$

which separates the two parts of the graph concentration problem. For blood, the reported terminal concentration is obtained using the incoming-flow-weighted discharge hematocrit. Along each vessel, concentration is updated by one-dimensional transport and exchange with the external field. At each node, incoming streams mix by conservation of total advected oxygen.

#### 5.4 Full network concentration solve

For general vascular graphs, including graphs with loops or merging upstream paths, the concentration field is determined by edgewise transport and the nodal balances explained in the previous section. The coupled solver evaluates these balances by directed propagation, with under-relaxed iterative nodal updates.

Before assembling the system, every vessel is oriented in the direction of its computed flow. Each edge therefore has an upstream node, a downstream node, and a nonnegative flow magnitude $|Q_e|$. During a single linear solve, the edge coefficients $\gamma_e$ and $\chi_e$ are held fixed. The coefficients may depend on the current vessel concentrations and the current external concentration field, but are frozen while the graph system itself is solved.

We define selector matrices $S_{\mathrm{up}}$ and $S_{\mathrm{down}}$ such that

$$(S_{\mathrm{up}} C)_e = C_{\mathrm{up}(e)}$$

and

$$(S_{\mathrm{down}} C)_e = C_{\mathrm{down}(e)}$$

Let $Q$, $\gamma$, and $\chi$ denote the edgewise vectors of flow magnitudes, attenuation factors, and external-field contributions. Furthermore, let $W = \mathrm{diag}(w_i)$ be the diagonal matrix of nodal flow weights defined earlier. Instead of solving directly from iteration $k$ to the next iteration $k+1$ ($C^{(k)} \to C^{(k+1)}$), we perform a Picard iteration such that we first form an initial solution $C^{(k)} \to C^{\dagger}$ then relax that solution $C^{\dagger} \to C^{(k+1)}$. Without hemoglobin, the full graph system at iteration $k$ is

$$A^{(k)} C^{\dagger} = b^{(k)},$$

where

$$A^{(k)} = W - S^T_{\mathrm{down}} \mathrm{diag}(|\mathrm{Q_e}| \odot \gamma_{\mathrm{e}}^{(\mathrm{k})}) \mathrm{S_{up}},$$

and

$$b = S^T_{\mathrm{down}} (|Q_e| \odot \chi_e^{(k)}).$$

Here, $\odot$ denotes elementwise multiplication and $C^{\dagger}$ denotes the provisional nodal concentration vector obtained before under-relaxation (an intermediate solve). This is just the matrix form of the nodal balance defined in the previous subsection. Equivalently, at each node $i$ at system iteration $k$,

$$w_i C_i^{\dagger} - \sum_{e \in \mathcal{I}(i)} |Q_e| \gamma_e^{(k)} C_{\mathrm{up}(e)}^{\dagger} = \sum_{e \in \mathcal{I}(i)} |Q_e| \chi_e^{(k)},$$

before imposing boundary conditions.

For blood, total oxygen content is linearized separately at incoming outlet states and outgoing nodal states. Define the outgoing flow-weighted content relation

$$\hat{T}_i(C) = \frac{1}{w_i} \sum_{e \in O(i)} |Q_e| T_e(C)$$

for internal nodes. At terminal nodes, use

$$\hat{T}_i(C) = C + \overline{H}_{D,i} C_{\mathrm{RBC}}^{\max} S(C/\alpha)$$

where $\overline{H}_{D,i}$ is the flow-weighted discharge hematocrit. The resulting linearized nodal balance, after affine approximations, at iteration $k$ is

$$w_i \mathcal{B}_i^{\mathrm{out},(k)} C_i^{\dagger} - \sum_{e \in \mathcal{I}(i)} |Q_e| \mathcal{B}_e^{\mathrm{in},(k)} \gamma_e^{(k)} C_{\mathrm{up}(e)}^{\dagger} = \sum_{e \in \mathcal{I}(i)} |Q_e| \left( \mathcal{B}_e^{\mathrm{in},(k)} \chi_e^{(k)} + d_e^{\mathrm{in},(k)} \right) - w_i d_i^{\mathrm{out},(k)}$$

Here $\mathcal{B}^{\mathrm{in},(k)}$ and $\mathcal{B}^{\mathrm{out},(k)}$ are the derivatives of $T_e$ and $\hat{T}_i$, evaluated at $C_{e,\mathrm{out}}^{(k)}$ and $C_i^{(k)}$, respectively. The corresponding affine intercepts are $d = T - \mathcal{B}C$, evaluated at the same states. In matrix form,

$$A^{(k)} = W \mathrm{diag}(\mathcal{B}^{\mathrm{out},(k)}) - S_{\mathrm{down}}^T \mathrm{diag}(Q \odot \mathcal{B}^{\mathrm{in},(k)} \odot \gamma^{(k)}) S_{\mathrm{up}},$$

$$b^{(k)} = S_{\mathrm{down}}^T [Q \odot (\mathcal{B}^{\mathrm{in},(k)} \odot \chi^{(k)} + d^{\mathrm{in},(k)})] - W d^{\mathrm{out},(k)}.$$

These expressions recover the hemoglobin-free system when $\mathcal{B} = 1$ and $d = 0$.

For either system, we express the linear solve using block elimination. If we denote $d$ as the Dirichlet root (inlet) nodes and $f$ as the remaining free nodes (non-inlet nodes, including terminals), we may split $C$ and $A$ into

$$C = \begin{bmatrix} C_f \\ C_d \end{bmatrix}, \qquad A = \begin{bmatrix} A_{ff} & A_{fd} \\ A_{df} & A_{dd} \end{bmatrix}$$

so that

$$\begin{bmatrix} A_{ff} & A_{fd} \\ A_{df} & A_{dd} \end{bmatrix} \begin{bmatrix} C_f^{\dagger} \\ C_d^{\dagger} \end{bmatrix} = \begin{bmatrix} b_f \\ b_d \end{bmatrix}$$

We impose Dirichlet boundary conditions on the root (inlet) nodes $C_d^{\dagger} = C_{\mathrm{in}}$. The corresponding boundary rows are replaced by $A_{df} = 0$, $A_{dd} = I$, and $b_d = C_{\mathrm{in}}$. Ultimately we solve the reduced system

$$A_{ff}^{(k)} C_f^{\dagger} = b_f^{(k)} - A_{fd}^{(k)} C_d^{\dagger}, \qquad C_d^{\dagger} = C_{\mathrm{in}} \text{ for } n \in \text{root nodes}$$

After this linear solve, we under-relax using $dC^{(k)} = C^{\dagger} - C^{(k)}$

$$C^{(k+1)} = C^{(k)} + \omega dC^{(k)} = (1 - \omega) C^{(k)} + \omega C^{\dagger}$$

where $0 < \omega \leq 1$. We repeat the whole process until $||dC^{(k)}|| < \epsilon_{\mathrm{convergence}}$.

The external concentration field is also updated at each iteration and depends on $C^{(k)}$. Details of the external field are explained in the following sections.

##### 5.5 Top-down concentration solve for tree networks

When the vascular graph is a bifurcating tree (or any other structure with no loops or merges), the concentration field can be computed more efficiently by exploiting the upstream path from the root to every vessel. In this case, no global concentration matrix is required. The solver instead advances concentrations through the tree in a marching proximal-to-distal order. This top-down solver gives the same concentration solution as the sparse graph solve, but avoids assembling and solving a global linear system. This solver is available but was not used for the studies in the main text.

#### 6 Oxygen transport theory

The following is the primary contribution of this work: a Yukawa-Green's function based solver which resolves both intravessel and intratissue concentration distributions.

##### 6.1 Setting up the steady-state equations

To solve for the oxygen field surrounding a set of vessels, we define each vessel (or subsection thereof) as a straight cylinder. About each of these vessels, we analyze the radial, cylindrical reaction-diffusion problem

$$\frac{D}{r}\frac{d}{dr}\left(r\frac{dC}{dr}\right) = \frac{V_{\max}C}{K_M + C}$$

where we assume axial diffusion is negligible in comparison to radial diffusion (see section F for an analysis on this). Note that we assume that the permeability of oxygen through the vessel wall is roughly the same as the surrounding tissue. This assumption is reasonable for small molecules but is not valid for large molecules such as proteins, which typically require facilitated mass transfer. We define a characteristic reaction-diffusion length scale $\lambda_i$ and a nondimensional radius $\phi_i$ for each vessel $i$.

$$\lambda_i = \sqrt{\frac{D(K_M + C_i^{\text{eff}})}{V_{\max}}} \qquad \phi_i = \frac{a_i}{\lambda_i}$$

More information on $\lambda_i$ and $C_i^{\text{eff}}$ can be found in section B. Using this length scale, we linearize the reaction term by freezing the denominator of the Michaelis-Menten uptake term locally as

$$\frac{V_{\max}C}{K_M + C} \approx \frac{V_{\max}C}{K_M + C_i^{\text{eff}}} = D\lambda_i^{-2}C$$

Near vessel $i$, we decompose the tissue concentration into a smooth regular field $C_i^{\text{reg}}(\vec{x})$ and a local radial disturbance $c_i(r, s)$,

$$C_i(\vec{x}) = C_i^{\text{reg}}(\vec{x}) + c_i(r, s)$$

Here $C_i^{\text{reg}}(\vec{x})$ represents the regular background field generated by all other vessels after removing the local, singular contribution of vessel $i$. Because this regular field should vary slowly over the vessel radius, we approximate its local value near the vessel wall by $C_i^{\text{reg}}(\vec{x}) \approx C_i^{\text{ext}}(s)$ where $C_i^{\text{ext}}(s)$ is the external concentration experienced by vessel $i$ at axial position $s$. Thus, the near-vessel concentration is locally approximated as

$$C_i(\vec{x}) \approx C_i^{\text{ext}}(s) + c_i(r, s)$$

The local disturbance $c_i(r, s)$ is the radially varying correction needed to match the channel concentration to this smooth external field. Under the locally linearized, frozen-coefficient approximation, $c_i$ satisfies

$$D\left(\frac{d^2c_i}{dr^2} + \frac{1}{r}\frac{dc_i}{dr}\right) - D\lambda_i^{-2}c_i = 0$$

We assume this operator is also locally satisfied for $C_i^{\text{ext}}(s)$. Now to solve the equation... we assume (relatively) fast radial diffusion inside the channels and we also prescribe a far-field matching condition on the disturbance.

$$C_i(a_i, s) = C_{w,i}(s) \quad \rightarrow \quad c_i(a_i, s) = C_{w,i}(s) - C_i^{\text{ext}}(s)$$

$$c_i(r \to \infty, s) \to 0$$

where $C_{w,i}(s)$ is the channel wall oxygen concentration along its axial position $s$ at $r = a_i$. Note that we are assuming that dissolved oxygen content and hemoglobin-bound oxygen content are in instantaneous equilibrium. From this, we find a solution in terms of the decaying modified cylindrical Bessel functions

$$c_i(r, s) = (C_{w,i}(s) - C_i^{\text{ext}}(s)) \frac{K_0\left(\frac{r}{\lambda_i(s)}\right)}{K_0(\phi_i(s))}$$

The resulting outward flux per unit length of the channel is then defined as

$$q_i(s) = 2\pi a_i J_n = 2\pi a_i \left( -D \frac{\partial c_i}{\partial r}\bigg|_{r=a_i} \right) = \kappa_i(s)(C_{w,i}(s) - C_i^{\text{ext}}(s))$$

with an effective interfacial mass transfer coefficient

$$\kappa_i(s) = 2\pi a_i \frac{D}{\lambda_i} \frac{K_1(\phi_i(s))}{K_0(\phi_i(s))}$$

#### 6.2 Solving for intravessel oxygen distributions

##### 6.2.1 Hemoglobin buffering and depletion/growth along the vessel

We note that this outward flux must cause a reduction of the bulk intravessel oxygen concentration $C_b(s)$ along $s$, or rather an increase if the surrounding external concentration is higher than the wall concentration $C_i^{\text{ext}}(s) > C_{w,i}(s)$.

We denote the bulk vessel concentration of vessel $i$ by $C_{b,i}(s)$ and the vessel wall concentration by $C_{w,i}(s)$. These two quantities are approximately equal when the lumen-tissue Biot number $\text{Bi}_i$ is small. More information on the closure between $C_{w,i}$ and $C_{b,i}$ is present in the following section. In the general case, $C_{w,i}$ determines the wall flux, while $C_{b,i}$ determines the oxygen content advected through the vessel.

From the tissue-side radial shell problem in Section 6.1, the outward flux per unit length for a hemoglobin-free perfusate is $q$, which appears in the axial balance

$$Q_i \frac{d(C_{b,i}(s))}{ds} = -q_i(s) = -\kappa_i(C_{w,i}(s) - C_i^{\text{ext}}(s))$$

However, for blood, hemoglobin will offload oxygen when concentration drops and will bind oxygen when dissolved concentration increases, buffering the rate of change of the bulk dissolved oxygen concentration. We assume that hemoglobin-bound oxygen and dissolved oxygen remain in instantaneous equilibrium. Let the total oxygen concentration be defined by

$$C_{\text{total}} = T(C_b) = C_b + C_{\text{Hb}} = \alpha P + C_{\text{Hb}}^{\text{max}} S(P_{O_2})$$

where $P$ is the partial pressure of free oxygen in blood ($C_b = \alpha P$), $\alpha$ is the oxygen solubility, $C_{\text{Hb}}^{\text{max}}$ is the hemoglobin oxygen capacity, and $S(P)$ is the hemoglobin saturation function (taken here as the Severinghaus oxyhemoglobin dissociation curve [41]). $C_{\text{Hb}}^{\text{max}} = H_D C_{\text{RBC}}^{\text{max}}$ where $H_D$ is discharge hematocrit and $C_{\text{RBC}}^{\text{max}}$ is the hemoglobin-bound oxygen capacity per unit volume of red blood cells.

$$S(P) = \frac{P^3 + 150P}{P^3 + 150P + 23400}$$

...where $P$ is oxygen partial pressure in mmHg.

Using the chain rule,

$$\frac{dC_{\text{total}}(s)}{ds} = \frac{d(C_b + C_{\text{Hb}})}{ds} = \frac{dC_{\text{total}}}{dP}\frac{dP}{ds} = \left(\alpha + C_{\text{Hb}}^{\text{max}} \frac{dS}{dP}\right)\frac{dP}{ds} = \frac{\beta(P)}{\alpha}\frac{dC_b}{ds}$$

Therefore, for vessel $i$,

$$Q_i \frac{dC_{\text{total}}(C_{b,i})}{ds} = -q_i(s)$$

so that

$$\frac{dC_{b,i}}{ds} = -\frac{\alpha}{\beta_i}\frac{\kappa_i(s)}{Q_i}(C_{w,i}(s) - C_i^{\text{ext}}(s)) = -\frac{\tilde{\kappa}_i}{Q_i}(C_{w,i}(s) - C_i^{\text{ext}}(s))$$

This is the hemoglobin-buffered axial balance. Here $\beta_i = \beta(\frac{C_{b,i}}{\alpha})$ (which may be frozen over a short axial interval). $\tilde{\kappa}$ is the buffered interfacial mass transfer coefficient such that $\tilde{\kappa} = \frac{\alpha}{\beta_i}\kappa_i$.

For segments evaluated on short enough intervals where $\tilde{\kappa}_i$ can be frozen, the well-mixed solution (small Biot number) is

$$C_{b,i}(s) = e^{-\frac{\tilde{\kappa}_i}{Q_i}s}\left(C_{b,i}(0) + \frac{\tilde{\kappa}_i}{Q_i}\int_0^s C_i^{\text{ext}}(\sigma)e^{\frac{\tilde{\kappa}_i}{Q_i}\sigma}d\sigma\right)$$

where the wall concentration would match the bulk concentration

$$C_{w,i} = C_{b,i}$$

Assuming a relatively constant external concentration along each axial vessel subdivision, the channel concentration changes according to

$$C_{b,i}(s) = C_i^{\text{ext}} + (C_{b,i}(0) - C_i^{\text{ext}})e^{-\frac{\tilde{\kappa}_i}{Q_i}s}$$

This exponential expression should be interpreted as the well-mixed-lumen limit. When finite radial diffusion inside the vessel is resolved, $C_{w,i} \neq C_{b,i}$, and the wall concentration is instead obtained from the Graetz closure to follow.

##### 6.2.2 Closure for radial oxygen profiles inside the channel

To resolve the finite radial diffusion inside the vessel, we introduce the intraluminal dissolved oxygen concentration inside the channel

$$C_{c,i}(r, s)$$

where at the wall

$$C_{w,i}(s) = C_{c,i}(a_i, s)$$

The bulk concentration is

$$C_{b,i}(s) = \frac{1}{Q_i}\int_0^{a_i} 2\pi r u_i(r) C_{c,i}(r, s)dr$$

For a straight vessel segment with constant radius $a_i$, the mean velocity is $U_i = \frac{Q_i}{\pi a_i^2}$. With $\tilde{r} = \frac{r}{a_i}$, the normalized velocity profile is

$$v_i(\tilde{r}) = \frac{u_i(r)}{U_i}$$

such that

$$2\int_0^1 \tilde{r} v_i(\tilde{r})d\tilde{r} = 1$$

The high-Péclet number (assuming axial diffusion is negligible compared to advection) intraluminal convection-diffusion problem is

$$\mathcal{B}_i u_i(r)\frac{\partial C_{c,i}}{\partial s} = D_v \frac{1}{r}\frac{\partial}{\partial r}\left(r\frac{\partial C_{c,i}}{\partial r}\right)$$

where $D_v$ is the intraluminal diffusivity of oxygen. For water, plasma, or cell media, $\mathcal{B}_i = 1$. For blood, the frozen buffer factor is $\mathcal{B}_i = \frac{\beta_i}{\alpha}$. We impose a centerline symmetry condition

$$\frac{\partial C_{c,i}}{\partial r}(0, s) = 0$$

and a flux continuity condition

$$-D_v\frac{\partial C_{c,i}}{\partial r}(a_i, s) = h_i(s)(C_{c,i}(a_i, s) - C_i^{\text{ext}}(s))$$

where $h_i(s) = \frac{\kappa_i(s)}{2\pi a_i}$. Thus it must be true that $C_{c,i}(a_i, s) = C_{w,i}(s)$.

For Poiseuille flow (parabolic flow profile),

$$v_i(\tilde{r}) = 2(1 - \tilde{r}^2)$$

Over a short axial interval, we freeze $C_i^{\text{ext}}$, $h_i$, $\kappa_i$, $\lambda_i$, $U_i$, $a_i$, and $\mathcal{B}_i$.

We define nondimensional lumen concentration $\Theta_i(\tilde{r}, \tilde{s})$, external concentration $\Theta_i^{\text{ext}}$, concentration difference $\tilde{\Theta}$, and axial coordinate $\tilde{s}$,

$$\Theta_i(\tilde{r}, \tilde{s}) = \frac{C_{c,i}(\tilde{r}, \tilde{s})}{C_{b,i}^{\text{in}}}, \qquad \Theta_i^{\text{ext}} = \frac{C_i^{\text{ext}}}{C_{b,i}^{\text{in}}}, \qquad \tilde{\Theta}_i(\tilde{r}, \tilde{s}) = \Theta_i(\tilde{r}, \tilde{s}) - \Theta_i^{\text{ext}}, \qquad \tilde{s} = \frac{D_v}{\mathcal{B}_i a_i^2 U_i} s$$

Where $C_{b,i}^{\text{in}}$ is the inlet bulk concentration to that vessel subdivision. We also define the lumen-tissue Biot number

$$\text{Bi}_i = \frac{a_i h_i}{D_v} = \frac{\kappa_i}{2\pi D_v}$$

which measures the strength of the tissue-side wall extraction relative to intraluminal radial diffusion. When the Biot number is small $\text{Bi}_i \ll 1$, the radial profile remains nearly uniform and $C_{w,i} \approx C_{b,i}$.

With these definitions, the intraluminal problem becomes

$$v_i(\tilde{r}) \frac{\partial \tilde{\Theta}_i}{\partial \tilde{s}} = \frac{1}{\tilde{r}} \frac{\partial}{\partial \tilde{r}} \left( \tilde{r} \frac{\partial \tilde{\Theta}_i}{\partial \tilde{r}} \right)$$

with centerline symmetry

$$\frac{\partial \tilde{\Theta}_i}{\partial \tilde{r}}(0, \tilde{s}) = 0$$

and wall condition

$$\frac{\partial \tilde{\Theta}_i}{\partial \tilde{r}}(1, \tilde{s}) = -\text{Bi}_i \tilde{\Theta}_i(1, \tilde{s})$$

We seek separated solutions of the form

$$\tilde{\Theta}_i(\tilde{r}, \tilde{s}) = \phi_{n,i}(\tilde{r}) e^{-\mu_{n,i}^2 \tilde{s}}$$

The radial eigenfunctions satisfy the general Graetz eigenproblem

$$\frac{1}{\tilde{r}} \frac{d}{d\tilde{r}} \left( \tilde{r} \frac{d\phi_{n,i}}{d\tilde{r}} \right) = -\mu_{n,i}^2 v_i(\tilde{r}) \phi_{n,i}$$

with $\phi'_{n,i}(0) = 0$ and $-\phi'_{n,i}(1) = \text{Bi}_i \phi_{n,i}(1)$. The eigenfunctions must be orthogonal under the velocity-weighted inner product

$$\langle f, g \rangle = \int_0^1 \tilde{r} v_i(\tilde{r}) f(\tilde{r}) g(\tilde{r}) d\tilde{r}$$

If $\tilde{\Theta}_{i,0}(\tilde{r})$ is the incoming dimensionless excess concentration profile at the beginning of the frozen axial interval of vessel $i$, then

$$\tilde{\Theta}_{i,0}(\tilde{r}) = \sum_n A_{n,i} \phi_{n,i}(\tilde{r}), \qquad A_{n,i} = \frac{\langle \tilde{\Theta}_{i,0}, \phi_{n,i} \rangle}{\langle \phi_{n,i}, \phi_{n,i} \rangle}$$

The resulting solution after relative axial distance $\tilde{s}$ is

$$\tilde{\Theta}_i(\tilde{r}, \tilde{s}) = \sum_n A_{n,i} \phi_{n,i} e^{-\mu_{n,i}^2 \tilde{s}}$$

We may then recover the dimensional wall concentration...

$$C_{w,i}(s) = C_{b,i}^{\text{in}} \Theta_i(1, \tilde{s}) = C_i^{\text{ext}} + C_{b,i}^{\text{in}} \sum_n A_{n,i} \phi_{n,i}(1) e^{-\mu_{n,i}^2 \frac{D_v}{\mathcal{B}_i a_i^2 U_i} s}$$

...the bulk concentration...

$$C_{b,i}(s) = 2C_{b,i}^{\text{in}} \int_0^1 \tilde{r} v_i(\tilde{r}) \Theta_i(\tilde{r}, \tilde{s}) d\tilde{r} = C_i^{\text{ext}} + 2C_{b,i}^{\text{in}} \int_0^1 \left( \tilde{r} v_i(\tilde{r}) \sum_n A_{n,i} \phi_{n,i}(\tilde{r}) e^{-\mu_{n,i}^2 \frac{D_v}{\mathcal{B}_i a_i^2 U_i} s} \right) d\tilde{r}$$

...and wall flux per unit length (for the disturbance)

$$q_i(s) = \kappa_i(s)(C_{w,i} - C_i^{\text{ext}}) = \kappa_i C_{b,i}^{\text{in}} \sum_n A_{n,i} \phi_{n,i}(1) e^{-\mu_{n,i}^2 \frac{D_v}{\mathcal{B}_i a_i^2 U_i} s}$$

In practice, because $\lambda_i$, $\kappa_i$, $h_i$, and $\text{Bi}_i$ all depend on the local wall concentration, this closure is solved by a simple fixed-point iteration over each frozen axial interval.

Figure S7 depicts the intraluminal oxygen profiles within a single vessel for $C_{\text{in}} > C^{\text{ext}}$ and $C_{\text{in}} < C^{\text{ext}}$ with flow from left to right.

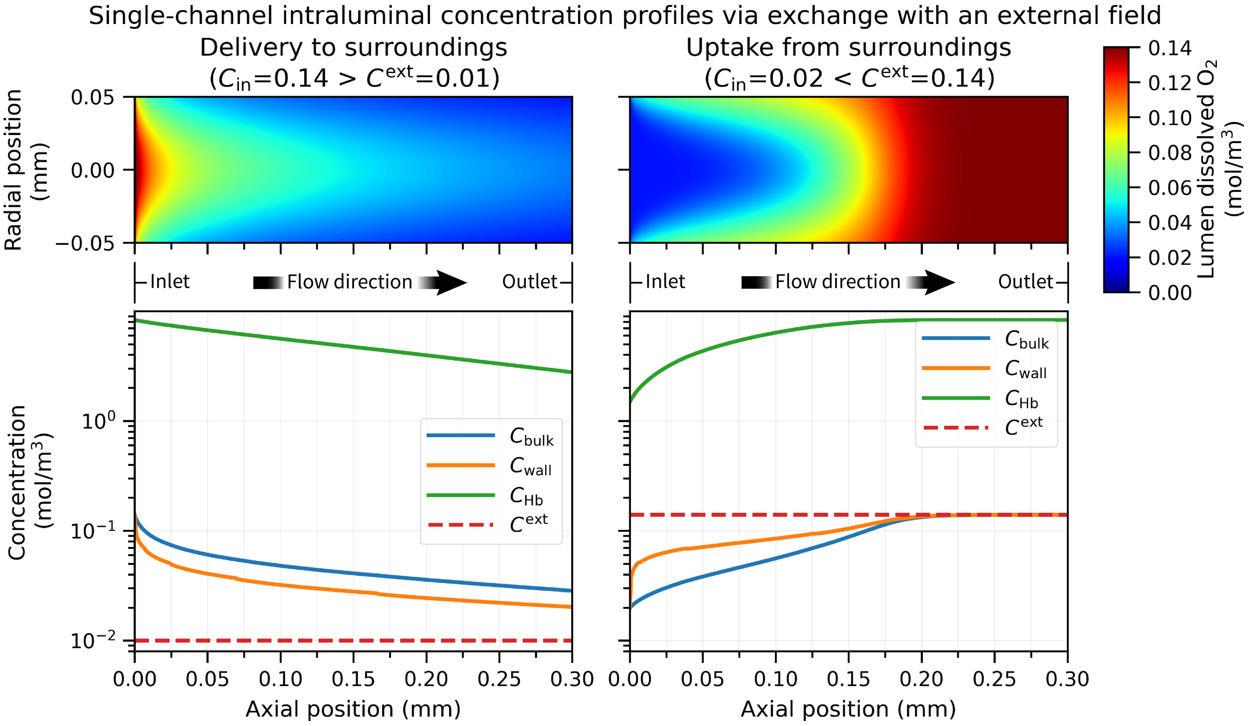

Figure S7: Representative intraluminal oxygen profiles within a single vessel acted upon by an external field either lower than the inlet concentration (left) or greater than the inlet concentration (right). Top plots show heatmaps of intraluminal concentration, bottom plots show bulk concentration ($C_{\text{bulk}}$), vessel wall concentration ($C_{\text{wall}}$), bulk hemoglobin-bound concentration ($C_{\text{Hb}}$), and prescribed external concentration $C^{\text{ext}}$ as a function of axial position. Flow runs from left to right. Bulk concentration $C_{\text{b}}$ is prescribed at the inlet. For both examples, $V_{\max} = 0.044 \frac{mol}{m^3 s}$, $K_M = 0.007 \frac{mol}{m^3}$, $Q = 0.0015 \frac{\mu L}{min}$. This flow rate is purposefully very low for visualization purposes.

#### 6.3 Solving for the tissue oxygen field

##### 6.3.1 Setting up the field equations

Treating the outward fluxes of each vessel as Yukawa line-sources

$$D\nabla^2 C - D\lambda^{-2} C = -S(\vec{x}), \qquad S(\vec{x}) = \sum_i \int_0^{L_i} q_i(s) \delta(\vec{x} - \vec{X}_i(s)) ds$$

we can define a solution in terms of Green's functions which satisfy

$$D(\nabla^2 - \lambda^{-2})G(r) = -\delta(\vec{x} - \vec{x}')$$

such that the source location is defined by the centerline of the source vessel $\vec{x}' = \vec{X}_i$. The Green's kernel is

$$G_i(r) = \frac{e^{-\frac{r_i}{\lambda_i}}}{4\pi r_i D}, \quad r_i = ||\vec{x} - \vec{X}_i(s)||$$

Using Green's second identity over the tissue domain $\Omega_t$ (Representation theorem)...

$$C(\vec{x}) = D \int_{\partial\Omega_t} (G(\nabla C \cdot \vec{n}) - C(\nabla G \cdot \vec{n}))\, dA'$$

where $G$ is $G(\vec{x}, \vec{x}')$, $C$ is $C(\vec{x}')$ on $\partial\Omega_t$, and $\vec{n}$ is the tissue-to-vessel normal vector (pointing into the vessel). The concentration at a vessel wall is $C_i(\vec{x}'(s,\theta)) \approx C_{w,i}(s)$.

##### 6.3.2 The monopole contribution

The first term in the $C(\vec{x})$ integral represents the flux/source monopole contribution. Defining the outward oxygen flux into the tissue as $J_n(\vec{x}') = D\nabla C(\vec{x}') \cdot \vec{n}$ so that $J_n > 0$ corresponds to oxygen entering the tissue,

$$C_{\text{monopole}}(\vec{x}) = D \int_{\partial\Omega_t} G(\nabla C \cdot \vec{n}) dA' = \int_{\partial\Omega_t} G J_n dA'$$

For vessel $i$, we parameterize the vessel wall as $\vec{x}_i'(s,\theta) = \vec{X}_i(s) - a_i \vec{n}_i(s,\theta)$ where $\vec{X}_i(s)$ is the vessel centerline oriented in the flow direction, $a_i$ is the vessel radius, and $\vec{n}_i(s,\theta)$ is the tissue-to-vessel normal (pointing into the vessel).

Assuming a relatively slender cylinder, $dA' \approx a_i d\theta ds$, and assuming the wall flux is approximately uniform around the vessel circumference, $J_{n,i}(s,\theta) \approx J_{n,i}(s)$. We define the total flux per unit length from vessel $i$ as

$$q_i(s) = \int_0^{2\pi} J_{n,i}(s) a_i d\theta = 2\pi a_i J_{n,i}(s)$$

Thus, the monopole contribution from vessel $i$ is

$$C_{\text{monopole},i}(\vec{x}) = \int_0^{L_i} \int_0^{2\pi} a_i J_{n,i}(s) G(\vec{x}, \vec{X}_i(s) - a_i \vec{n}_i(s,\theta)) d\theta ds$$

To obtain the finite-radius expansion, we then expand the Green's function about the centerline point $\vec{X}_i(s)$. Let $\nabla$ represent differentiation with respect to the source coordinate.

$$G(\vec{x}, \vec{X}_i - a_i\vec{n}) = G(\vec{x}, \vec{X}_i) - a_i \vec{n}_i \cdot \nabla G(\vec{x}, \vec{X}_i) + \frac{a_i^2}{2}(\vec{n}_i \otimes \vec{n}_i) : \nabla\nabla G(\vec{x}, \vec{X}_i) + O(a^3)$$

Substituting into the surface integral,

$$C_{\text{monopole},i}(\vec{x}) = \int_0^{L_i} a_i J_{n,i}(s) \int_0^{2\pi} \left( G(\vec{x}, \vec{X}_i) - a_i \vec{n}_i \cdot \nabla G(\vec{x}, \vec{X}_i) + \frac{a_i^2}{2}(\vec{n}_i \otimes \vec{n}_i) : \nabla\nabla G(\vec{x}, \vec{X}_i) + O(a^3) \right) d\theta ds$$

Inserting $q_i(s)$,

$$C_{\text{monopole},i}(\vec{x}) = \int_0^{L_i} q_i(s) G(\vec{x}, \vec{X}_i(s)) ds - \int_0^{L_i} \frac{a_i q_i(s)}{2\pi} \left[ \int_0^{2\pi} \vec{n}_i d\theta \right] \cdot \nabla G(\vec{x}, \vec{X}_i(s)) ds$$
$$+ \int_0^{L_i} \frac{a_i^2 q_i(s)}{4\pi} \left[ \int_0^{2\pi} \vec{n}_i \otimes \vec{n}_i d\theta \right] : \nabla\nabla G(\vec{x}, \vec{X}_i(s)) ds + ...$$

we see three terms (to order $O(a^2)$)

$$C_{\text{monopole},i}(\vec{x}) = C^{(0)}_{\text{monopole},i}(\vec{x}) + C^{(1)}_{\text{monopole},i}(\vec{x}) + C^{(2)}_{\text{monopole},i}(\vec{x})$$

The leading order term (expressed below as a total contribution)

$$C^{(0)}_{\text{monopole}}(\vec{x}) = \sum_i \int_0^{L_i} q_i(s) G(\vec{x}, \vec{X}_i(s)) ds$$

will make the largest contribution.

Note that for a circular cross-section, $\int_0^{2\pi} \vec{n}_i d\theta = 0$, so the first-order correction $C^{(1)}_{\text{monopole},i}$ is zero.

Furthermore, the circular cross-section yields $\int_0^{2\pi} \vec{n}_i \otimes \vec{n}_i d\theta = \pi p_i$ where $p_i$ is the projection tensor onto the plane normal to the local vessel tangent vector $\vec{t}_i(s)$.

$$p_i = I - \vec{t}_i \otimes \vec{t}_i$$

Here each vessel is a straight channel such that the tangent vector is a constant along the vessel arc length $\vec{t}_i(s) = \vec{t}_i$.

$$\vec{t}_i = \frac{\vec{X}_i(L_i) - \vec{X}_i(0)}{||\vec{X}_i(L_i) - \vec{X}_i(0)||}$$

Noting that $p_i : \nabla\nabla G = \nabla^2 G - \vec{t}_i \cdot (\nabla\nabla G) \cdot \vec{t}_i$, we may write the total second-order monopole correction at point $\vec{x}$ as

$$C^{(2)}_{\text{monopole}}(\vec{x}) = \sum_i \int_0^{L_i} \frac{a_i^2 q_i(s)}{4} \left( \nabla^2 G(\vec{x}, \vec{X}_i(s)) - \vec{t}_i \cdot (\nabla\nabla G(\vec{x}, \vec{X}_i(s))) \cdot \vec{t}_i \right) ds$$

For the Yukawa Green's function, $G_i = G(\vec{x}, \vec{X}_i(s)) = \frac{e^{-\frac{r_i}{\lambda_i}}}{4\pi D r_i}$ where $r_i = ||\vec{x} - \vec{X}_i(s)||$. With $\lambda_i$ held fixed locally, we may define the unit orientation vector,

$$\hat{r}_i = \frac{\vec{x} - \vec{X}_i(s)}{r_i}$$

Then evaluate the operators and resulting contractions as...

$$\nabla^2 G_i = \frac{1}{\lambda_i^2} G_i$$

$$\nabla\nabla G_i = \left[ \left( \frac{3}{r_i^2} + \frac{3}{\lambda_i r_i} + \frac{1}{\lambda_i^2} \right) \hat{r}_i \hat{r}_i - \left( \frac{1}{r_i^2} + \frac{1}{\lambda_i r_i} \right) I \right] G_i$$

$$\vec{t}_i \cdot (\nabla\nabla G_i) \cdot \vec{t}_i = \left[ \left( \frac{3}{r_i^2} + \frac{3}{\lambda_i r_i} + \frac{1}{\lambda_i^2} \right) (\vec{t}_i \cdot \hat{r}_i)^2 - \left( \frac{1}{r_i^2} + \frac{1}{\lambda_i r_i} \right) \right] G_i$$

so that

$$p_i : \nabla\nabla G_i = \left[ (1 - 3(\vec{t}_i \cdot \hat{r}_i)^2) \left( \frac{1}{r_i^2} + \frac{1}{\lambda_i r_i} \right) + \frac{1 - (\vec{t}_i \cdot \hat{r}_i)^2}{\lambda_i^2} \right] G_i$$

and

$$C^{(2)}_{\text{monopole}}(\vec{x}) = \sum_i \int_0^{L_i} \frac{a_i^2 q_i(s)}{4} \left[ (1 - 3(\vec{t}_i \cdot \hat{r}_i)^2) \left( \frac{1}{r_i^2} + \frac{1}{\lambda_i r_i} \right) + \frac{1 - (\vec{t}_i \cdot \hat{r}_i)^2}{\lambda_i^2} \right] G_i ds$$

and the total monopole contribution to order $O(a^2)$ is

$$C_{\text{monopole}}(\vec{x}) = C^{(0)}_{\text{monopole}}(\vec{x}) + C^{(2)}_{\text{monopole}}(\vec{x})$$

$$C_{\text{monopole}}(\vec{x}) = \sum_i \int_0^{L_i} \left( 1 + \frac{a_i^2}{4} \left[ (1 - 3(\vec{t}_i \cdot \hat{r}_i)^2) \left( \frac{1}{r_i^2} + \frac{1}{\lambda_i r_i} \right) + \frac{1 - (\vec{t}_i \cdot \hat{r}_i)^2}{\lambda_i^2} \right] \right) q_i(s) G(\vec{x}, \vec{X}_i(s)) ds$$

##### 6.3.3 The dipole contribution

The second term in the integral (the concentration dipole contribution), from the full boundary representation, is

$$C_{\text{dipole}}(\vec{x}) = -D \int_{\partial\Omega_t} C(\vec{x}')(\nabla G(\vec{x},\vec{x}') \cdot \vec{n}) dA' = -D \sum_i \int_0^{L_i} \int_0^{2\pi} C_{w,i}(s)(\nabla G(\vec{x},\vec{x}') \cdot \vec{n}_i) a_i d\theta ds$$

with $dA' = a_i d\theta ds$.

Now we expand the Green's function gradient about the centerline point $\vec{X}_i(s)$ and dot with $\vec{n}$:

$$\nabla G(\vec{x}, \vec{X}_i - a_i\vec{n}_i) \cdot \vec{n}_i = \nabla G(\vec{x}, \vec{X}_i) \cdot \vec{n}_i - a_i\vec{n}_i \cdot \left(\nabla\nabla G(\vec{x}, \vec{X}_i)\right) \cdot \vec{n}_i + O(a_i^2)$$

Substituting this into the integral for the concentration contribution of a single vessel:

$$C_{\text{dipole},i}(\vec{x}) = -a_i D \int_0^{L_i} C_{w,i}(s) \int_0^{2\pi} \left(\nabla G(\vec{x}, \vec{X}_i) \cdot \vec{n}_i - a_i\vec{n}_i \cdot \left(\nabla\nabla G(\vec{x}, \vec{X}_i)\right) \cdot \vec{n}_i + O(a_i^2)\right) d\theta ds$$

The leading-order term is

$$C^{(1)}_{\text{dipole},i}(\vec{x}) = -a_i D \int_0^{L_i} C_{w,i}(s) \nabla G(\vec{x}, \vec{X}_i) \cdot \left[\int_0^{2\pi} \vec{n}_i d\theta\right] ds$$

For a circular cross-section, $\int_0^{2\pi} \vec{n}_i d\theta = 0$; therefore, the leading order term vanishes $C^{(1)}_{\text{dipole},i}(\vec{x}) = 0$ because a uniform wall concentration around a circular vessel has no net dipole moment. Note that there is no $C^{(0)}_{\text{dipole}}$ term.

The next term is

$$C^{(2)}_{\text{dipole},i}(\vec{x}) = a_i^2 D \int_0^{L_i} C_{w,i}(s) \int_0^{2\pi} \vec{n}_i \cdot \left(\nabla\nabla G(\vec{x}, \vec{X}_i)\right) \cdot \vec{n}_i d\theta ds$$

Since $\vec{n}_i \cdot (\nabla\nabla G) \cdot \vec{n}_i = (\vec{n}_i \otimes \vec{n}_i) : \nabla\nabla G$,

$$C^{(2)}_{\text{dipole},i}(\vec{x}) = a_i^2 D \int_0^{L_i} C_{w,i}(s) \left[\int_0^{2\pi} \vec{n}_i \otimes \vec{n}_i d\theta\right] : \nabla\nabla G(\vec{x}, \vec{X}_i(s)) ds$$

For a circular cross-section,

$$\int_0^{2\pi} \vec{n}_i \otimes \vec{n}_i d\theta = \pi p_i$$

where $p_i = I - \vec{t}_i \otimes \vec{t}_i$.

Since $p_i : \nabla\nabla G = \nabla^2 G - \vec{t}_i \cdot (\nabla\nabla G) \cdot \vec{t}_i$, the first nonzero finite-radius correction (due to the dipole integral term) is

$$C_{\text{dipole}}(\vec{x}) = C^{(2)}_{\text{dipole}}(\vec{x}) = \pi D \sum_i a_i^2 \int_0^{L_i} C_{w,i}(s) \left(\nabla^2 G(\vec{x}, \vec{X}_i(s)) - \vec{t}_i(s) \cdot (\nabla\nabla G(\vec{x}, \vec{X}_i(s))) \cdot \vec{t}_i(s)\right) ds$$

Using $G_i$ and $\hat{r}_i$ from Section 6.3.2 for locally frozen (constant) $\lambda_i$, we recall that

$$p_i : \nabla\nabla G_i = \left[(1 - 3(\vec{t}_i \cdot \hat{r}_i)^2)\left(\frac{1}{r_i^2} + \frac{1}{\lambda_i r_i}\right) + \frac{1 - (\vec{t}_i \cdot \hat{r}_i)^2}{\lambda_i^2}\right] G_i$$

Therefore, the constant-$\lambda$ dipole correction is

$$C_{\text{dipole}}(\vec{x}) = \pi D \sum_i a_i^2 \int_0^{L_i} C_{w,i}(s) \left[(1 - 3(\vec{t}_i \cdot \hat{r}_i)^2)\left(\frac{1}{r_i^2} + \frac{1}{\lambda_i r_i}\right) + \frac{1 - (\vec{t}_i \cdot \hat{r}_i)^2}{\lambda_i^2}\right] G(\vec{x}, \vec{X}_i(s)) ds$$

##### 6.3.4 The final-form concentration field equation $C(\vec{x})$

Collecting all nonzero terms through order $O(a^2)$, the concentration field is...

$$C(\vec{x}) = C^{(0)}_{\text{monopole}}(\vec{x}) + C^{(2)}_{\text{monopole}}(\vec{x}) + C^{(2)}_{\text{dipole}}(\vec{x})$$

$$C(\vec{x}) = \sum_i \int_0^{L_i} \left[ q_i(s)G(\vec{x},\vec{X}_i(s)) + a_i^2 \left( \frac{q_i(s)}{4} + \pi D C_{w,i}(s) \right) \left( \nabla^2 G(\vec{x},\vec{X}_i(s)) - \vec{t}_i \cdot (\nabla\nabla G(\vec{x},\vec{X}_i(s))) \cdot \vec{t}_i \right) \right] ds$$

and for locally frozen $\lambda_i$ values,

$$C(\vec{x}) = \sum_i \int_0^{L_i} \left[ q_i(s) + a_i^2 \left( \frac{q_i(s)}{4} + \pi D C_{w,i}(s) \right) \left[ (1 - 3(\vec{t}_i \cdot \hat{r}_i)^2) \left( \frac{1}{r_i^2} + \frac{1}{\lambda_i r_i} \right) + \frac{1 - (\vec{t}_i \cdot \hat{r}_i)^2}{\lambda_i^2} \right] \right] G(\vec{x},\vec{X}_i(s)) ds$$

To leading order,

$$C(\vec{x}) = \sum_i \int_0^{L_i} q_i(s) G(\vec{x},\vec{X}_i(s)) ds$$

For vessel-vessel interactions, we expand only the finite-radius source vessel contribution and do not additionally expand the target vessel wall average. This is because the source-side finite-radius corrections arise from the singular near-field structure of the emitting vessel, whereas the target-side correction acts only on the smooth regular external field $C^{\text{ext}}$, which we assume to vary slowly over the vessel radius. Consequently, the target-side term $\frac{a_i^2}{4} p_i : \nabla\nabla C_i^{\text{ext}}$ is neglected.

See Appendix G for some simple scaling analyses of $C^{(2)}_{\text{monopole}}$ and $C^{(2)}_{\text{dipole}}$ relative to $C^{(0)}_{\text{monopole}}$.

#### 6.4 Solving for the external concentration field $C_i^{\text{ext}}$

The closed-form Green's function formulation also allows us to produce an expression for $C_i^{\text{ext}}$ in terms of the Green's function solutions (to $O(a^2)$ in source radius) from vessel-vessel interactions

$$C_i^{\text{ext}}(s) = \sum_j \int_0^{L_j} \left[ q_j(s')G_{ij}(s,s') + a_j^2 \left( \frac{q_j(s')}{4} + \pi D C_{w,j}(s') \right) p_j : \nabla\nabla G_{ij}^{\text{ext}}(s,s') \right] ds'$$

Where $p_j = I - \vec{t}_j \otimes \vec{t}_j$ is the projection tensor onto the plane normal to the source vessel axis. To leading order only,

$$C_i^{\text{ext}}(s) = \sum_j \int_0^{L_j} q_j(s') G_{ij}(s,s') ds'$$

Where $G_{ij}$ denotes the local Green's field evaluated at target vessel $i$ from source vessel $j$.

$$G_{ij}(s,s') = \frac{1}{4\pi D} \frac{e^{\frac{-||\vec{X}_i(s) - \vec{X}_j(s')||}{\lambda_j}}}{||\vec{X}_i(s) - \vec{X}_j(s')||}$$

Note that when evaluating the $C_i^{\text{ext}}$ sum computationally, we subtract out the self-contribution of the individual node for which we are performing the calculation.

##### 6.4.1 Iteration scheme

Please see Appendix B for information regarding iteration to convergence.

### 7 Tissue viability prediction from oxygen concentration

We defined a single critical cutoff oxygen concentration $C^*$, at which prolonged anoxic stress causes cell necrosis or permanent loss of function.

$$\text{Viability} = \mathcal{H}(C - C^*)$$

We assume that, generally, a value of about $0.0014 \frac{\text{mol}}{\text{m}^3}$ (equivalent to 1 mmHg partial pressure of oxygen) is a reasonable choice for $C^*$[35, 36, 43, 46, 47]. This approach is supported by experimental studies [43, 47, 50] and has precedent in numerical modeling studies [2, 4, 36].

### 8 Validation of simulation results via comparison to FEM

#### 8.1 Finite element simulation setup

To validate the accuracy of the methods developed in this work, we performed 25 simulations in both our Green's function solver and using the finite-element method (FEM) performed in COMSOL Multiphysics 6.3. Flow and concentration were computed via the Laminar Flow module coupled to the Transport of Diluted Species module. The tissue domain was represented as a uniform distribution of identical cells with a Michaelis-Menten consumption term.

FEM simulations were performed under identical conditions to the Green's-based method:

- Prescribed inlet flow ($Q$) and outlet pressure(s) ($P_{out}$)
- Prescribed inlet oxygen concentration
- Outlet boundary is an "open" exit boundary
- No-slip walls
- Infinite medium: tissue outer boundaries are "open"
- Solutions were computed to steady-state
- Tissue was represented as a cubic domain of volume 0.037, 0.125, or 1 $\text{cm}^3$
- Vessels were represented by a 1-inlet, 1(or more)-outlet "angiosome" structure connected by dozens or hundreds of smaller vessels.

In the main text, we identified two dimensionless parameters that govern the viability of a certain voxel of tissue: oxygen supply versus metabolic demand ($\Phi$), and average vessel to vessel distance versus characteristic reaction diffusion length ($\Pi$).

We selected 25 case studies to analyze via FEM such that they span a reasonable set of ($\Pi$,$\Phi$) pairs. We spanned $\Pi = 0.5$ (tight vessel spacing) to 3.4 (under-vascularized), and $\Phi = 0.2$ (low supply) to 93.5 (very abundant supply) ensuring that diffusivity, kinetics, and flow rate remained reasonable for what one might see either in native biology or in an engineered tissue construct (which are typically at lower cell densities than native tissues, use immature cell types, or use highly porous hydrogel materials).

A representative vascular structure of these case studies (Case 11) is shown in Figure S8.

#### 8.2 Mesh creation

Meshes for the FEM simulations were produced via an implicit-web meshing algorithm (with post-processing). Three example surface structures produced by this method (with increasing vascular density) are shown in Figure S9. The mesh production followed these steps:

1. Define the computational domain: A bounding box is computed from all vascular segment endpoints and padded by a multiple of the maximum vessel radius. The domain is discretized into a 3D Cartesian grid with a small voxel size.
2. Initialize an implicit field: A scalar field $\phi(x)$ is initialized on the grid with a large finite value. Each vessel segment is represented by endpoints and radius ($\vec{p}_0, \vec{p}_1, a$).

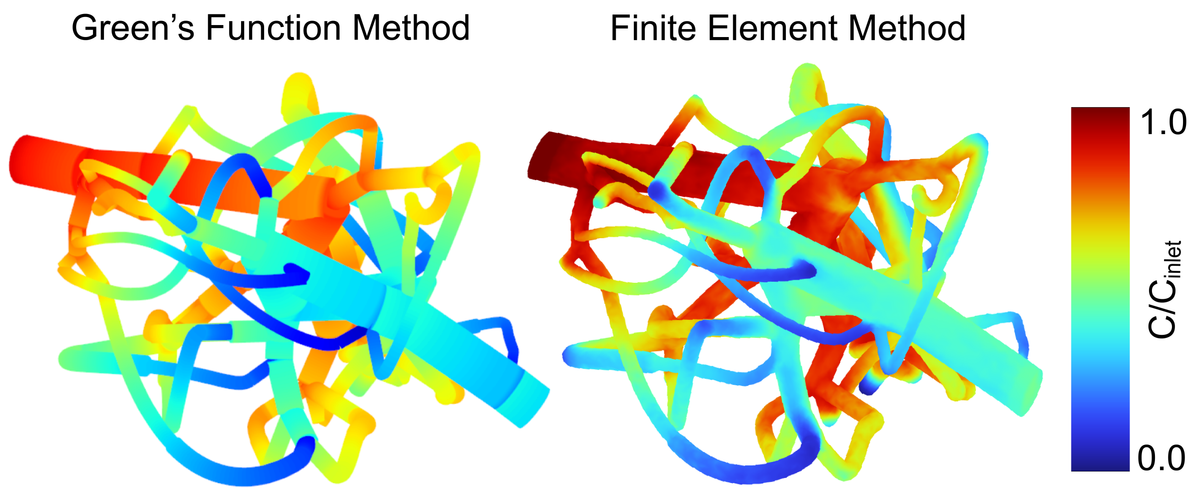

Figure S8: Case 11 with Green's function (left) and FEM (right) solutions for channel wall concentrations. This case represents a 0.125 $cm^3$ cubic domain of tissue, consisting of HEK-293T cells seeded at 60 million cells/mL in 10% (w/v) GelMA, being perfused by 100 $\mu$L/min of cell media.

3. Accumulate normalized distance to vessel network: For each grid point $\vec{x}$, the algorithm computes the minimum normalized distance to all centerline segments:

$$\phi(\vec{x}) = \mathrm{min_i} \frac{d(\vec{x}, S_i)}{a_i}$$

where $d(\vec{x}, S_i)$ is the shortest distance from the voxel point $\vec{x}$ to segment $S_i$, and $a_i$ is that segment's radius. Computation is restricted to local subvolumes around each segment for efficiency.

4. Extract the vessel surface: The vascular surface is defined as the iso-surface $\phi(\vec{x}) = 1$. This corresponds to points approximately one local vessel radius away from the centerline. Marching cubes is used to convert this iso-surface into a triangular mesh.

5. Post-processing in PyMeshLab: MeshLab filters are used to remove unreferenced vertices and duplicate faces, repair non-manifold edges, close holes, and apply isotropic remeshing/ball-pivoting reconstruction (which smooths the cubes into a uniform curved surface).

6. Enforce topology and clean geometry: The largest connected structure is retained, and PyMeshFix is used to enforce a closed two-manifold surface. Duplicate/degenerate faces, unreferenced vertices, and inconsistent normals are then repaired.

7. Validation and export: The final surface mesh is exported and checked for watertightness, manifold status, and Euler number. If needed, additional repair operations are performed.

8. Importing into COMSOL and surface processing: The surface mesh is imported into COMSOL Multiphysics, then simplified via conversion to an implicit body and refined by splitting the longest edge of each triangular face.

9. Creating volume mesh: Once an implicit domain is identified, a volumetric mesh is created via COMSOL's adaptive tetrahedral meshing protocol. Corner refinement is performed to improve meshing at bends and twists. Any intersecting faces or edges are collapsed and locally remeshed. To complete the mesh, several quad-mesh boundary layers are added to accurately capture parabolic flow profiles. The tissue domain is meshed via a similar protocol using a tetrahedral mesh.

10. Iterative refinement: The mesh is further refined until the solution converges to a relatively constant result (identical flow and concentration distributions).

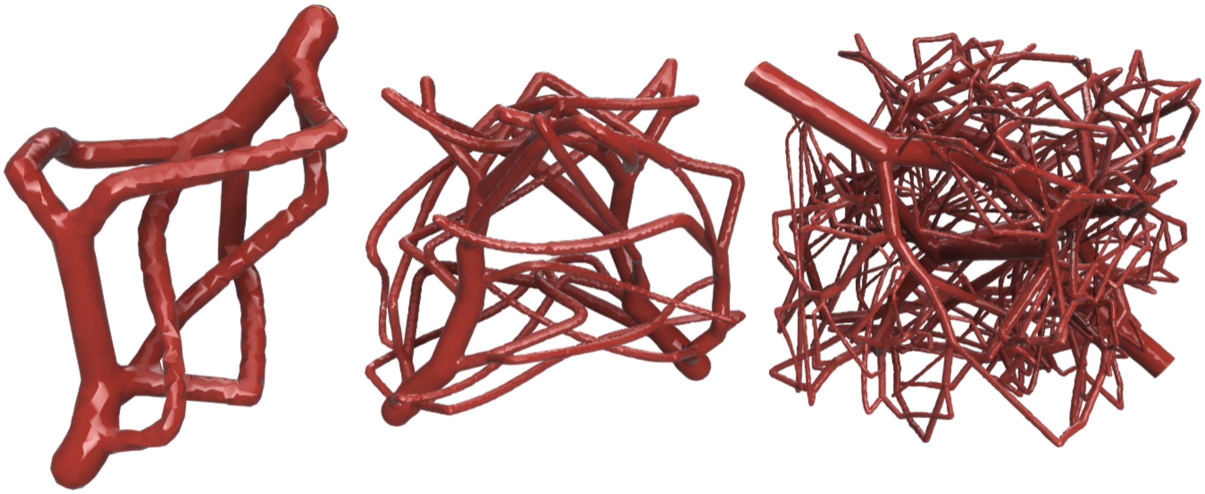

Figure S9: Three example meshes, increasing in vascular density from left to right.

##### 8.3 1-to-1 comparison results

For each case, 10,000 oxygen point sampling locations were chosen on a uniform grid. Any points reading `NaN` from the FEM or CASCADE solution were removed, resulting in an average number of evaluations per case of 9326.

Using all cases 1-25 pooled together, in Table S2 we report error metrics for all evaluations (left) and "low-concentration" evaluations, defined as any FEM-predicted concentration value below 0.005 mol/m$^3$. For reference, the Michaelis-Menten half-saturation constant $K_M$ is roughly this value for most mature cell types (with respect to oxygen), so "low-concentration" samples can be interpreted as point samples from regions likely to be substrate-limited and potentially associated with locally anaerobic metabolism.

| Error metric | All evaluations | Low-C measurements |
|---|---|---|
| Total number of point evaluations ($N_{evaluations}$) | 233140 | 144513 |
| Concordance correlation coefficient (CCC) | 0.9551 | 0.7374 |
| Root mean square error (RMSE) | 0.0087 | 0.0010 |
| Mean absolute error (MAE) | 0.0039 | 0.00057 |
| Coefficient of determination ($R^2 = 1 - SSE/SST$) | 0.9193 | 0.5347 |
| Squared Pearson correlation ($\rho^2$) | 0.9296 | 0.5918 |
| Mean signed error / bias | -0.0021 | 0.00033 |
| Live/dead prediction accuracy | 96.8% | N/a |
| False-alive prediction rate | 2.0% | N/a |
| False-dead prediction rate | 4.5% | N/a |

Table S2: Error metrics reported for all oxygen point evaluations across 25 case studies comparing the Green's function method to FEM. RMSE, MAE, and bias are reported in $mol/m^3$.

We report both the predictive coefficient of determination $R^2$ and the squared Pearson correlation $\rho^2$. We also report the concordance correlation coefficient (CCC), which measures agreement relative to the 1:1 parity line rather than only linear correlation:

$$\rho_{CCC} = \frac{2\rho\sigma_x\sigma_y}{\sigma_x^2 + \sigma_y^2 + (\mu_x - \mu_y)^2}$$

where $\rho$ is the Pearson correlation, $\mu_x$ and $\mu_y$ are the means, and $\sigma_x$ and $\sigma_y$ are the standard deviations of the two simulation outputs. CCC is important here because a high ordinary correlation or $\rho^2$ can still occur even when one simulation is systematically biased or scaled incorrectly; CCC penalizes both lack of correlation and deviation from the 1:1 parity line.

Overall, the pooled results show strong agreement across all evaluations, with CCC≈0.96, $R^2$≈0.92, $\rho^2$≈0.93, RMSE≈0.009, MAE≈0.004, and a small negative bias of -0.0021. In the low-concentration subset, absolute errors are much smaller, with RMSE≈0.001 and MAE≈0.0006, but the agreement metrics drop to CCC≈0.74, $R^2$≈0.53, and $\rho^2$ ≈0.59. This is expected in part because low-concentration samples span a narrower dynamic range and occur near more nonlinear reaction behavior (increased reaction nonlinearity near $K_M$).

Finally, using a live/dead cutoff of 0.0014 $\mathrm{mol/m^3}$ (equivalent to 1 mmHg partial pressure of oxygen), the two methods agree on classification for 96.8% of all points, with low false-alive and false-dead rates of 2.0% and 4.5%, respectively.

Figure S10 displays all 25 cases on both 1:1 parity scatterplots and through an array of confusion matrices showing the oxygen field agreement and live/dead classification agreement. Especially on the log/log parity plot, the strong agreement between the FEM evaluations and Green's function evaluations can be clearly seen. Thus, despite some degradation in pointwise concordance in the most challenging low-C, nonlinear regimes, the Green's function method developed in this work reproduces the FEM solution well overall, both in continuous concentration values and in the biologically relevant live/dead classification.

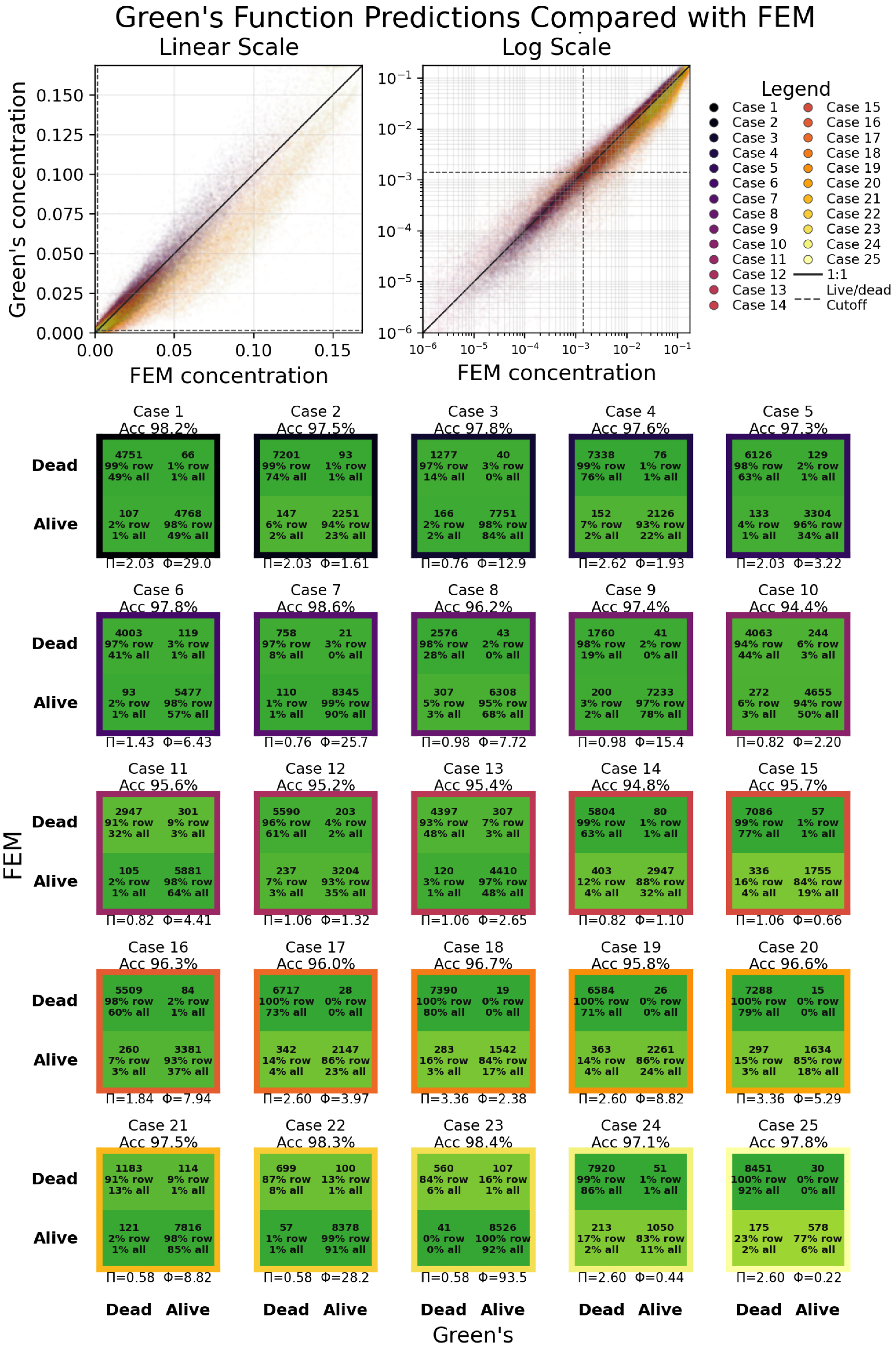

Figure S10: 1-to-1 parity plots (top) showing oxygen point samples for the same 25 steady-state simulation case studies performed with FEM and our Green's function solver. Two scales are shown: a linear scale (left) and log-base-10 scale (right). Scatter-points signify oxygen measurements at one (x,y,z) location in the perfused tissue domain. Each case study is paired with a confusion matrix (bottom) showing live/dead classification accuracy.

### 9 Big $O$ time scaling and important solver algorithms

#### 9.1 Big $O$ time scaling theory

In the main text of this paper, we reported time scaling for existing Laplace-GFMs (LGFMs), naive YGFM, windowed YGFM, and FFT YGFM protocols. Below are the derivations for each method. Henceforth $N$ is the vessel quadrature (number of source points), $M$ refers to the number of tissue points evaluated (in the LGFM case they are also consumptive sinks), $n$ is the tissue-grid resolution along one dimension (note $M = n^3$), $B$ and $G$ are the number of FFT $\lambda$ bins and cells respectively, $K_v$ and $K_t$ are the average number of $\lambda$-window-retained vessel and tissue sources per vessel/tissue target, and $W$ is the so-called "Window Factor" or number of $\lambda$s within which we retain source contributions. Iteration counters are left out for simplicity.

**LGFM**: Each vessel source $N$ and each tissue sink $M$ has a pairwise interaction. Therefore,

$$T_{LGFM} = O(N^2 + 2NM + M^2) = O((N + M)^2)$$

**Naive YGFM**: Each of the $N$ vessel targets receives contributions from every one of the $N$ vessel sources,

$$T_v \sim O(N^2)$$

and each of the $M$ tissue targets receives contributions from all $N$ vessel sources,

$$T_t \sim O(NM)$$

therefore the total scaling is

$$T_{naive} = O(N^2 + NM)$$

**Windowed YGFM**: A spatial index first assigns vessel sources to cells. A target then examines only sources inside its $W\lambda$ neighborhood. The spatial-index construction is approximately $O(N)$ for a uniform list (sort- or tree-based implementations could potentially reach $O(N\log N)$, but this part is generally cheap in comparison to other terms).

Each of the $N$ vessel targets examines an average of $K_v$ retained sources

$$T_v = O(K_v N)$$

and each $M$ tissue target examines an average of $K_t$ retained sources

$$T_t = O(K_t M)$$

Therefore

$$T_{windowed} = O(N + K_v N + K_t M)$$

or, dropping the relatively small index construction,

$$T_{windowed} = O(K_v N + K_t M)$$

**FFT YGFM**: The FFT replaces the windowed vessel-vessel background and tissue calculations, with an optional small correction in a reduced number of $\lambda$s (typically 0.5-1$\lambda$) in a radius around each quadrature element.

Each vessel quadrature point deposits into an interpolation stencil $O(N)$.

Each $\lambda$ bin has a different Yukawa multiplier, and accounting for all $B$ bins requires approximately $O(BG\log G)$. This is a typical cost for a fast Fourier transform.

The combined field is interpolated once at every vessel target $O(BN)$. Any vessel corrections or tissue corrections scale like $O(k_{corr,v}N)$ and $O(k_{corr,t}M)$, respectively. The grid field tissue sampling also costs $O(M)$.

So in total,

$$T_{FFT} = O(BN + BG\log G + k_{corr,v}N + (1 + k_{corr,t})M)$$

where $k_{corr} \ll K_v$ and $K_t$.

#### 9.2 Descriptions of important acceleration methods

To achieve the computation speed reported, we designed and implemented many key sub-algorithms and numerical solver acceleration tricks. The following list reports some sub-algorithms we employed.

1. **$\lambda$ screening and spatial indexing:** When computing tissue oxygen $C(\vec{x})$ and external concentration $C^{\text{ext}}$, we need not consider the contributions of every single vessel, rather only the contributions that are non-negligible. Instead, we screen all of our neighbor queries by a finite radius of $N\lambda$ where $\lambda$ is the contributing vessel's reaction-diffusion length scale.

   At a distance of $N\lambda$ from a contributing channel, for the first 6 $N$ values, the exponential screening factor decreases to (~36.8%, ~13.5%, ~5.0%, ~1.8%, ~0.7%, ~0.3%) respectively. We find that results settle around $5\lambda$, but use $6\lambda$ for all analyses performed in the main text. See Appendix C for an example.

   Spatial grid-cell lists and KD-tree indices are used to perform these screened neighbor searches efficiently, so that each tissue point (for $C(\vec{x})$) or vessel target (for $C^{\text{ext}}$) queries only nearby spatial regions before exact distance checks and Green's function evaluations are applied.

2. **GPU acceleration and custom kernels:** Intensive, parallelizable calculations are offloaded to the GPU wherever practical (Green's function evaluations, FFT grid deposition and sampling, tissue point sampling, Graetz updates, etc.). All GPU-accelerated calculations were distributed across 20 custom CUDA C kernels implemented through CuPy RawKernel (with the exception of our FFT solver which runs on cuFFT). This allows the most expensive loops to run with thousands of concurrent threads while the CPU handles setup, spatial indexing, and batching.

3. **Fast Fourier transform (FFT) solver:** For tissue scale computations ($\lesssim$ 10 cm$^3$), we found that $C^{\text{ext}}$ calculations could be accelerated >10× if deposited on a regular Cartesian grid and solved via FFT. Appendix D explains this and shows an example.

4. **External field $C^{\text{ext}}$ iteration tricks:** We compute the external field using a combination of multiple numerical tricks...

   (a) **Frozen sources:** Sources whose values are stabilized for many iterations can be frozen, having their contribution cached and reused.

   (b) **Anderson acceleration:** fixed-point iterations are accelerated by solving a least-squares problem over a window of past residuals to compute an extrapolated update that better approximates the fixed point. This decreases the number of required iterations for convergence, especially for large trees.

   (c) **Adaptive Aitken relaxation:** When Anderson is not stable, the solver falls back to adaptive Aitken $\omega$ updates. Aitken's method dynamically updates the relaxation parameter $\omega$ based on the observed convergence behavior of successive iterates, increasing or decreasing damping to improve stability and convergence.

   (d) **Trust/rejection logic and state recovery:** A series of rules are implemented so that residual-worsening steps can be rejected and retried with smaller relaxation. This avoids wasting iterations on unstable nonlinear updates. The solver tracks the best-seen residual state and can revert or stop the solve when the nonlinear map stalls or walks uphill.

   (e) **Decoupled Green's initialization:** We perform a cheap predictor solve before the fully coupled $C^{\text{ext}}$ iteration begins. The solver starts with all $C^{\text{ext}}$ values set to zero, propagates intravascular concentrations downstream once, then computes one Green's-function $C^{\text{ext}}$ field from the resulting vessel source strengths. This is then used as the initial guess for the real coupled iteration, instead of starting the nonlinear solve from an all-zero external field. Although this does not change the final converged solution, the decoupling of this initial preconditioning step greatly improves the stability of the solver.

5. **Lookup tables:** Any computations which are very frequent and only have one or two input parameters are accelerated with persisted lookup tables. Examples include modal-basis tables over Biot number ranges (for Graetz intraluminal diffusion) and Bessel $K_1/K_0$ ratio evaluations (for mass transfer constant $\kappa$ computations), among others.

### Appendix A Definitions and values

| Symbol | Name | Explanation | Units | Citation |
|---|---|---|---|---|
| $a_i$ | Vessel radius | Radius of vessel or channel $i$; also used as $a$ when the vessel index is omitted. | m | |
| $\alpha$ | Oxygen solubility | Solubility relating dissolved oxygen concentration to partial pressure, $C = \alpha P$. Value at 37°C is $1.408 \times 10^{-3} \frac{\text{mol}}{\text{m}^3\text{mmHg}}$ | $\text{mol}\,\text{m}^{-3}\,\text{mmHg}^{-1}$ | [24] |
| $\alpha_k$ | Fourier-mode radial decay rate | Mode-dependent radial decay rate when axial diffusion is retained, $\alpha_k = \sqrt{k^2 + \lambda^{-2}}$. | $\text{m}^{-1}$ | |
| $A_{n,i}$ | Graetz modal coefficient | Expansion coefficient multiplying eigenfunction $\phi_{n,i}$ in the intraluminal Graetz eigenfunction solution. | | |
| $B$ | Incidence matrix | Edge–node incidence matrix for the hydraulic network; maps nodal pressures to edge pressure drops. | | |
| $\text{B}_i$ | Hemoglobin buffer factor | Frozen buffer multiplier in the intraluminal convection–diffusion equation; equals 1 for water/plasma/media and $\beta_i/\alpha$ for blood. | | |
| $\beta(P)$ | Hemoglobin buffering derivative | Derivative factor $\alpha + C_{\text{Hb}}^{\text{max}}\, dS/dP$ governing the effect of hemoglobin buffering on dissolved oxygen changes. | $\text{mol}\,\text{m}^{-3}\,\text{mmHg}^{-1}$ | |
| $\text{Bi}_i$ | Lumen–tissue Biot number | Dimensionless ratio measuring tissue-side wall extraction relative to intraluminal radial diffusion, $\text{Bi}_i = a_i h_i / D_v = \kappa_i / (2\pi D_v)$. | | |
| $c_i(r,s)$ | Local radial disturbance | Radially varying disturbance field around vessel $i$ after subtracting the smooth external/background field. | $\text{mol}\,\text{m}^{-3}$ | |
| $\hat{c}_k(r)$ | Fourier-mode disturbance | Radial amplitude of axial Fourier mode $k$ for the disturbance field when axial diffusion is retained. | $\text{mol}\,\text{m}^{-3}$ | |
| $C$ | Concentration | Generic oxygen concentration; context determines whether it is nodal, intravessel, or tissue concentration. | $\text{mol}\,\text{m}^{-3}$ | |
| $C^\dagger$ | Unrelaxed concentration solution | Intermediate concentration solution obtained before Picard under-relaxation. | $\text{mol}\,\text{m}^{-3}$ | |
| $C(\vec{x})$ | Tissue oxygen concentration | Pointwise oxygen concentration in the tissue domain at location $\vec{x}$. | $\text{mol}\,\text{m}^{-3}$ | |
| $C_i(\vec{x})$ | Near-vessel concentration | Local tissue concentration near vessel $i$, decomposed as $C_i^{\text{reg}}(\vec{x}) + c_i(r,s)$. | $\text{mol}\,\text{m}^{-3}$ | |
| $C^*$ | Critical oxygen concentration threshold | Oxygen concentration below which prolonged anoxic stress leads to cell death or major loss of function. Taken as the dissolved concentration equivalent to about 1 mmHg $O_2$ partial pressure. | 0.0014 $\text{mol}\,\text{m}^{-3}$ | [35, 36, 43, 46, 47] |
| $C_i^{\text{reg}}(\vec{x})$ | Regular background concentration | Smooth background concentration near vessel $i$ after removing the singular/local vessel contribution. | $\text{mol}\,\text{m}^{-3}$ | |
| $C_i^{\text{ext}}(s)$ | External concentration | Regular external concentration experienced by vessel $i$ at axial position $s$ due to surrounding vessels/tissue field. | $\text{mol}\,\text{m}^{-3}$ | |
| $C_i^{\text{eff}}(s)$ | Effective concentration for length scale | Closure concentration used to evaluate the reaction–diffusion length scale for vessel $i$ | $\text{mol}\,\text{m}^{-3}$ | |
| $C_{\text{in}}$ / $C_{\text{root}}$ | Inlet/root concentration | Prescribed oxygen concentration at the root/inlet node. Measured to be $0.22\ \frac{\text{mol}}{\text{m}^3}$ for cell media and taken to be $0.14\ \frac{\text{mol}}{\text{m}^3}$ (about 100 mmHg) for arterial blood. | $\text{mol}\,\text{m}^{-3}$ | [4, 25] |
| $C_b$ | Bulk intravessel concentration | Flow-weighted dissolved oxygen concentration in the vessel lumen. | $\text{mol}\,\text{m}^{-3}$ | |
| $C_c$ | Intraluminal concentration | Generic dissolved oxygen concentration inside the channel/lumen. | $\text{mol}\,\text{m}^{-3}$ | |

| Symbol | Name | Explanation | Units | Citation |
|---|---|---|---|---|
| $C_{\text{Hb}}$ | Hemoglobin-bound oxygen concentration | Oxygen concentration bound to hemoglobin. $C_{\text{Hb}} = C_{\text{Hb}}^{\max} S(P_{O_2})$. | mol m$^{-3}$ | |
| $C_{\text{Hb}}^{\max}$ | Advected hemoglobin oxygen capacity | Maximum oxygen concentration capacity associated with hemoglobin binding, used in the advected oxygen-content relation. $C_{\text{Hb}}^{\max} = H_D C_{\text{RBC}}^{\max}$ | mol m$^{-3}$ | |
| $C_{\text{RBC}}^{\max}$ | Red-cell oxygen capacity | Fully saturated hemoglobin-bound oxygen capacity per unit red-cell volume | mol m$^{-3}$ of red cells | |
| $C_{w,i}(s)$ | Vessel wall concentration | Oxygen concentration at the vessel/channel wall of vessel $i$ along axial coordinate $s$. | mol m$^{-3}$ | |
| $C_{\text{total}}$ | Total blood oxygen concentration | Sum of dissolved and hemoglobin-bound oxygen, $C_b + C_{\text{Hb}}$. Equivalent to $T(C_b)$ | mol m$^{-3}$ | |
| $C_{\text{monopole}}$ | Monopole concentration contribution | Contribution to $C(\vec{x})$ from flux/source monopole terms in the Green's-function representation. | mol m$^{-3}$ | |
| $C_{\text{monopole}}^{(0)}$ | Leading-order monopole term | Leading line-source contribution to the tissue concentration field. | mol m$^{-3}$ | |
| $C_{\text{monopole}}^{(1)}$ | First-order monopole correction | First finite-radius monopole correction; zero for circular cross-sections under the stated assumptions. | mol m$^{-3}$ | |
| $C_{\text{monopole}}^{(2)}$ | Second-order monopole correction | Second-order finite-radius monopole correction proportional to $a_i^2 p_i : \nabla\nabla G_i$. | mol m$^{-3}$ | |
| $C_{\text{dipole}}$ | Dipole concentration contribution | Contribution to $C(\vec{x})$ from the concentration dipole term in the boundary representation. | mol m$^{-3}$ | |
| $C_{\text{dipole}}^{(0)}$ | Zeroth-order dipole term | Not present/zero in the expansion described; included because the text explicitly notes there is no $C_{\text{dipole}}^{(0)}$ term. | mol m$^{-3}$ | |
| $C_{\text{dipole}}^{(1)}$ | First-order dipole correction | First dipole correction; vanishes for a circular cross-section with uniform wall concentration. | mol m$^{-3}$ | |
| $C_{\text{dipole}}^{(2)}$ | Second-order dipole correction | First nonzero finite-radius dipole correction in the boundary representation. | mol m$^{-3}$ | |
| CCC | Concordance correlation coefficient | Agreement metric used in validation against FEM; also denoted $\rho_{\text{CCC}}$. | | |
| $\gamma_e$ | Edge attenuation factor | Factor describing how strongly edge $e$ retains its upstream concentration after transport and exchange. | | |
| $d(\vec{x}, S_i)$ | Point-to-segment distance | Shortest distance from mesh grid point $\vec{x}$ to centerline segment $S_i$ in the meshing algorithm. | m | |
| $D$ | Tissue oxygen diffusivity | Effective diffusivity of oxygen in the tissue/hydrogel domain. Taken as $2.8 \times 10^{-9}\,\frac{\text{m}^2}{\text{s}}$ for cellular tissues fabricated with 10% (w/v) GelMA (calculated via Millington-Quirk model), and as $\sim 2 \times 10^{-9}\,\frac{\text{m}^2}{\text{s}}$ for cardiac tissues. | m$^2$ s$^{-1}$ | [3, 34] |
| $D_v$ | Intraluminal oxygen diffusivity | Diffusivity of oxygen inside the vessel/channel lumen. At 37°C, its value is $3.2 \times 10^{-9}\,\frac{\text{m}^2}{\text{s}}$ for water/cell media, $2.18 \times 10^{-9}\,\frac{\text{m}^2}{\text{s}}$ for plasma, and $1.62 \times 10^{-9}\,\frac{\text{m}^2}{\text{s}}$ for human blood at 42% hematocrit. | m$^2$ s$^{-1}$ | [12] |
| $\delta(\vec{x} - \vec{X}_i)$ | Dirac delta | Three-dimensional Dirac forcing/source distribution that localizes a vessel line source at the centerline. | m$^{-3}$ | |
| $e$ | Edge index | Directed vessel segment/edge in the hydraulic and concentration graphs. | | |
| $\epsilon_i$ | Radius-to-length-scale ratio | Dimensionless ratio $a_i/\lambda_i$ used in scaling arguments. | | |

| Symbol | Name | Explanation | Units | Citation |
|---|---|---|---|---|
| $f(s)$ | Axial wall mismatch | Difference driving the disturbance in the axial-diffusion appendix, $f(s) = C_w(s) - C^{\text{ext}}(s)$. | $\text{mol m}^{-3}$ | |
| $\hat{f}(k)$ | Fourier transform of wall mismatch | Axial Fourier transform of $f(s)$. | $\text{mol m}^{-2}$ | |
| $F_i$ | Nondimensional curvature factor | Dimensionless form of the geometric curvature factor used in finite-radius scaling. | | |
| $\Pi$ | Vessel spacing parameter | Dimensionless parameter identified in the main-text: average vessel-vessel distance relative to a global reaction–diffusion length. | | |
| $\Phi(\vec{x})$ | FFT background concentration field | Smooth grid-based background concentration field satisfying a screened Poisson equation in the FFT approximation. | $\text{mol m}^{-3}$ | |
| $\phi_i$ | Nondimensional radius | Vessel radius scaled by local reaction–diffusion length, $\phi_i = a_i/\lambda_i$. | | |
| $\phi_{n,i}(\tilde{r})$ | Graetz radial eigenfunction | Radial eigenfunction for mode $n$ in vessel $i$ in the intraluminal Graetz problem. | | |
| $\phi'_{n,i}$ | Eigenfunction radial derivative | Derivative of the Graetz radial eigenfunction with respect to nondimensional radius $\tilde{r}$. | | |
| $G$ | Green's function | Yukawa/screened Green's function used to compute concentration from line-source fluxes. | $\text{s m}^{-3}$ | |
| $G_i(r)$ | Vessel-$i$ Green's kernel | Green's kernel for contribution from vessel $i$ with local distance $r_i$ and length scale $\lambda_i$. | $\text{s m}^{-3}$ | |
| $G_{ij}(s, s')$ | Pairwise Green's function | Green's function produced by source vessel $j$ at axial location $s'$ and evaluated at target vessel $i$ at axial location $s$. | $\text{s m}^{-3}$ | |
| $G_{ij}^{\text{ext}}(s, s')$ | External-field pairwise Green's function | Pairwise Green's-function term used in finite-radius external-field corrections. | $\text{s m}^{-3}$ | |
| $\nabla\nabla G$ | Green's-function Hessian | Second spatial derivative tensor of the Green's function. | $\text{s m}^{-5}$ | |
| $h_i(s)$ | Wall mass transfer coefficient | Local wall coefficient related to $\kappa_i$ by $h_i = \kappa_i/(2\pi a_i)$. | $\text{m s}^{-1}$ | |
| $H_D$ | Discharge hematocrit | Upstream/inlet discharge hematocrit used in blood rheology and phase separation. At the inlet of vessel structures, this was set to 42% (within reported ranges). | | [6] |
| $H_{T,i}$ | Tube hematocrit | Tube/vessel hematocrit in vessel $i$. | | |
| $i$ | Vessel/node index | Index for a vessel, target vessel, or graph node depending on context. | | |
| $\mathcal{I}(i)$ | Incoming-edge set | Set of edges flowing into node $i$. | | |
| $I$ | Identity tensor/matrix | Identity tensor used in the projection tensor and Hessian contractions; also identity matrix in boundary-condition elimination. | | |
| $j$ | Source vessel/node index | Index often used for a source vessel in pairwise vessel–vessel interactions or a downstream node in an edge. | | |
| $J_n$ | Normal oxygen flux | Outward oxygen flux per unit area entering the tissue from the vessel wall. | $\text{mol m}^{-2}\,\text{s}^{-1}$ | |
| $k$ | Iteration index | Picard/outer-iteration counter, as in superscript $(k)$. | | |
| $k$ | Axial Fourier wavenumber | Fourier wavenumber in the axial-diffusion appendix. | $\text{m}^{-1}$ | |
| $K_e$ | Hydraulic conductance | Hydraulic conductance of vessel/edge $e$, $K_e = R_e^{-1}$. | $\text{m}^3\,\text{Pa}^{-1}\,\text{s}^{-1}$ | |

| Symbol | Name | Explanation | Units | Citation |
|---|---|---|---|---|
| $K_M$ | Michaelis–Menten constant | Half-saturation constant in the oxygen uptake reaction rate. Taken as $0.036\,\frac{\text{mol}}{\text{m}^3}$ for HEK-293T cells and $0.007\,\frac{\text{mol}}{\text{m}^3}$ for human myocardial tissue (mature cardiomyocytes). | $\text{mol}\,\text{m}^{-3}$ | [16, 21, 23] |
| $K_0(\cdot)$ | Modified Bessel function | Modified Bessel function of the second kind of order zero. | | |
| $K_1(\cdot)$ | Modified Bessel function | Modified Bessel function of the second kind of order one. | | |
| $\mathcal{K}_i$ | Geometric curvature factor | Factor $p_i : \nabla\nabla G_i / G_i$ used in finite-radius scaling arguments; written as $K_i$ in the derivation but renamed here to avoid conflict with hydraulic conductance. | $\text{m}^{-2}$ | |
| $\kappa_i(s)$ | Interfacial mass transfer coefficient | Effective per-length coefficient relating wall–external concentration difference to outward flux per unit length. | $\text{m}^2\,\text{s}^{-1}$ | |
| $\tilde{\kappa}_i$ | Buffered interfacial coefficient | Hemoglobin-buffered mass transfer coefficient, $\tilde{\kappa}_i = (\alpha/\beta_i)\kappa_i$. | $\text{m}^2\,\text{s}^{-1}$ | |
| $L$ | Cubic-domain side length | Side length of a cubic tissue volume in the complexity-scaling discussion. | m | |
| $L_i$ | Vessel length | Length of vessel $i$ over which axial coordinate $s$ is integrated. | m | |
| $L_{v-v}$ | Average vessel-to-vessel distance | Mean distance between vessels used in iteration-scaling observations. | m | |
| $\lambda$ | Reaction–diffusion length scale | Generic characteristic length scale for locally linearized Michaelis–Menten uptake. | m | |
| $\lambda_{v,i}(s)$ | Vessel apparent length scale | Reaction–diffusion scale of vessel $i$ used for vessel-wall flux calculations and vessel–vessel external-field interactions. | m | |
| $\lambda_{t,i}(r,s)$ | Tissue apparent length scale | Reaction–diffusion scale of vessel $i$ used to evaluate the tissue Green's-function contribution at a field point. | m | |
| $\lambda_{\text{avg}}$ | Average length scale | Average reaction–diffusion length scale over all vessels, used in iteration and grid-resolution heuristics. | m | |
| $m$ | Substep index | Axial substep index within a vessel/edge subdivision. | | |
| $\mu_i$ | Dynamic viscosity | Effective dynamic viscosity in vessel/edge $i$ for hydraulic resistance calculations. | Pa s | |
| $\mu_{n,i}$ | Graetz eigenvalue | Eigenvalue for radial mode $n$ in vessel $i$ in the Graetz eigenproblem. | | |
| $\mu_x, \mu_y$ | Validation means | Means of two simulation outputs used in the concordance correlation coefficient. | $\text{mol}\,\text{m}^{-3}$ | |
| $n$ | Mode/root index | Eigenmode index in Graetz expansions; also used as a node/root-node index in the graph-solve text. | | |
| $N\lambda$ | Interaction screening length | Cutoff distance equal to $N$ reaction–diffusion lengths for excluding weak Green's-function interactions. | m | |
| $n_m$ | Number of retained eigenmodes | Number of Graetz eigenmodes retained in the intraluminal expansion. | | |
| $n_r$ | Number of radial profile points | Number of discrete radial points used to resolve the intraluminal profile. | | |
| $\vec{n}_i(s,\theta)$ | Boundary normal vector | Tissue-to-vessel normal vector on the vessel wall; points into the vessel. | | |
| $\mathcal{O}(i)$ | Outgoing-edge set | Set of edges flowing out of node $i$. | | |
| $O(a^2)$ | Second-order finite-radius order | Order notation for terms of size proportional to $a^2$ in the finite-radius expansion. | | |

| Symbol | Name | Explanation | Units | Citation |
|---|---|---|---|---|
| $\omega$ | Under-relaxation factor | Picard under-relaxation parameter satisfying $0 < \omega \leq 1$. | | |
| $\Omega_t$ | Tissue domain | Tissue domain over which the Green's representation is written. | $m^3$ | |
| $\partial\Omega_t$ | Tissue boundary | Boundary of the tissue domain used in Green's second identity. | $m^2$ | |
| $p_i$ | Projection tensor | Projection tensor onto the plane normal to vessel $i$'s tangent vector, $p_i = I - \vec{t}_i \otimes \vec{t}_i$. | | |
| $\vec{p}_0, \vec{p}_1$ | Segment endpoints | Endpoints of a vessel segment (in 3D space) used in the implicit-web meshing description. | m | |
| $P_{\text{in}}$ | Inlet pressure | Prescribed or solved pressure at the inlet/root node. Also used in vascular design generation. For 1 $cm^3$ cube of myocardium, 6666.1 Pa; for full heart, 12665.6 Pa. | Pa | [5, 14] |
| $P_{\text{out}}$ / $P_{\text{term}}$ | Outlet/terminal vessel pressure | Prescribed outlet pressure used in FEM validation conditions. Also used in vascular design generation. Typically set to 4000 Pa. | Pa | [5, 14] |
| $\Delta P_e$ | Pressure drop | Pressure drop across edge/vessel segment $e$. | Pa | |
| $\Phi$ | Supply-demand parameter | Dimensionless parameter identified in the main text: oxygen supply versus metabolic demand. | | |
| $Q$ | Flow rate | Generic volumetric flow rate through a vessel/channel. | $m^3\,s^{-1}$ | |
| $Q_{\text{in}}$ | Total inlet flow rate | Prescribed or solved total flow entering the root/inlet node. Also used in vascular design generation. For HEK-293T constructs, $25\frac{\mu L}{min}$; for 1 $cm^3$ cube of myocardium, $900\frac{\mu L}{min}$; for LCA, $189.6\frac{mL}{min}$; for RCA, $85.0\frac{mL}{min}$. | $\mu L\ min^{-1}$ | [26, 51, 52] |
| $q_i(s)$ | Outward flux per unit length | Oxygen flux per unit vessel length from vessel $i$ into the tissue. | $mol\,m^{-1}\,s^{-1}$ | |
| $\hat{q}_{r,s}(k)$ | Fourier wall-flux mode | Fourier transform of the axial+radial wall flux per unit length. | $mol\,s^{-1}$ | |
| $r$ | Radial coordinate/separation | Radial coordinate from a vessel centerline or generic source–target separation distance, depending on context. | m | |
| $r_i$ | Distance to vessel $i$ | Distance from field point $\vec{x}$ to vessel centerline location $\vec{X}_i(s)$. | m | |
| $r_{ij}$ | Pairwise vessel distance | Distance between target vessel point $\vec{X}_i(s)$ and source vessel point $\vec{X}_j(s')$. | m | |
| $\hat{r}_i$ | Unit orientation vector | Unit vector from source vessel point $\vec{X}_i(s)$ to tissue evaluation point $\vec{x}$. | | |
| $\tilde{r}$ | Nondimensional radius | Radial coordinate normalized by vessel radius, $\tilde{r} = r/a_i$. | | |
| $R_e$ | Hydraulic resistance | Hagen–Poiseuille hydraulic resistance of vessel/edge $e$. | $Pa\,s\,m^{-3}$ | |
| $R^2$ | Coefficient of determination | Validation statistic comparing Green's-function and FEM outputs. | | |
| $\rho$ | Pearson correlation | Pearson correlation coefficient used in the concordance correlation coefficient expression. | | |
| $\rho(\vec{x})$ | Gridded source density | FFT/grid source density formed by depositing vessel line-source fluxes onto a Cartesian grid. | $mol\,m^{-3}\,s^{-1}$ | |
| $\rho_i$ | Nondimensional distance | Distance normalized by reaction–diffusion length, $\rho_i = r_i/\lambda_i$. | | |
| $s$ | Axial coordinate | Coordinate along a vessel centerline or axial direction. | m | |
| $\tilde{s}$ | Nondimensional axial coordinate | Axial coordinate scaled for the Graetz intraluminal diffusion problem. | | |
| $\vec{s}$ | Source/sink flow vector | Vector containing prescribed hydraulic source or sink flows at graph nodes. | $m^3\,s^{-1}$ | |
| $s_{\text{root}}$ | Root source flow | Hydraulic source term at the root node, set to $Q_{\text{in}}$ for prescribed inlet flow. | $m^3\,s^{-1}$ | |

| Symbol | Name | Explanation | Units | Citation |
|---|---|---|---|---|
| $S(P)$ | Hemoglobin saturation | Severinghaus oxyhemoglobin binding/dissociation. Function of oxygen partial pressure. | | [41] |
| $S_i$ | Centerline segment | Vessel centerline segment used in the implicit-mesh distance calculation. | m | |
| $S_{\text{up}}$ / $S_{\text{down}}$ | Upstream/downstream selector matrix | Matrix selecting the upstream or downstream-node concentration for each edge. | | |
| $S(\vec{x})$ | Yukawa source distribution | Line-source forcing distribution in the tissue reaction–diffusion equation. | mol m$^{-3}$ s$^{-1}$ | |
| $\sigma$ | Dummy axial integration variable | Integration variable used in the well-mixed intravessel solution. | m | |
| $\sigma_x, \sigma_y$ | Validation standard deviations | Standard deviations of two simulation outputs in the concordance correlation coefficient. | mol m$^{-3}$ | |
| $\vec{t}_i$ | Vessel tangent vector | Unit tangent vector of vessel $i$, oriented with the flow direction. | | |
| $T$ | Temperature | Temperature of incubation conditions; body temperature. 37°C. | °C | |
| $\theta$ | Azimuthal angle | Angular coordinate around a circular vessel cross-section. | radians | |
| $\Theta_i(\tilde{r}, \tilde{s})$ | Nondimensional lumen concentration | Intraluminal concentration nondimensionalized by the inlet bulk concentration to the vessel subdivision. | | |
| $\Theta_i^{\text{ext}}$ | Nondimensional external concentration | External concentration nondimensionalized by subdivision inlet bulk concentration. | | |
| $\tilde{\Theta}_i(\tilde{r}, \tilde{s})$ | Nondimensional excess concentration | Difference between nondimensional lumen and external concentrations, $\Theta_i - \Theta_i^{\text{ext}}$. | | |
| $\tilde{\Theta}_{i,0}(\tilde{r})$ | Incoming excess profile | Incoming nondimensional excess concentration profile at the start of a frozen axial interval. | | |
| $u_i(r)$ | Velocity profile | Dimensional intraluminal velocity profile in vessel $i$. | m s$^{-1}$ | |
| $U_i$ | Mean velocity | Mean velocity in a straight vessel segment, $U_i = Q_i/(\pi a_i^2)$. | m s$^{-1}$ | |
| up($e$) / down($e$) | Upstream/downstream node | Node at the upstream or downstream end of edge $e$ after orienting the edge with the computed flow. | | |
| $v_i(\tilde{r})$ | Normalized velocity profile | Velocity profile normalized by mean velocity, $v_i = u_i/U_i$; for Poiseuille flow, $v_i = 2(1 - \tilde{r}^2)$. | | |
| $V_{\max}$ | Maximum uptake rate | Maximum oxygen uptake/consumption rate in the Michaelis–Menten reaction term. $3.85 \times 10^{-17} \frac{\text{mol}}{\text{cell s}}$ for HEK-293T cells and $0.044 \frac{\text{mol}}{\text{m}^3\text{s}}$ for human myocardium. | mol m$^{-3}$ s$^{-1}$ or mol cell$^{-1}$ s$^{-1}$ | [8, 9, 18, 40, 48] |
| $w_i$ | Nodal flow weight | Flow-weight normalization for node $i$ in the graph concentration balance. | m$^3$ s$^{-1}$ | |
| $W$ | Nodal flow-weight matrix | Diagonal matrix with entries $w_i$. | m$^3$ s$^{-1}$ | |
| $\vec{x}$ | Field/evaluation point | Point in tissue or on a grid where oxygen concentration is evaluated. | m | |
| $\vec{x}'$ | Source/boundary point | Source coordinate or boundary point in the Green's representation. | m | |
| $\vec{X}_i(s)$ | Vessel centerline | Centerline location of vessel $i$ at axial coordinate $s$, oriented in the flow direction. | m | |
| $\chi_e$ | External-field contribution | Additive contribution in the edge concentration update; for constant external field, $\chi_e = (1 - \gamma_e)C_e^{\text{ext}}$. | mol m$^{-3}$ | |
| $\langle f, g \rangle$ | Velocity-weighted inner product | Inner product used to exploit orthogonality of Graetz eigenfunctions. | | |

| Symbol | Name | Explanation | Units | Citation |
|---|---|---|---|---|
| $\nabla$ | Gradient operator | Spatial gradient, usually with respect to source or field coordinates as specified in text. | $m^{-1}$ | |
| $\nabla^2$ | Laplacian operator | Spatial Laplacian in the reaction–diffusion or Green's-function equations. | $m^{-2}$ | |
| $\cdot$ | Dot product | Tensor single contraction, e.g., $\vec{t}_i \cdot \hat{r}_i$. | | |
| : | Double contraction | Tensor double contraction, e.g., $p_i : \nabla\nabla G$. | | |
| $\|\ \|$ | Euclidean norm | Vector norm used for distances and convergence criteria. | same as argument | |
| $\odot$ | Elementwise product | Hadamard/elementwise product used in the graph concentration system. | | |
| $\otimes$ | Outer product | Tensor outer product, e.g., $\vec{n}_i \otimes \vec{n}_i$ or $\vec{t}_i \otimes \vec{t}_i$. | | |

Table S3: List of all parameters used with explanations, values, units, and citations

### Appendix B The steady-state reaction–diffusion length scale $\lambda$ and the outer iteration scheme

#### B.1 Roles of the length scale

We distinguish two uses of the reaction–diffusion length scale:

- **Vessel apparent scale:** $\lambda_{v,i}(s)$, used in flux computations (through $\kappa_i$ and $q_i$), and vessel-vessel interactions through $C_i^{\text{ext}}$.
- **Tissue apparent scale:** $\lambda_{t,i}(r, s)$, used in the Green's function evaluation of $C(\vec{x})$ through additive contributions across nearby vessels.

These play distinct physical roles and are constructed differently. However, because they are consistently defined, both are written as $\lambda_i$ for simplicity.

#### B.2 Vessel side: Solving for intravessel concentration profiles

**(a) Initialization.** For our initial guess of $\lambda$, we first neglect vessel–vessel interactions prior to the first iteration ($k = 0$):

$$C_i^{\text{ext},(0)}(s) = 0,$$

and solve for $C_{w,i}^{(0)}(s)$. This lets us form initial guesses for each $\lambda$ parameter. The vessel scale is defined directly from the intravessel concentration:

$$\lambda_{v,i}^{(0)}(s) = \sqrt{\frac{D\left(K_M + C_{w,i}^{(0)}(s)\right)}{V_{\max}}}.$$

Using $\lambda_{v,i}^{(0)}$, we can compute the initial guesses for the interfacial mass transfer coefficient $\kappa_i^{(0)}$ and outward flux per unit length $q_i^{(0)}$ for each vessel $i$.

**(b) Compute far-field concentrations for each vessel.** To compute the external concentration $C_i^{\text{ext},(k)}$ felt by vessel $i$, we may sum contributions from all nearby channels $j$ by using $\lambda_{v,j}^{(k-1)}$ as the input to each $G_{ij}$ term.

Using the previous iteration's $(k-1)$ pairwise Green's solution, we estimate the external concentration each vessel $i$ experiences at the new iteration $(k)$.

$$C_i^{\text{ext},(k)}(s) = \sum_j \int_0^{L_j} G_{ij}^{(k-1)}(s, s')\, q_j^{(k-1)}(s')\, ds',$$

with the previous iteration's pairwise Green's function defined as

$$G_{ij}^{(k-1)}(s,s') \sim \frac{e^{-\frac{r_{ij}}{\lambda_j^{(k-1)}(s')}}}{4\pi r_{ij} D}, \quad r_{ij} = ||\vec{X}_i(s) - \vec{X}_j(s')||$$

Note that $G_{ij}(s,s')$ is the Green's function produced by vessel $j$ (source) at axial location $s'$ felt by vessel $i$ (target) at axial location $s$.

**(c) Update intravessel profiles.** We now wish to know how $C_{w,i}$ changes along each vessel $i$'s length at iteration $(k)$. Here we may use the previous iteration's $\tilde{\kappa}_i$ value to maintain a linear system. We assume $Q_i$ does not vary along $s$.

$$Q_i \frac{d(C_{b,i}^{(k)})}{ds} = -\tilde{\kappa}_i^{(k-1)}(s)\left(C_{w,i}^{(k)} - C_i^{\text{ext},(k)}\right).$$

**(d) Update characteristic length scales.**

**Interfacial (near-wall):** This scale remains with respect to the channel concentration, as this is the relevant concentration near the interface.

$$\lambda_{v,i}^{(k)}(s) = \sqrt{\frac{D\left(K_M + C_{w,i}^{(k)}(s)\right)}{V_{\max}}}.$$

We then update the mass transfer coefficient $\kappa_i(s)$ and outward flux per unit length $q_i(s)$ for each vessel given their new channel concentration profile $C_{w,i}(s)$ and external concentration $C_i^{\text{ext}}$.

$$\kappa_i^{(k)}(s) = 2\pi a_i \frac{D}{\lambda_{v,i}^{(k)}(s)} \frac{K_1\left(\frac{a_i}{\lambda_{v,i}^{(k)}(s)}\right)}{K_0\left(\frac{a_i}{\lambda_{v,i}^{(k)}(s)}\right)},$$

$$q_i^{(k)}(s) = \kappa_i^{(k)}(s)\left(C_{w,i}^{(k)}(s) - C_i^{\text{ext},(k)}(s)\right).$$

**(e) Repeat.** The iterative cycle repeats until the channel concentrations and external concentrations stagnate or their changes fall below a prescribed tolerance, or if a prespecified iteration count is reached. In practice, we found that any relative changes after the first 1-2 iterations are minimal. Thus, a computation with only a few iterations (ending the cycle somewhere around $k = 1 - 5$) yields a good balance of precision and efficiency. Figure S11 shows how the distributions of $C_{w,i}$, $C_i^{\text{ext}}$, $\lambda_i$, and $q_i$ shift with iteration count for a 1 million vessel tree perfusing 1 $\text{cm}^3$ of myocardium. We find that the number of iterations required scales with $\frac{\lambda_{avg}}{L_{v-v}}$ (where $\lambda_{avg} = \frac{1}{N}\sum_i^N \lambda_i$ and $L_{v-v}$ is the average vessel-to-vessel distance). We also noted that a constant $C^{\text{ext}} = 0$ is sufficient when $\frac{L_{v-v}}{\lambda_{avg}} \gtrsim 3$ to 4.

#### B.3 Tissue side: Solving for the intratissue concentration field

The concentration field $C(\vec{x})$ is evaluated pointwise within the tissue medium as a superposition of Green's functions from each nearby vessel. This evaluation is done after the final iteration $k_N$ once the vessel-vessel interactions have converged.

$$C(\vec{x}) = D^{-1} \sum_i \int q_i^{(k_N)}(s) \frac{e^{-\frac{r_i}{\lambda_{t,i}(r,s)}}}{4\pi r_i} ds,$$

The characteristic reaction-diffusion length scale used in the tissue is $\lambda_{t,i}$ and–similarly to the vessel side closure–is specific to each contributing vessel. $\lambda_{t,i}$ is computed as follows,

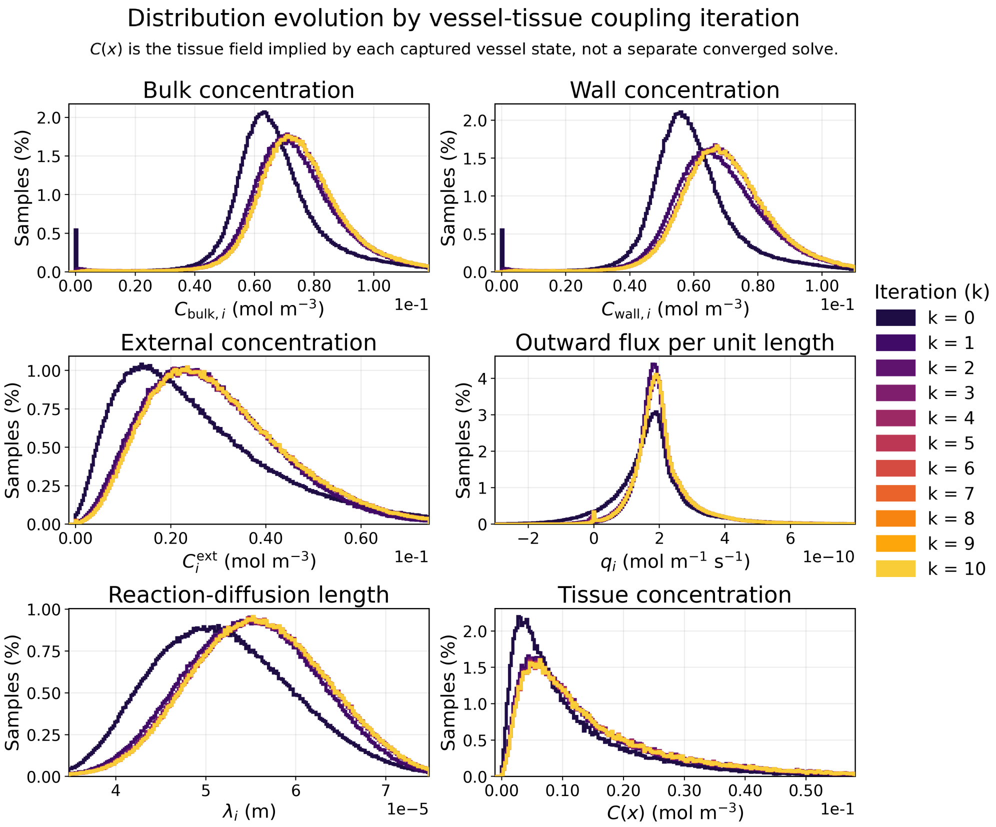

Figure S11: Representative image showing the iterative evolution of $C_{w,i}$, $C_{b,i}$, $C_i^{\mathrm{ext}}$, $\lambda_i$, $q_i$, and $C(\vec{x})$ distributions. System is mostly converged by iteration $k = 2$ and nearly entirely converged by $k = 3$. Shown sample is from a solve on a 1 million vessel tree structure.

**(a) Initialization.** For the contribution from each vessel $i$, we start with an initial guess based on solely the channel concentration $C_i^{\mathrm{eff}}(s) = C_{w,i}(s)$. We use this $C_i^{\mathrm{eff}}(s)$ within $\lambda_{t,i}(s)$.

**(b) One-shot Picard iteration using Bessel solution.** Instead of an intensive iteration scheme, we simply perform a one-shot iteration on each vessel $i$ out to the location $\vec{x}$ for the tissue point being measured. We compute this initial guess using the single-vessel modified cylindrical Bessel solution with no external field,

$$C_i^{\mathrm{eff}}(s) \approx C_{w,i}(s)\frac{K_0\left(\frac{r_i(s)}{\lambda_{t,i}(s)}\right)}{K_0\left(\frac{a_i}{\lambda_{t,i}(s)}\right)}$$

**(c) Compute final pointwise tissue concentration.** Using the one-shot Bessel guess for $C_i^{\mathrm{eff}}(s)$ to define a final $\lambda_{t,i}(s)$, we solve for $C(\vec{x})$ directly through a sum of all contributions from each vessel.

### Appendix C Reducing the pairwise problem to accelerate tissue and external concentration field calculation

In this work, we perform a Green's Function Method (GFM) using Yukawa sources (sources with embedded linear sink terms) as opposed to using Laplace sources (sources with no consumption) on the vessels and Laplace sinks distributed throughout the tissue.

Therefore, in theory, for all pairwise interactions we have reduced a problem which is typically $O(N^2 + NM + M^2)$, for $N$ sources and $M$ sinks, to an accelerated $O(N^2 + NM)$ problem, at the cost of some accuracy due to the necessary local linearizations of the Green's kernel.

Despite no longer having the $O(M^2)$ term (which at constant resolution would grow as $O(L^6)$ if L is the side length of a cubic tissue volume), we still have a problem which may become computationally expensive as the number of vessel quadrature points $N$ grows. To remedy this, we introduce a screening length, past which we no longer consider vessel-vessel or vessel-tissue interactions. Because each vessel already has a characteristic reaction-diffusion length $\lambda$, we define this window in terms of a number $N$ of these lengths such that the window length is $N\lambda$. Any Green's function contributions from vessels with distance $r > N\lambda$ from the target are excluded (whether it be a vessel for external field $C_i^{\text{ext}}$ computation or for tissue oxygen $C(\vec{x})$ computation).

To find the smallest $N$ value acceptable for reasonable results, we performed sweeps of this parameter and compared against a full pairwise solve, which includes all $N - N$ and all $N - M$ interactions. Figure S12 shows a representative case for a structure with 1 million vessel segments. In all tested cases, $N = 5$ displayed nearly identical results to the full pairwise solution while accelerating the solution greatly. In the example shown, the solution with local screening was accelerated 94-fold.

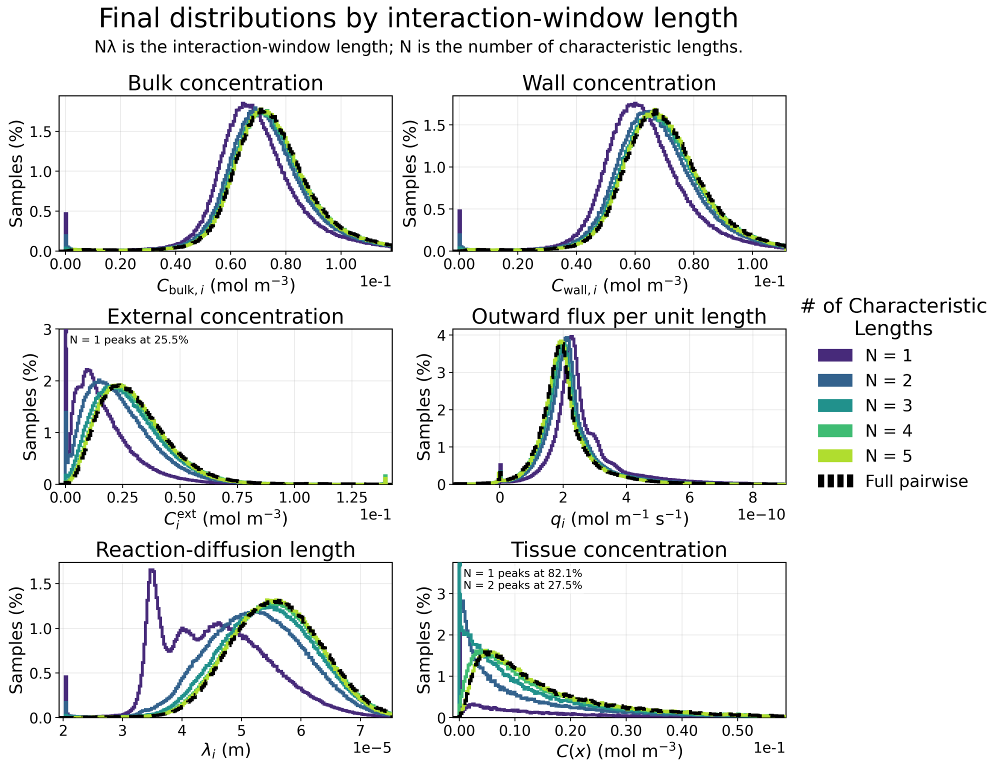

Figure S12: Representative image showing converged solutions (for $C_{w,i}$, $C_{b,i}$, $C_i^{\text{ext}}$, $\lambda_i$, $q_i$, and $C(\vec{x})$ distributions) using different $N\lambda$ interaction screening lengths. Shown sample is from a solve on a 1 million vessel tree structure truncated at $k = 5$ iterations on $C^{\text{ext}}$.

### Appendix D Rapid evaluation of the external field $C^{\text{ext}}$ using FFT on a regular grid

In the previous appendix, we showed a trick to reduce the scaling complexity of our Yukawa GFM. However, this method still requires on the order of tens of seconds to converge. If a user requires evaluation of tens of millions of vessels in single-digit seconds, the following method can be used.

To accelerate our calculations further, we decompose the external field into a smooth background contribution (as opposed to a summation of nearby pairwise interactions). The background field is evaluated on a regular Cartesian grid enclosing the vascular domain. Each vessel quadrature point is treated as a source with strength proportional to its outward flux per unit length and quadrature weight. These point source strengths are then deposited onto the grid using a compact interpolation stencil. We use a triangular-shaped-cloud (TSC) assignment as our default but offer compatibility with cloud-in-cell (CIC) assignment as well. Doing this produces a gridded source density $\rho(\vec{x})$

$$\rho(\vec{x}) \approx \sum_i \int_0^{L_i} q_i(s)\delta(\vec{x} - \vec{X}_i(s))ds$$

For a fixed reaction-diffusion length scale $\lambda$, the corresponding background field $\Phi$ satisfies the screened Poisson equation,

$$D(\nabla^2 - \lambda^{-2})\Phi(\vec{x}) = -\rho(\vec{x})$$

This equation is the smoothed, grid-based equivalent of summing the same exponentially screened Green's function contributions from all vessel source points. Instead of evaluating every source-target pair directly, we solve for the field on the grid and then interpolate the resulting background concentration back to the vessel quadrature points. In this way, the grid solve provides a fast approximation to the long-range part of $C^{\text{ext}}(s)$. In our implementation, this gridded screened Poisson problem is solved in Fourier space using a Fast Fourier Transform (FFT) method. After transforming the source density to Fourier space, the screened Poisson operator becomes diagonal, so each Fourier mode can be divided by its corresponding screened operator coefficient. An inverse FFT then returns the background concentration field on the grid.

Because $\lambda$ can vary between vessels and along the network, we group source points into a finite number of $\lambda$ bins. For each bin, we solve one screened Poisson problem, then add all resulting fields together. This captures the dominant effect of different decay lengths without requiring a separate solve for every individual source point. To remove self-node-contributions, we first smooth the self contribution using the same FFT operation, then subtract the result from the smoothed external field.

The grid field is most accurate for smooth, long-range contributions. Very nearby vessel interactions are more sensitive to the exact distance between source and target points, and these interactions can be blurred by the grid.

With high enough grid resolution and $\lambda$ bin count, this method should, in theory, approach the pairwise interaction result. Thus, we expected there to be a grid resolution and bin resolution that results in a converged solution. A sweep of grid resolutions and bin counts on various tree architectures (specifically for myocardial tissue) revealed that a grid resolution of about 128-196 $\text{cm}^{-1}$ with a bin count of 5 results in a reasonable balance of accuracy and speed. This is equivalent to a grid resolution of approximately $1\lambda_{\text{avg}}$ ($\pm 20\%$). Further testing found this to be a reasonable benchmark for resolution selection, while 5-6 bins remained a functional choice. Shown in Figure S13 is a sweep of grid resolutions for a representative vessel architecture using 5 bins. We see in this example that a resolution near or past $1\lambda_{\text{avg}}$ is sufficient for matching the full pairwise solution. However, we should note that the long tails of these distributions are somewhat poor, and very high $C_i^{\text{ext}}$ values are generally underestimated by the FFT regular grid approximation. Figure S14 shows a crosscut of the xy-plane (z=0), depicting the additive contribution of each of these five $\lambda$ bins.

In the example case shown, the FFT regular grid is about 19 times faster. To trade for this speed, the accelerated method increases prediction error, increasing MAE by  1.1% and RMSE by  1.9%.

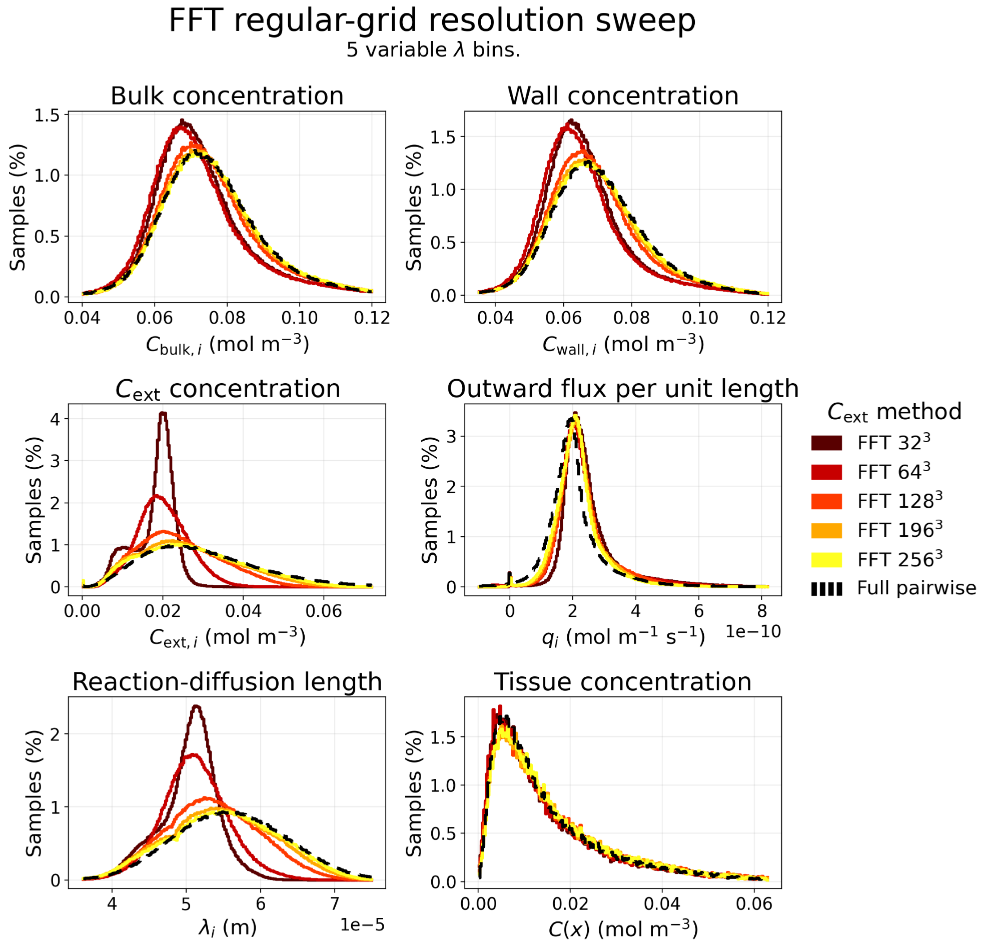

Figure S13: Representative image showing converged solutions (for $C_{w,i}$, $C_{b,i}$, $C_i^{\text{ext}}$, $\lambda_i$, $q_i$, and $C(\vec{x})$ distributions) using different FFT regular grid resolutions within a 1 cm$^3$ cube of myocardial tissue. Shown sample is from a solve on a 1 million vessel tree structure truncated at $k = 5$ iterations on $C^{\text{ext}}$.

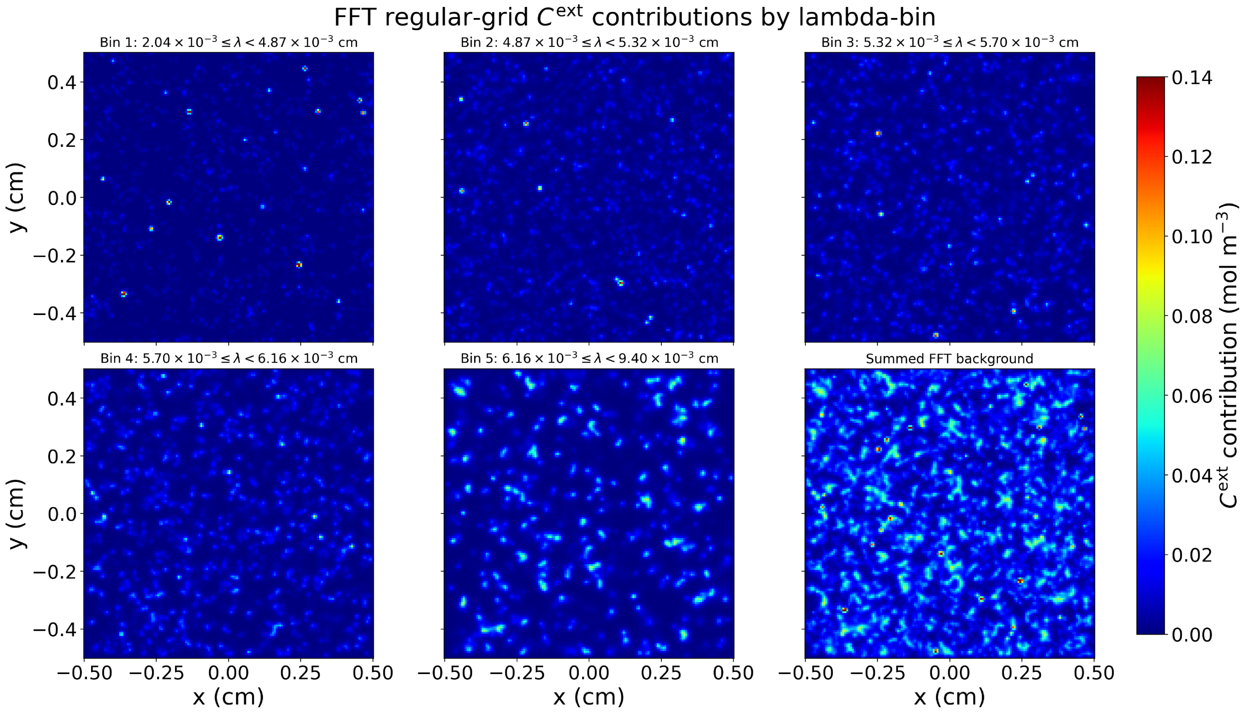

Figure S14: External concentration field contributions from each lambda bin along the z=0 plane. Shown is a 1 million vessel tree structure, $k = 5$ iterations, $256^3$ grid resolution.

#### Appendix E Graetz eigenfunction resolution assessment for intraluminal diffusion

As with the previous section, each discretized solution of a continuous function must be carefully assessed to ensure a reasonable balance of speed and self-consistency. Here we look to quantify the ideal settings for our discretized eigenfunction expansion solver for the concentration profiles within the vessel lumens. Figure S15 shows a sweep of $n_r$ number of discrete radial profile points and $n_m$ number of retained eigenmodes. Higher $n_r$ means a smoother profile to which the eigenmodes are fit, and higher $n_m$ means more eigenmodes can be used for the expansion. $n_r = 6$ and $n_m = 2$ appear to be sufficient. Generally, $n_m \geq 2$ yields diminishing returns. However, $n_r = 8$ and $n_m = 3$ or 4 should be used if very high accuracy is desired for $C_b$ and $C_w$ computation.

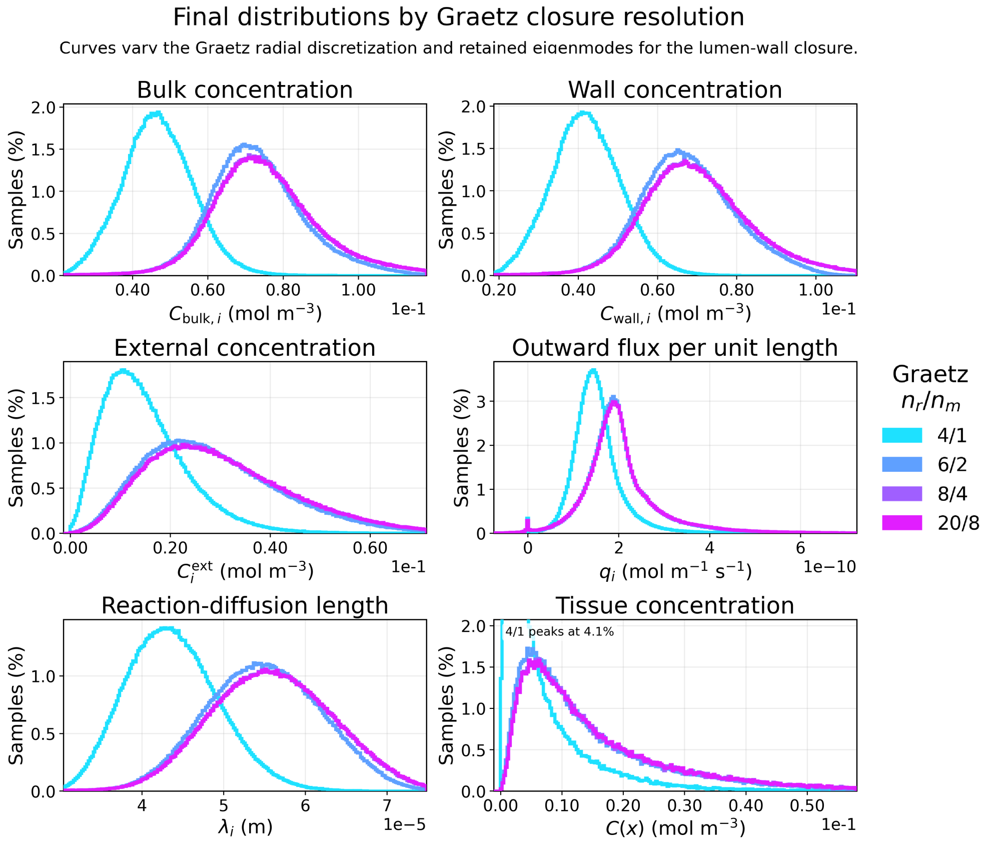

Figure S15: Representative image showing converged solutions (for $C_{w,i}$, $C_{b,i}$, $C_i^{\mathrm{ext}}$, $\lambda_i$, $q_i$, and $C(\vec{x})$ distributions) with increasing number of discrete radial profile points $n_r$ and number of retained eigenmodes $n_m$. The system is mostly consistent with settings of 6/2. Shown sample is from a solve on a 1 million vessel tree structure.

#### Appendix F Impact of neglecting axial diffusion $D\frac{\partial^2 c}{\partial s^2}$ in the diffusion operator $D\nabla^2 c$.

##### F.1 Theory for including the axial diffusion term

Even with the axisymmetric disturbance field assumption $\frac{1}{r^2}\frac{\partial^2 c}{\partial \theta^2} \approx 0$, there still remains the axial variation term $\frac{\partial^2 c}{\partial s^2}$ to consider. For the disturbance field $c$, any axial variation of tissue concentration is driven by the vessel-wall mismatch

$C_w(s) - C^{\text{ext}}(s)$. With axial diffusion included, the axisymmetric disturbance equation becomes

$$D\left(\frac{\partial^2 c}{\partial r^2} + \frac{1}{r}\frac{\partial c}{\partial r} + \frac{\partial^2 c}{\partial s^2}\right) - D\lambda^{-2}c = 0$$

with boundary conditions

$$c(a,s) = f(s) = C_w(s) - C^{\text{ext}}(s)$$

and

$$c(r\to\infty, s) \to 0$$

To solve this problem, we must decompose the axial wall mismatch into Fourier modes,

$$f(s) = \frac{1}{2\pi}\int_{-\infty}^{\infty} \hat{f}(k)e^{iks}dk$$

Because the governing equation is linear with constant $\lambda$, each Fourier mode evolves independently. This gives...

$$c(r,s) = \frac{1}{2\pi}\int_{-\infty}^{\infty} \hat{c}(r)e^{iks}dk$$

such that each axial mode $k$ satisfies

$$\frac{\partial^2 \hat{c}_k}{\partial r^2} + \frac{1}{r}\frac{\partial \hat{c}_k}{\partial r} - (k^2 + \lambda^{-2})\hat{c}_k = 0$$

Defining

$$\alpha_k = \sqrt{k^2 + \lambda^{-2}}$$

the decaying solution is the modified Bessel function $K_0$. Applying the wall condition $\hat{c}_k(a) = \hat{f}(k)$ the mode's solution is

$$\hat{c}_k(r) = \hat{f}(k)\frac{K_0(\alpha_k r)}{K_0(\alpha_k a)}$$

Therefore the full axial-diffusion-corrected disturbance field is

$$c(r,s) = \frac{1}{2\pi}\int_{-\infty}^{\infty} \hat{f}(k)\frac{K_0(\alpha_k r)}{K_0(\alpha_k a)}e^{iks}dk$$

The corresponding outward wall flux per unit vessel length is

$$\hat{q}_{r,s}(k) = -2\pi a D\frac{\partial \hat{c}_k}{\partial r}\bigg|_{r=a} = 2\pi a D\alpha_k\frac{K_1(\alpha_k a)}{K_0(\alpha_k a)}\hat{f}(k)$$

$$q_{r,s}(s) = \frac{1}{2\pi}\int_{-\infty}^{\infty} \hat{q}_{r,s}(k)e^{iks}dk$$

Note that the radial-only result is recovered from the zero-wavenumber mode. For $k = 0$,

$$\alpha_0 = \lambda^{-1}$$

The radial-only Bessel closure corresponds to the leading axial mode, i.e. the limit in which $f(s)$ is locally uniform along the vessel. The importance of axial diffusion is controlled by the dimensionless quantity $k\lambda$. For slowly varying axial profiles ($k\lambda \ll 1$), we have

$$\alpha_k = \lambda^{-1}\sqrt{1 + k^2\lambda^2} \approx \lambda^{-1}\left(1 + \frac{(k\lambda)^2}{2}\right)$$

In this limit, the mass transfer constant collapses to the radial-only solution $\kappa(k) \approx \kappa_r$. If the characteristic axial variation length is $L_s$, then we may approximate

$$k \sim \frac{1}{L_s}$$

and the relevant parameter is $\frac{\lambda}{L_s}$. In the radial-only solution, $L_s = \frac{\mathcal{B}Q}{\kappa_r}$.

#### F.2 Numerical comparison with the axial+radial BVP

In the numerical comparisons shown in Figure S16, we compare the wall flux and intravascular oxygen profiles predicted by the radial-only approximation and the full axial+radial boundary value problem (BVP) solved in MATLAB. For the radial-only model, the wall flux is given locally by $q_r(s) = \kappa_r(C_w(s) - C^{\text{ext}}(s))$ where $\kappa_r$ is the radial-only mass transfer coefficient.

When axial diffusion is retained, the wall flux is no longer determined by a purely local relation because axial transport couples neighboring axial locations along the channel. In the full model, $\lambda$, $C_w(s)$, and $q(s)$ are coupled through the governing equations and evolve self-consistently along the vessel.

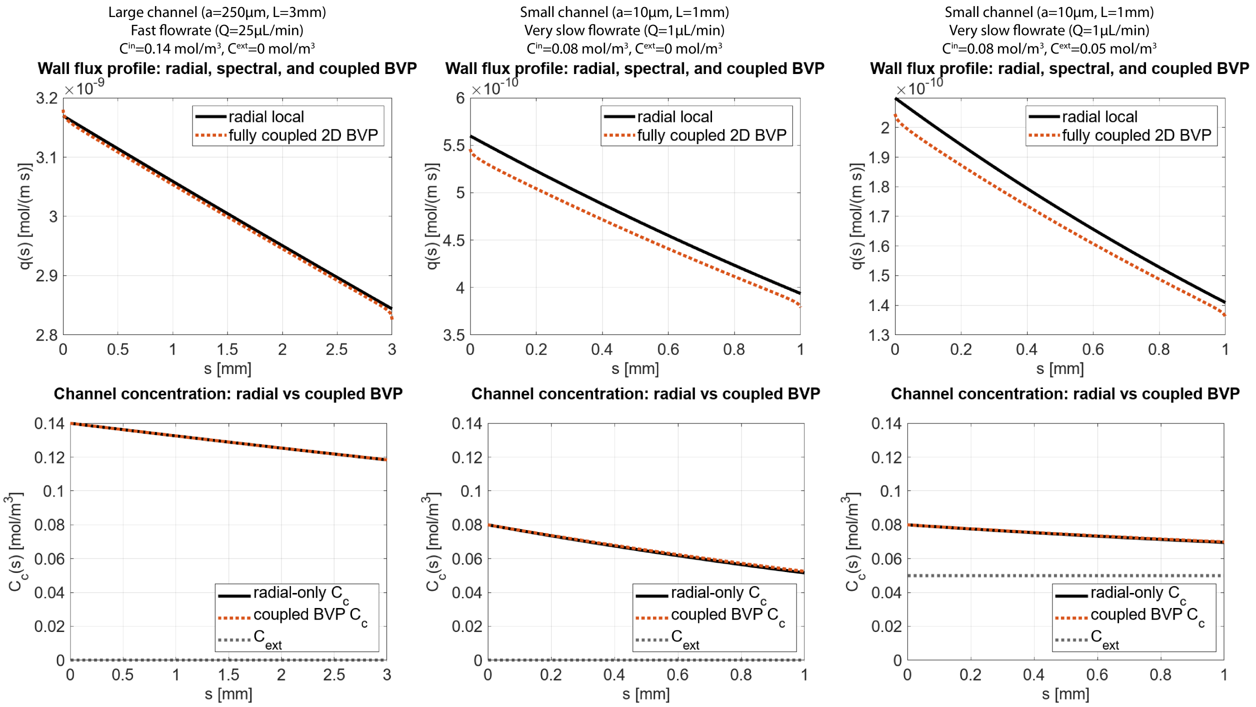

Figure S16: Comparison between the full boundary value problem (with axial and radial diffusion) and the local, radial approximation used in our derivations. Flux per unit length (top) and intravessel concentration (bottom) comparisons for three representative cases of vessels flowing through myocardium.

### Appendix G Scaling arguments on $O(a^2)$ terms

#### G.1 Scaling setup and definitions

We now estimate the relative size of the finite-radius corrections compared with the leading-order line-source monopole term

$$C(\vec{x}) = \sum_i \int_0^{L_i} q_i(s) G(\vec{x}, \vec{X}_i(s)) ds$$

We will do so for the constant $\lambda_i$ case.

It is convenient to define the geometric curvature factor

$$\mathcal{K}_i = \frac{p_i : \nabla\nabla G_i}{G_i} = (1 - 3(\vec{t}_i \cdot \hat{r}_i)^2)\left(\frac{1}{r_i^2} + \frac{1}{\lambda_i r_i}\right) + \frac{1 - (\vec{t}_i \cdot \hat{r}_i)^2}{\lambda_i^2}$$

Equivalently, with the nondimensional distance $\rho_i = \frac{r_i}{\lambda_i}$ we may write

$$\mathcal{K}_i = \frac{1}{\lambda_i^2}\mathcal{F}_i(\rho_i, (\vec{t}_i \cdot \hat{r}_i)^2)$$

Where

$$\mathcal{F}_i(\rho_i, (\vec{t}_i \cdot \hat{r}_i)^2) = (1 - 3(\vec{t}_i \cdot \hat{r}_i)^2)\left(\frac{1}{\rho_i^2} + \frac{1}{\rho_i}\right) + (1 - (\vec{t}_i \cdot \hat{r}_i)^2)$$

The factor $\mathcal{F}$ is order one when $r_i \sim \lambda_i$, but it can become much larger for evaluation points closer to the vessel centerline.

We also use the wall flux relation $q_i(s) = \kappa_i(s)(C_{w,i}(s) - C_i^{\text{ext}}(s))$ with $\kappa_i = \frac{2\pi a_i D}{\lambda_i}\frac{K_1(\frac{a_i}{\lambda_i})}{K_0(\frac{a_i}{\lambda_i})}$.

Let $\Delta C_i = C_{w,i} - C_i^{\text{ext}}$ and $\epsilon_i = \frac{a_i}{\lambda_i}$

The magnitudes of each term in SI units are...

- $a \sim 10^{-6} - 10^{-4}$, mostly $10^{-6}$
- $C_w \sim 10^{-5} - 10^{-1}$, mostly $10^{-2}$
- $C^{\text{ext}} \sim 10^{-5} - 10^{-1}$, mostly $10^{-2}$ or $10^{-3}$ and almost always $< C_w$
- $\Delta C_i$ typically not lower than $10^{-4}$, often larger (usually $10^{-3}$)
- $V_{\max} \sim 10^{-3} - 10^{-2}$
- $K_M \sim 10^{-3} - 10^{-2}$
- $\lambda \sim 10^{-5} - 10^{-4}$, mostly $10^{-4}$
- $\frac{K_0}{K_1} \sim 10^{-2} - 10^{0}$, mostly $10^{-2}$
- $r$ evaluated numerically from $a$ to $\sim 4\lambda$ such that $\left(\frac{1}{r} + \frac{1}{\lambda}\right)^2 \sim 10^7 - 10^{12}$ and in the range of $10^8 - 10^9$ for most evaluations where $r \sim \lambda$
- $|\mathcal{F}| \sim 10^0$ where $r \sim \lambda$
- $|\mathcal{K}| \sim \frac{|\mathcal{F}|}{\lambda^2} \sim 10^8$ on most occasions

#### G.2 Scaling of second-order monopole term $C^{(2)}_{\text{monopole}}/C^{(0)}_{\text{monopole}}$

The second-order monopole correction is

$$C^{(2)}_{\text{monopole}} = \sum_i \int_0^{L_i} \frac{a_i^2 q_i(s)}{4} p_i : \nabla\nabla G_i ds$$

Over one local vessel contribution, the factors $q_i(s)$, $G_i$, and local integration length cancel in the ratio, giving

$$\frac{C^{(2)}_{\text{monopole}}}{C^{(0)}_{\text{monopole}}} \sim \frac{1}{4}\left(\frac{a}{\lambda}\right)^2 \mathcal{F}$$

In the typical evaluation case where $r \sim \lambda$ and vessels are capillary-scale,

$$\left|\frac{C^{(2)}_{\text{monopole}}}{C^{(0)}_{\text{monopole}}}\right| \sim 10^{-5} - 10^{-4}$$

although larger contributions are possible.

##### G.3 Scaling of second-order dipole term $C^{(2)}_{\text{dipole}}/C^{(0)}_{\text{monopole}}$

The second-order dipole correction is

$$C^{(2)}_{\text{dipole}} = \pi D \sum_i a_i^2 \int_0^{L_i} C_{w,i}(s) p_i : \nabla\nabla G_i ds$$

therefore a local contribution (using $q = \kappa \Delta C$) has a relative magnitude of

$$\frac{C^{(2)}_{\text{dipole}}}{C^{(0)}_{\text{monopole}}} \sim \frac{\pi D C_w}{q} \left(\frac{a}{\lambda}\right)^2 \mathcal{F} \sim \frac{1}{2} \frac{a}{\lambda} \frac{C_w}{\Delta C} \frac{K_0(\frac{a}{\lambda})}{K_1(\frac{a}{\lambda})} \mathcal{F}$$

Using the typical values for each parameter,

$$\left| \frac{C^{(2)}_{\text{dipole}}}{C^{(0)}_{\text{monopole}}} \right| \sim 10^{-3} - 10^{-2}$$

Thus, the dipole finite-radius correction is usually still small but can be noticeable.

#### Appendix H Murray's Law derivation

Murray's Law can be understood as a minimum-work principle for living vascular networks. Driving flow through a cylindrical vessel requires mechanical pumping power $\dot{W}_{\text{flow}} = Q\Delta P$. With Poiseuille flow, pressure drop scales as $\Delta P \sim \mu L Q / r^4$. Murray proposed that a living vessel also has a maintenance cost proportional to its volume.

$$V = \pi r^2 L \;\rightarrow\; \text{Cost} \propto r^2 L \;\therefore\; \dot{W}_{\text{maintenance}} = \; k r^2 L$$

Therefore,

$$\dot{W}_{\text{total}} = \dot{W}_{\text{flow}} + \dot{W}_{\text{maintenance}} = \frac{8\mu L Q^2}{\pi} \frac{1}{r^4} + kLr^2 = \frac{\alpha}{r^4} + \beta r^2$$

Taking the derivative,

$$\frac{d\dot{W}_{\text{total}}}{dr} = 0 = -\frac{4\alpha}{r^5} + 2\beta r$$

we may find the ideal flow rate (Q) scaling

$$r^6 = \frac{2\alpha}{\beta} = \frac{16\mu Q^2}{k\pi} \;\rightarrow\; Q_{\text{ideal}} \propto r^3$$

By extension, if at a bifurcation

$$Q_{\text{in}} = \sum_{\text{out}} Q_i$$

then the ideal radius scaling would be

$$r_{\text{in}}^3 = \sum_{\text{out}} r_i^3$$

This is Murray's Law for vascular bifurcations.

#### Appendix I Net resistance scaling

For a single cylindrical vessel segment, Hagen-Poiseuille flow suggests the resistance (R) scaling

$$R_i \propto \frac{L_i}{r_i^4}$$

We will assume that bifurcation statistics tend toward a roughly symmetric binary tree. Murray's Law on an equally bifurcating tree suggests

$$r_k = r_0 2^{-\frac{k}{3}}$$

where $k$ is the generation number and $r_0$ is the root segment radius. We see that the average vessel length scales as $\overline{L} \propto N^{-\alpha}$, so the length of each generation's vessels may be expressed as $L_k = L_0 2^{-\alpha k}$ where $L_0$ is the root segment length.

With these scalings, the single-segment resistance can be expressed as

$$R_k \sim \frac{L_k}{r_k^4} \sim \frac{2^{-\alpha k}}{2^{-\frac{4k}{3}}} = 2^{\left(\frac{4}{3}-\alpha\right)k}$$

The total equivalent resistance for generation $k$ would then be

$$R_k^{eq} \sim \frac{R_k}{2^k} = 2^{\left(\frac{1}{3}-\alpha\right)k}$$

and the net resistance

$$R_{net} \sim \sum_{k=0}^{k_{\max}} R_k^{eq}$$

Furthermore, the total length would be

$$L_{tot} \sim \sum_{k=0}^{\mathrm{k_{max}}} 2^k L_k \sim \sum_{k=0}^{\mathrm{k_{max}}} 2^{(1-\alpha)k} \sim 2^{(1-\alpha)k_{\max}} \sim N^{1-\alpha}$$

assuming that the final generation $k$ contributes the most to the total length of the tree. Therefore, the resistance per unit length scaling depends greatly on the value of $\alpha$. We may perform another scaling analysis to find $\alpha$.

We are building these vascular trees in a bounded tissue domain where the mean spacing between the terminal channels scales as $d_{\mathrm{t}} \sim \left(\frac{V}{N}\right)^{\frac{1}{3}} \sim N^{-\frac{1}{3}}$. New terminals add branches to the nearest existing vessel, giving a new connection distance (new vessel length addition) of $\Delta L \sim d_{\mathrm{t}}$. Therefore, the marginal length increment obeys

$$\frac{dL_{\mathrm{tot}}}{dN} \sim N^{-\frac{1}{3}}$$

Integrating gives

$$L_{\mathrm{tot}} \sim \int_1^N n^{-\frac{1}{3}} dn \sim N^{\frac{2}{3}}$$

Since $L_{\mathrm{tot}} \sim N^{1-\alpha}$, this gives us a theoretical value of $\alpha \approx \frac{1}{3}$. In our CCO-generated trees, we measure this exponent to be about 0.64 (and therefore $\alpha$ =0.36), very close to our predicted value.

Using 1/3 (and ignoring any slowly-varying logarithmic factors), net resistance per unit length scales as

$$\frac{R_{\mathrm{net}}}{L_{\mathrm{tot}}} \sim \frac{1}{N^{1-\alpha}} = N^{-\frac{2}{3}} \;\rightarrow\; \log_{10}\left(\frac{R_{\mathrm{net}}}{L_{\mathrm{tot}}}\right) \sim -\frac{2}{3}\log_{10}N$$

This is very close to the scaling we measure for CCO-generated trees, which is

$$\log_{10}\left(\frac{R_{\mathrm{net}}}{L_{\mathrm{tot}}}\right) \sim -0.71\log_{10}N$$

It is unsurprising that a bifurcation-optimizing method like CCO somewhat outperforms binary-tree predictions.

### Appendix J Krogh cylinder model derivation and comparison to FEM/Green's

#### J.1 Introduction to the Krogh cylinder model

The Krogh cylinder model is a classical analytical framework for estimating oxygen transport from capillaries into surrounding tissues [17]. In this model, each vessel is idealized as a straight cylindrical oxygen source surrounded by a concentric cylindrical tissue domain (a region of influence), with the outer radius representing the effective territory supplied by that vessel. Oxygen is assumed to only diffuse radially and be consumed within the tissue volume via a 0th order reaction rate.

#### J.2 Krogh cylinder model derivation

The 0th order reaction rate, radial-diffusion-only mass balance equation in the tissue is...

$$\frac{D}{r}\frac{d}{dr}\left(r\frac{dC}{dr}\right) = k$$

At the vessel boundary $r = a$, concentration is matched $C(a,s) = C_w(s)$. At the Krogh radius (the midpoint between channels) $r = R$, a zero-flux condition is prescribed $\frac{dC}{dr}\big|_{r=R} = 0$.

Solving this gives

$$C(r,s) = C_w(s) + \frac{k}{4D}\left(r^2 - a^2 - 2R^2 \ln\left(\frac{r}{a}\right)\right)$$

The axial depletion is equivalent to the flux delivered per unit length for a zeroth-order reaction

$$Q\frac{dC}{ds} = -q(s) = -2\pi a J(s) = 2\pi a D\frac{dC}{dr}\Big|_{r=a} = -\pi k\left(R^2 - a^2\right)$$

Figure S17 shows a schematic of the Krogh model setup. Below is a shortlist of assumptions made by this model:

- Each vessel may be represented by a straight cylinder with constant-radius, circular cross-section.
- The tissue supplied by a vessel may be represented as a concentric cylindrical shell with constant outer radius $R$. No oxygen crosses this boundary.
- Neighboring vessels are replaced by a symmetry boundary, and this is the extent to which vessels interact with each other. There is no vessel-vessel external field interaction, oxygen re-uptake, or other phenomena regarding more than one vessel.
- The tissue domain is axisymmetric around the vessel.
- Radial diffusion dominates.
- Diffusivity is constant in the tissue.
- No vessel wall resistance.
- Fast radial diffusion inside the channels (uniform channel concentration).
- Tissue oxygen consumption is zeroth order; therefore, the reaction rate is spatially uniform having no dependence on local concentration (cellular metabolism is the same everywhere).
- All vessels are parallel with each other.

#### J.3 Comparison to Green's for 1 channel or 2 parallel channels

We tested the Krogh model against our Green's function method in two simple geometries where its assumptions should be most favorable: a single straight channel and two parallel, equal-size, equal-region-of-influence straight channels. For each case, we compare the Krogh-predicted tissue oxygen field against a Green's function solution using the same vessel geometry, flow rate, inlet concentration, and diffusivity.

The Krogh radius was defined using the circumscribed Voronoi distance of nearest-neighbor Voronoi cells (see subsection J.4).

The $0^{\text{th}}$ order reaction rate was treated as a fitted parameter. We varied $0 \le k \le V_{\max}$ to give the best agreement with the Green's solution according to the CCC metric.

Figure S18 shows these two tests. Results for a single channel are on the left, dual channel on the right. For each geometry, the top left and top right plots show a cross-section of a cellular medium representing this work's experimental setup (10% (w/v) GelMA, 60M HEK cells/mL). In each plot, a thin white line represents the contour past which we would expect cells to be oxygen-limited and perhaps necrose long-term ($C < C^*$). The bottom left plot shows the difference between the two cross-sections. The bottom right shows a parity plot (of point $O_2$ measurements) of the two methods on a linear scale.

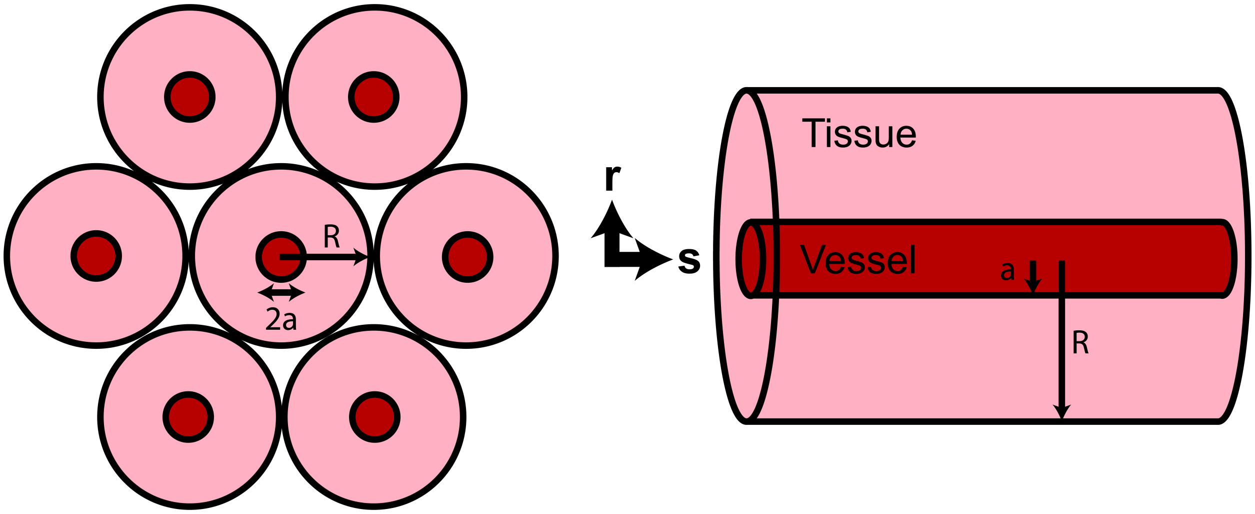

Figure S17: Schematic of the Krogh cylinder model seen from along the flow direction (left) and from side view (right).

Note that fitting $k$ by mean absolute error (MAE) or root mean squared error (RMSE) produced visually identical results.

From this comparison, in both geometries, the Krogh model captured the qualitative oxygenated region around the channels, but produced a more compact, sharper oxygen profile than the Green's model, causing the Krogh model to overpredict oxygen close to the vessel and underpredict it farther away. This was found to be so even with remarkably low fitted oxygen consumption rates of $k \approx 0.015V_{\max}$ (single channel) and $k \approx 0.02V_{\max}$ (dual channel).

For the live/dead contours to have approximately the same placement as in the Green's method, this $k/V_{\max}$ ratio must be about 0.011 and 0.016 respectively. With these consumption rates, the oxygen values within the contour (closer to the channels) are unanimously higher than the Green's method predictions; in some places the Krogh model predicts oxygen concentrations up to 34x the Green's prediction.

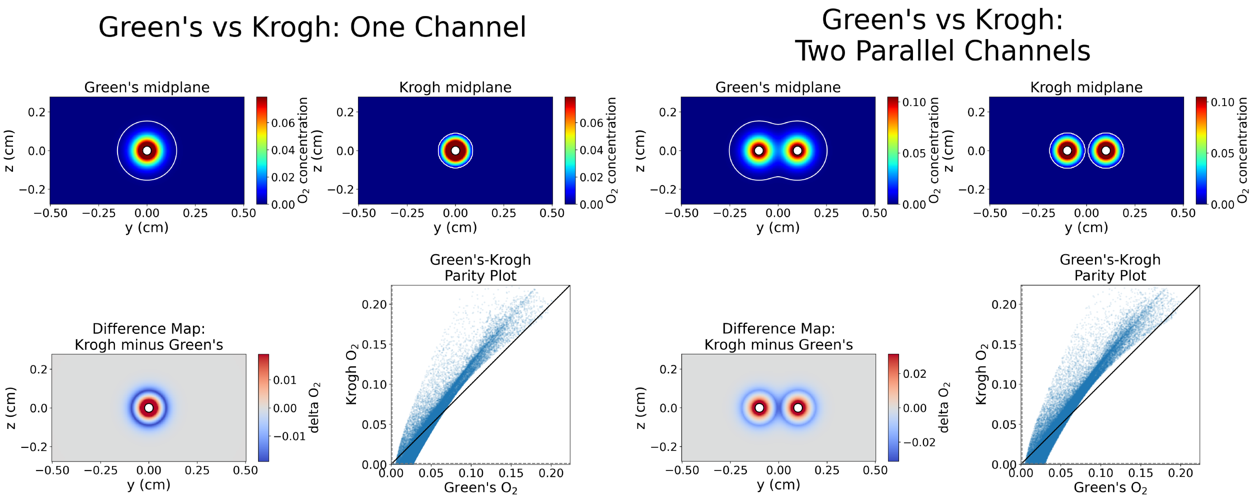

Figure S18: One channel and dual channel comparisons between the Krogh cylinder model and this work's Green's function model. Plots show cross-sectional oxygen fields, field deltas, and point-measurement parity plots.

#### J.4 Assignment of Krogh cylinder radii to non-parallel vessels and comparison to FEM for complex vasculature architectures

For biomimetic networks with non-parallel vessels supplying nonuniform volumes of tissue, a single geometric spacing $2R$ between neighboring vessels is not well defined. We therefore assigned each tissue point to its nearest vessel segment and tested several ways of converting the resulting vessel territories into effective Krogh cylinder radii.

We tried eight methods of assigning vessels Krogh radii and compared them to the same 25 FEM case studies used in Section 8:

- **Same volume**: the same tissue volume as the vessel's Voronoi-assigned region.
- **Same surface area**: the same surface area as the vessel's Voronoi-assigned region.
- **Classical spacing**: half of the nearest non-incident vessel spacing.
- **Global vessel length density**: one global radius from the total tissue volume divided by total vessel length.
- **Lattice area equivalent**: area-equivalent radius based on hypothetical square lattice spacing of vessels.
- **Inscribed Voronoi**: conservative inner radius selection from the sampled Voronoi region.
- **Circumscribed Voronoi**: the maximum radial extent of the sampled Voronoi-assigned tissue region.
- **Moment-matched**: radius chosen to match the second radial moment of assigned tissue points $R = \sqrt{2\langle r^2\rangle_{\mathbf{Vor}} - a^2}$.

Of these, all methods performed similarly with respect to CCC and live/dead classification accuracy, except for ”classical spacing” and ”lattice area equivalent” which performed very poorly in both metrics. The best performer overall was the ”circumscribed Voronoi” method, which exhibited a mean CCC of 0.72, pooled dead-class accuracy of 98.6%, pooled live-class accuracy of 40.8%, and overall live/dead accuracy of 70.6%. As with the previous section, these results show that the Krogh model consistently overestimated the amount of tissue that would be dead or nonfunctional (or, if $k$ was optimized for live/dead classification, would greatly overpredict oxygen distributions).

Similarly to the previously shown parity plots/confusion matrices (Figure S10) for the Green's method, we also report Figure S19 which shows the same analyses for the Krogh model. In this figure, we show the Krogh model with Krogh radii assigned by the ”circumscribed Voronoi” method and $0^{\text{th}}$ order reaction rate consumption $k$ optimized for highest CCC with FEM results.

We report all error metrics for the Krogh model with $k/V_{\max}$ optimized for high CCC (Table S4) and for high live/dead classification accuracy (Table S5). Note that these metrics are for per-case optimizations, meaning that this is the ”best” performance that could be achieved by the Krogh model in this setting. In reality, one would need to estimate $k$ from experiments, reducing the model's accuracy further.

| Error metric | All evaluations | Low-C measurements |
|---|---|---|
| Total number of point evaluations ($N_{\text{evaluations}}$) | 233300 | 144520 |
| Concordance correlation coefficient (CCC) | 0.8000 | 0.0551 |
| Root mean square error (RMSE) | 0.0207 | 0.0062 |
| Mean absolute error (MAE) | 0.0106 | 0.0017 |
| Coefficient of determination ($R^2 = 1 - SSE/SST$) | 0.5484 | -16.77 |
| Squared Pearson correlation ($\rho^2$) | 0.6502 | 0.0152 |
| Mean signed error / bias | -0.00181 | 0.00052 |
| Live/dead prediction accuracy | 70.6% | N/a |
| False-alive prediction rate | 1.4% | N/a |
| False-dead prediction rate | 59.2% | N/a |

Table S4: Error metrics reported for all oxygen point evaluations across 25 case studies comparing the Krogh cylinder model method to FEM with $k$ optimized directly on the data for CCC. RMSE, MAE, and bias are reported in $mol/m^3$.

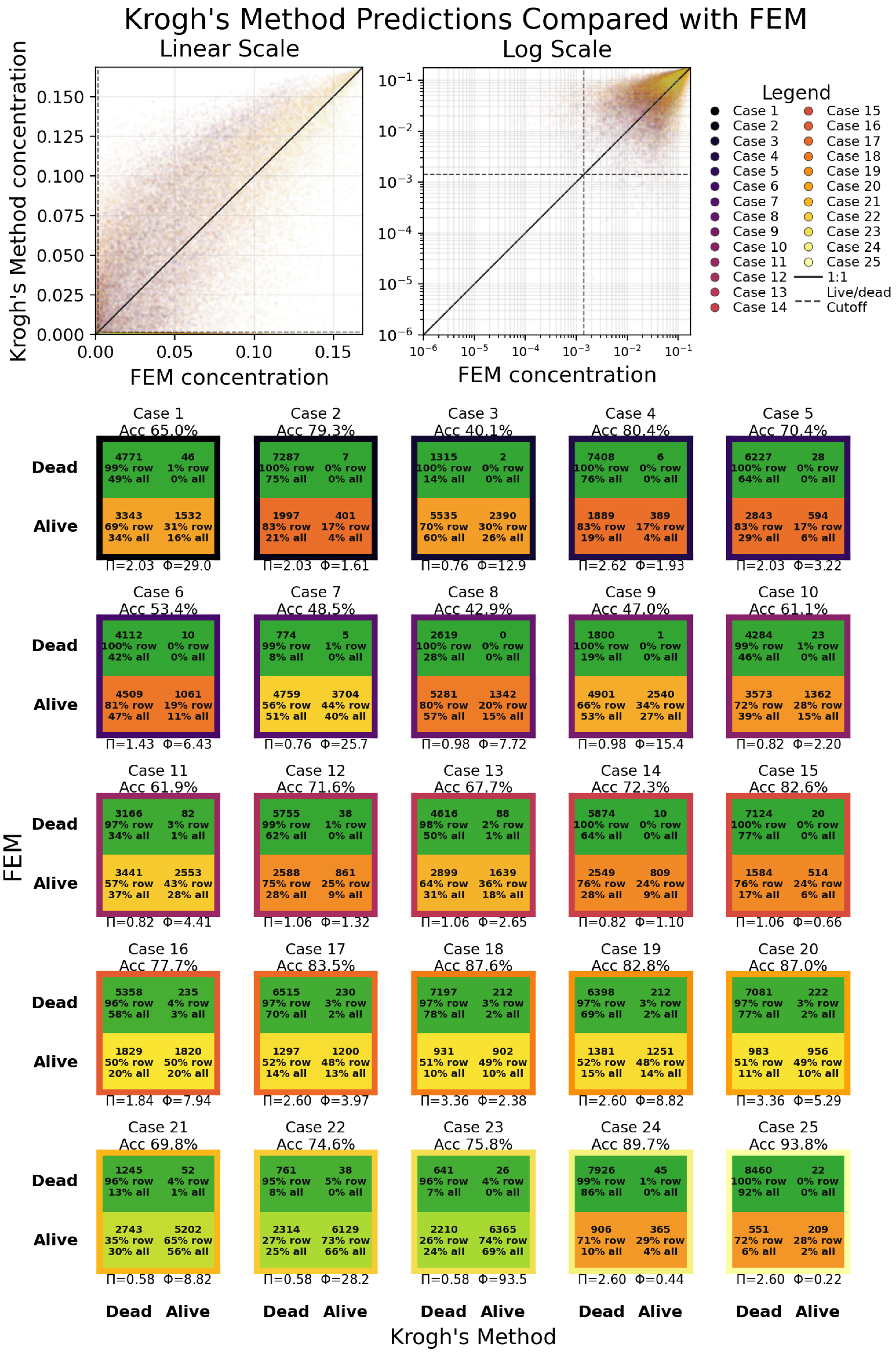

Figure S19: 1-to-1 parity plots (top) showing oxygen point samples for the same 25 steady-state simulation case studies performed with FEM and the Krogh cylinder method. Two scales are shown: a linear scale (left) and log-base-10 scale (right). Scatter-points signify oxygen measurements at one (x,y,z) location in the perfused tissue domain. Each case study is paired with a confusion matrix (bottom) showing live/dead classification accuracy.

| Error metric | All evaluations | Low-C measurements |
|---|---|---|
| Total number of point evaluations ($N_{evaluations}$) | 233300 | 144520 |
| Concordance correlation coefficient (CCC) | 0.5333 | 0.0436 |
| Root mean square error (RMSE) | 0.0454 | 0.0284 |
| Mean absolute error (MAE) | 0.0280 | 0.0119 |
| Coefficient of determination ($R^2 = 1 - SSE/SST$) | -1.167 | -368.3 |
| Squared Pearson correlation ($\rho^2$) | 0.5132 | 0.2174 |
| Mean signed error / bias | -0.0250 | 0.0113 |
| Live/dead prediction accuracy | 84.1% | N/a |
| False-alive prediction rate | 13.8% | N/a |
| False-dead prediction rate | 18.1% | N/a |

Table S5: Error metrics reported for all oxygen point evaluations across 25 case studies comparing the Krogh cylinder model method to FEM with $k$ optimized directly on the data for live/dead classification accuracy. RMSE, MAE, and bias are reported in $mol/m^3$.

### Appendix K Sensitivity of $V_{\max}$ multiplier and $C^*$ choice in simulations of GelMA constructs

In the main text, we multiplied $V_{\max}$ by 0.7 and prescribed a critical threshold of 1.4 mmol/m$^3$ (about 1 mmHg). Figure S20 shows a sensitivity analysis of how the simulated surviving diameter around the channel (for the single, axial channel) changes with respect to these two choices.

We found that varying the multiplier instead from 0.5 to 1 resulted in a viable diameter change of +17% to -16% (with constant $C^*$). Varying $C^*$ from 0.1× to 5× the chosen value changed viable diameter by +7% to -27%. Two-parameter variation yielded extremes of +27% and -37%.

In the GelMA construct experiments, the experiment-averaged single-channel viable ring diameter dropped from 90% viability to 10% viability over a distance equivalent to 45% of the mean diameter at 50% viability.

$$\frac{|d_{90\%} - d_{10\%}|}{d_{50\%}} \approx 0.45$$

We should note that the YGFM theory predicts the viability stepwise cutoff to be around where the experiments reached 70-80% viability (on average), so the simulation accuracy is more sensitive to higher $C^*$ and metabolic multiplier.

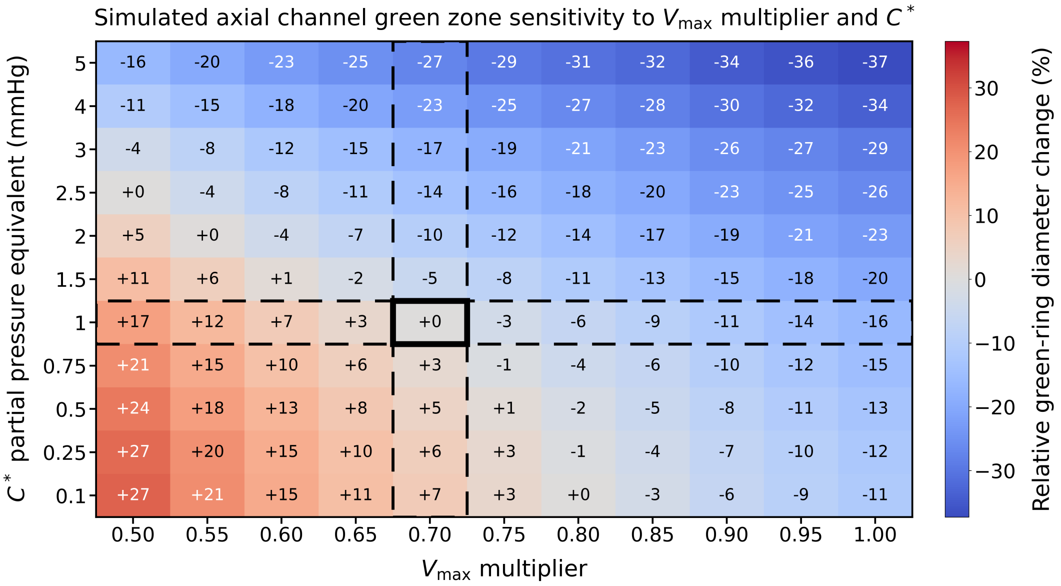

Figure S20: Sensitivity of simulated axial-channel viable-zone diameter to metabolic scaling and oxygen-limited threshold. The heatmap shows the relative change in green-zone diameter surrounding the axial channel as a function of applied $V_{max}$ multiplier and $C^*$. Values are reported relative to the chosen values of 0.7 (multiplier) and 1 mmHg (cutoff), outlined in solid black. The dashed row/column indicate sensitivity with only one parameter being varied.

### Appendix L  Additional methodological details

- **Computer specifications:** All computations performed in this study, including those shown in the supplemental information here, were performed on a Dell Precision 7560 workstation. This computer is equipped with an 11th Gen Intel Core i9-11950H @ 2.60 GHz processor and an NVIDIA GeForce RTX 3080 Laptop GPU (64.0 GB of RAM and 16 GB GPU memory in total).
- **Image editing:** Live/dead images shown in the main text were refined for presentation purposes. The two live/dead images in the second figure received a slight boost in red/green exposure for presentation purposes; all layers in the confocal .tiff image stack were superimposed to show the full thickness of the imaged section in one figure. We also cropped the displayed images to the construct region. All quantitative analyses were performed on raw live/dead image stacks without any of these edits present.
- **Evaluation of line integrals:** Gauss-Legendre quadrature was used to evaluate all line integrals in this study. Unless otherwise stated, we used five-point Gauss-Legendre quadrature.
- **Why a serpentine channel instead of a bi/trifurcating channel?** A serpentine channel was chosen over a bifurcating or trifurcating design to limit (a) the trapping of and failure to purge air bubbles from collateral channels and (b) rotation about a confinement axis, both of which we observed in early experiments.
- **How long until oxygen equilibrated in the HEK-293T cellular constructs?** Although oxygen-measurement experiments perfused the constructs for 24 hours, the spatial oxygen measurements visually appeared to reach equilibrium within about 1-2 hours of beginning perfusion.
